# Repertoire-level generation of T-cell epitopes with a large-scale generative transformer

**DOI:** 10.1101/2025.01.13.632824

**Authors:** Minuk Ma, Wilson Tu, Carlos Vasquez-Rios, Jiarui Ding

**Affiliations:** Department of Computer Science, University of British Columbia, Vancouver, British Columbia, Canada, V6T 1Z4

**Keywords:** single-cell TCR-sequencing, TCR-peptide interaction, sequence-to-sequence generation, semi-supervised learning, deep learning, transformer

## Abstract

Single-cell TCR sequencing enables high-resolution analysis of T Cell Receptor (TCR) diversity and clonality, offering valuable insights into immune responses and disease mechanisms. However, identifying cognate epitopes for individual TCRs requires complex and costly functional assays. We address this challenge with EpitopeGen, a largescale transformer model based on the GPT-2 architecture that generates potential cognate epitope sequences directly from TCR sequences. To overcome the scarcity of TCR-epitope binding pairs (≈ 100, 000), EpitopeGen uses a semi-supervised learning method, termed BINDSEARCH, which searches over 70 billion potential pairs and incorporates high binding affinity predictions as pseudo-labels. To incorporate CD8^+^ T cell biology into the model as an inductive bias, EpitopeGen employs a novel data balancing method, termed Antigen Category Filter, that carefully controls antigen category ratios in its training dataset. EpitopeGen significantly outperforms baseline approaches, generating epitopes with high binding affinity, diversity, naturalness, and biophysical stability. Notably, the epitopes generated by EpitopeGen follow biologically plausible antigen category distributions, a crucial feature not achieved by other models. Using EpitopeGen, we directly identify subsets of clonally expanded tumorinfiltrating lymphocytes that recognize tumor-associated antigens, exhibiting elevated cytotoxicity and reduced exhaustion markers. From COVID-19 patients, EpitopeGen detects T cells that recognize COVID-19 spike proteins and non-structural proteins with distinct transcriptomic characteristics. In conclusion, EpitopeGen represents the first computational method that enables direct inference of antigen recognition profiles of CD8^+^ T cells from plain TCR repertoires.

## 1 Introduction

The adaptive immune system is a specialized defense mechanism in vertebrates that provides long-lasting protection against pathogens by recognizing and memorizing specific antigens. T cells play a vital role in identifying and eliminating infected cells through their unique T cell receptors (TCRs). CD8^+^ T cells, in particular, inspect endogenous peptides displayed on class I Major Histocompatibility Complex (MHC) molecules, expressed ubiquitously across human cells^1^. Upon recognition of abnormal peptides, such as those of viral or tumoral origin, cytotoxic CD8^+^ T cells can initiate apoptosis in the target cells, given appropriate co-stimulatory signals^2^. The specific antigen fragment recognized by the immune system is termed an epitope. For T cells, epitopes are peptide fragments presented on MHC molecules, serving as key targets for TCR recognition. The Complementarity Determining Region 3 (CDR3) of the TCR is primarily responsible for epitope binding, with the interaction affinity determined by the physicochemical properties of both protein sequences. The extreme polymorphism of CDR3, resulting from VDJ recombination^3, 4^, enables a diverse range of immune responses but presents challenges for quantitative modeling due to the vast sequence diversity at the TCR-pMHC (peptide-loaded MHC) interface.

Prior works in computational TCR analysis can be broadly grouped into three categories: diversity metrics for TCR repertoire analysis, TCR clustering methods, and TCR-pMHC binding affinity prediction models. To quantify the focused nature of immune responses, various TCR diversity metrics have been proposed, including Shannon entropy^5^, Simpson’s diversity index^6^, and Rényi diversity^7, 8^. Janarthanam et al^9^. employ D50, the minimum number of unique clonotypes constituting 50% of the total, to track clonality changes in pediatric eosinophilic esophagitis patients during dietary interventions. TCRDivER^10^ introduces a similarity-sensitive diversity measure that jointly considers clone size and sequence similarity. While these diversity metrics effectively capture T cell clonal expansion patterns, they provide limited insight into antigen specificity. Analysis of TCR repertoire data has been facilitated by various TCR clustering methods^11–14^. These methods aim to group TCRs with potentially similar antigen specificity. GLIPH2^11^ uses a statistical method to identify TCR motifs that are overrepresented in query sets compared to background TCR repertoires. DeepTCR^13^ leverages deep learning-based autoencoders to learn latent TCR representations that facilitate clustering of similar sequences. However, these clustering approaches, while valuable for repertoire analysis and motif discovery, face two key limitations. First, TCRs within the same cluster may not share antigen specificity. Second, the methods cannot directly predict the cognate epitopes for the identified TCR clusters. Recent advances in machine learning have enabled direct prediction of TCR-pMHC binding affinity^15–27^. Models such as pMTnet^15^ address data scarcity through transfer learning, while PanPep^16^ employs model-agnostic meta-learning to enhance generalization to unseen data. However, these binding affinity prediction models have limited utility in analyzing TCR repertoires, as repertoire data typically lack corresponding epitope information, creating a barrier to functional analysis.

To address these challenges, we explore the generative modeling of epitope sequences, inspired by the success of large language models in open-ended text generation^28–30^. By developing an efficient algorithm to identify cognate epitopes of TCRs, we aim to bridge the gap between TCR repertoire data and functional analysis. This approach can provide valuable insights for advancing personalized medicine and enhancing patient outcomes. For example, identifying tumor-specific TCRs can improve the precision of cancer therapies, allowing better targeting of cancer cells while minimizing harm to healthy tissues^31^. Furthermore, accurately identifying TCRs that respond to viral epitopes from a patient can help in the design of more effective, customized vaccines^32^.

We introduce EpitopeGen, a large-scale generative transformer model that predicts cognate epitope sequences from TCR sequences. We adopt a decoder-only architecture to maximize the model’s capacity to learn the conditional probability distribution of epitope sequences given TCR inputs. The self-attention mechanism captures the relationships between TCR tokens and generated epitopes. To leverage the large number of unpaired TCR and epitope sequences from high-throughput experiments, we systematically evaluated over 70 billion TCR-epitope pairs and selected high-confidence interactions based on predicted binding affinity. A key innovation in our approach is the Antigen Category Filter (ACF), a novel data balancing method that calibrates the distribution of antigen categories in our training set according to established immunological principles of CD8^+^ T cell recognition^33–48^. These distributional constraints were essential for establishing an appropriate prior in the generative model, ensuring biological plausibility when applied to repertoire-level analysis.

To the best of our knowledge, EpitopeGen represents the first sequence-to-sequence generative model for predicting epitopes from TCRs. We evaluate the generated epitopes across multiple dimensions, including binding affinity, chemical properties, and naturalness. The results show that the generated epitopes exhibit high binding affinity to the input TCRs and possess chemical properties similar to those of natural epitopes. In particular, most epitopes could be successfully matched to entries in known epitope databases^49^. In repertoire-level evaluations, EpitopeGen generates diverse epitopes that conform to natural antigen category distributions. We next employ molecular dynamics simulations^50^ as an orthogonal evaluation method to deep learning-based affinity predictors, further corroborating the high binding affinities of the generated epitopes. Finally, we apply EpitopeGen to two practical scenarios: (1) identifying and quantifying tumor-associated CD8^+^ T cells within the tumor microenvironment^51, 52^, and (2) characterizing COVID-19-associated peripheral blood mononuclear CD8^+^ T cells. EpitopeGen enables the analysis of the abundance, clonal expansion, and differentially expressed genes of disease-associated CD8^+^ T cells. Tumor-associated CD8^+^ T cells in cancer patients, especially those clonally expanded in both tumor and normal adjacent tissues, exhibit significantly enhanced cytotoxic markers and reduced exhaustion signatures. These results support the previous hypothesis that tumor-specific CD8^+^ T cells are continuously replenished from the periphery^52^. EpitopeGen identifies COVID-19-associated T cells that show clonal expansion and clear cytotoxic profiles in patients with mild and moderate symptoms. In contrast, COVID-19-specific T cells in severe patients retain naive and memory phenotypes rather than acquiring cytotoxic effector functions, potentially explaining the impaired viral control in these individuals. In conclusion, EpitopeGen enables more accurate quantification of T cell repertoires and may lead to novel insights into disease mechanisms.

## 2 Results

### Developing a large-scale transformer to generate cognate epitopes

EpitopeGen is a decoder-only transformer based on GPT-2^28^. EpitopeGen takes as input the CDR3*β* sequence of a TCR and outputs its potential cognate epitope sequences. The original GPT-2 model contains 1.5 billion parameters and was trained on natural language data derived from approximately 800 million Web pages. However, the currently available TCR-epitope binding pairs number ∼ 100, 000, which poses a significant challenge in training an effective deep learning model. To address this limitation, we propose a semi-supervised learning method, termed BINDSEARCH, that incorporates high-confidence predicted TCR-epitope pairs as pseudo-labels for model training (**Fig. 1a**).

**Figure 1:**
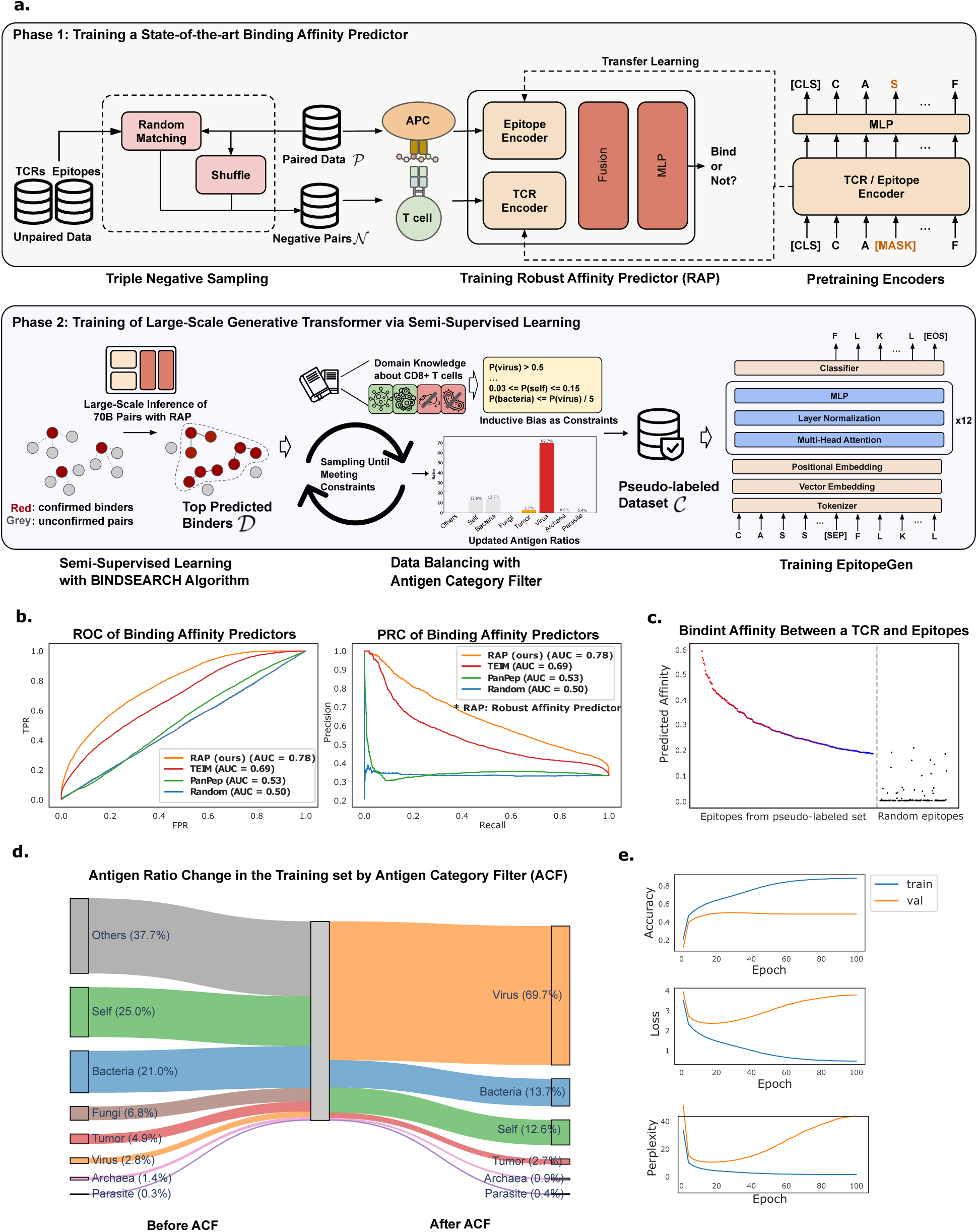
Overview of EpitopeGen development process based on semi-supervised learning and systematic data balancing. **a.** Core development process of EpitopeGen. In the first phase, the encoders for TCRs and epitopes are trained based on the masked amino acid prediction task. Triple Negative Sampling was proposed to diversity the sources of negative samples to train Robust Affinity Predictor (RAP), which reliably predicts the binding affinity between a TCR and an epitope. In the second phase, to leverage unlabeled data for semi-supervised learning, largescale inference was performed by randomly pairing unpaired TCRs and epitopes to find potential binders. Antigen Category Filter, an essential component for providing an inductive bias, aligns the antigen source distribution of the epitopes in the dataset with the reference antigen category distribution through iterative sampling. The resulting pseudo-labeled data are used to train a GPT-2 small architecture, which learns to generate epitope sequences from CDR3*β* inputs. **b.** ROC and PRC plots of binding affinity predictors, RAP (orange), TEIM (red), PanPep (green), and Random (blue) on the test set merged from four publicly available datasets. The negative pairs were obtained by shuffling positive pairs. **c.** Binding affinity distribution for an example TCR “CAISEGTWETQYF” against epitopes from pseudo-labeled and random datasets. The yaxis shows binding affinity, while the x-axis shows epitope sources. Left shows the binding affinity between TCRs and the epitopes from pseudo-labeled data in decreasing order, and right shows the binding affinity between TCRs and arbitrary epitopes. **d.** Antigen category distributions in pseudo-labeled datasets before and after the application of the Antigen Category Filter (ACF). Left: Pre-ACF distribution, with many ‘Others’, ‘Self’, and ‘Bacteria’ as the dominant categories. Right: Post-ACF distribution, showing a shift to ‘Virus’ as the predominant category. ‘Self’ and ‘Tumor’ denote self-antigen and tumor-associated antigens, respectively. **e.** The learning curve of EpitopeGen. The average masked token prediction accuracy (top), loss (middle), and perplexity (bottom) values (y-axis) on the training and validation set are plotted by epoch (x-axis).

BINDSEARCH requires a reliable binding affinity predictor. We therefore developed the Robust Affinity Predictor (RAP) in the first phase, built upon TABR-BERT^24^. RAP features a custom BERT^53^ architecture pre-trained on peptide sequences through masked amino acid prediction, followed by an MLP head optimized with the cross-entropy loss. A key challenge in training binding predictors is the preparation of validated non-binding pairs. To address this challenge, we introduced Triple Negative Sampling, a technique that diversifies the sources of negative pairs to minimize potential biases. Given a set of paired data, the negative pairs are generated by randomly shuffling the paired data or pairing them with external TCRs or epitopes (**Methods**). The paired data were collected from four publicly available sources: VDJdb^54^, IEDB^55^, PIRD^56^, and McPAS-TCR^57^. After preprocessing, the paired data were partitioned into training, validation, and test sets with a 7:2:1 ratio. Robust Affinity Predictor demonstrated superior or comparable performance to previous models, including TABR-BERT, PanPep, and TEIM^25^, as measured by the Area Under the Receiver Operating Curve (AUROC) and Area Under the Precision-Recall Curve (AUPRC) metrics, in multiple test sets (**Fig. 1b** and **Extended Data Fig. 1**).

We collected a pool of 7, 331, 478 unique TCR sequences from TCRdb^58^. The 21,801,187 epitope sequences were obtained from three previous publications: NetMHCPan v4.0^59^, MHCflurry v2.0^60^, and SysteMHC^61^ (**Supplementary Note 1**). In this study, we focus specifically on the binding between CD8^+^ T cells and epitopes displayed on class I MHC molecules, while excluding interactions involving CD4^+^ T cells and epitopes presented by class II MHC molecules. Class I MHC molecules bind and present short epitopes (8-12 amino acids) to CD8^+^ T cells in a restricted manner, which warrants a focused modeling approach^62^.

In the second phase, BINDSEARCH assessed the binding affinities of each unpaired TCR against 10,000 random epitopes sampled from a set of unpaired epitopes, identifying pairs exhibiting significantly higher affinities. As expected, the majority of TCR-epitope combinations showed negligible binding affinities; for example, the TCR sequence CAISEGTWETQYF exhibited a median binding affinity of 0.000 against randomly selected epitopes (**Fig. 1c**). However, we identified a subset of high-affinity interactions, with the top 256 predicted binders showing a median affinity of 0.429. To ensure specificity and reduce potential false positives, we filtered out promiscuous epitopes that were predicted to bind more than 100 different TCRs. This systematic pseudo-labeling approach generated an intermediate dataset D, containing 17 million TCR-epitope pairs.

A critical consideration in constructing the pseudo-labeled dataset was ensuring a biologically plausible distribution of antigen sources (e.g., viral, bacterial, and self-antigens). Although there are no universally accepted ratios for antigen-specific CD8^+^ T cells, virus-specific cells often constitute a significant portion^33–35^, while self-antigen and tumor-antigenspecific CD8^+^ T cells generally occur at much lower frequencies in healthy individuals^36–39^. Our goal was to design a training set that effectively represented a large-scale TCR repertoire so that the model can learn not only the binding affinity relationships but also the natural distributional patterns. We proposed the Antigen Category Filter (ACF) algorithm, which calibrates antigen category distribution based on five fundamental immunological principles of CD8^+^ T cells: (1) Viral Dominance, (2) Limited Bacterial Presence, (3) Endogenous Epitope Presence (Self-Antigen and Tumor-Associated Antigen), (4) Rare Fungi and Parasites, and (5) Absence of Pathogenic Archaea (**Methods**, **Supplementary Note 2**). Initial analysis of dataset D revealed that 37.7% of epitopes were derived from others (mostly Eukaryotic), potentially introducing undesirable bias. After applying the Antigen Category Filter, the resulting dataset C, which we call Corpus, predominantly consisted of virus-associated epitopes (**Fig. 1d**).

We developed a custom tokenizer, trained on C, to capture frequently occurring motifs in CDR3*β* and epitope sequences. The input sequences were processed using the tokenizer and augmented with positional embeddings for model training (**Methods**). The model architecture was a GPT-2-small, trained for 100 epochs using four NVIDIA L40S GPUs. The validation loss and perplexity initially decreased smoothly but began to increase after epoch 20 (**Fig. 1e**). The maximum next-token prediction accuracy reached 0.8843 (at epoch 100) and 0.5026 (at epoch 28) for the training and validation sets, respectively. To mitigate potential overfitting, we selected the model checkpoint from epoch 28 for subsequent analyses. For inference, we used top-*k* top-*p* sampling^63, 64^ as the decoding scheme to generate the most probable *k* tokens for a given input TCR, selecting from tokens whose cumulative probability reaches *p* to enhance diversity in the output.

### EpitopeGen generates high-affinity, diverse, and biologically sane epitopes

We evaluated the binding affinities between the input TCRs and the generated epitopes across multiple test scenarios. We partitioned the test set of Corpus C into four subsets based on TCR and epitope exposure during training: UnseenEpi (TCRs seen, epitopes unseen), UnseenTCR (epitopes seen, TCRs unseen), SeenBoth (both seen, but not as a pair), and UnseenBoth (neither seen). EpitopeGen generated epitope sequences for each TCR in the test sets. We then measured the binding affinity between each TCR-epitope pair using the Robust Affinity Predictor. For comparison, we also measured the binding affinities between each TCR and 100 randomly sampled epitopes and calculated the percentile ranks of the binding affinities. The mean percentile ranks for UnseenEpi, UnseenTCR, SeenBoth, and UnseenBoth were 81.5, 81.3, 81.9, and 81.2, respectively (**Fig. 2a**). These results indicated that EpitopeGen can generate epitopes with high binding affinities for both previously encountered and novel TCRs.

**Figure 2:**
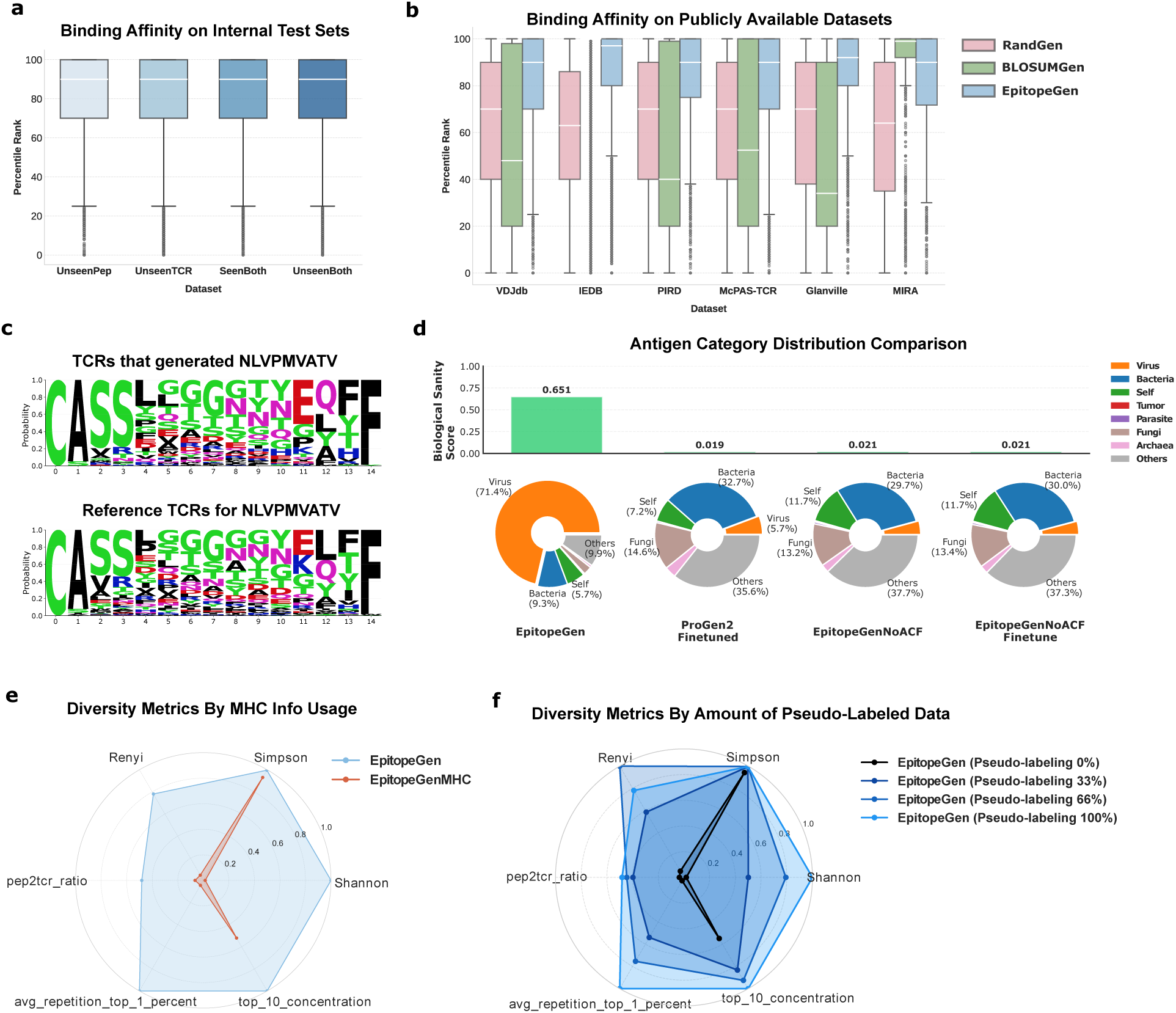
Evaluation of the binding affinity, diversity, and distributional characteristics of the generated epitopes. **a.** Box-and-whisker plots showing mean percentile rank (y-axis) of binding affinities between generated epitopes and query TCR sequences across four subsets of the pseudo-labeled test set (x-axis): UnseenEpi, UnseenTCR, SeenBoth, and UnseenBoth. For each generated epitope, 100 random epitopes were sampled to calculate the percentile rank of binding affinity. **b.** Box-and-whisker plots showing mean percentile rank (y-axis) of binding affinities between generated epitopes and query TCR sequences across test sets of six publicly available datasets (x-axis). For each generated epitope, 100 random epitopes were sampled to calculate the percentile rank. Error bars indicate 95% confidence intervals. Binding affinities were measured using Robust Affinity Predictor. **c.** Logomaker plots comparing TCRs that generated the epitope NLVPMVATV with reference TCRs known to recognize this epitope from the VDJdb dataset. **d.** Distributions of antigen categories for epitopes generated by four models: EpitopeGen, ProGen2 Finetuned, EpitopeGenNoACF, and EpitopeGenNoACFFinetune on the 10x dataset. The Biological Sanity Score reflects how similar the distribution of antigens is to the reference distribution which occurs in real biological systems. ‘Self’ denotes self-antigens, and ‘Tumor’ represents tumor-associated antigens. **e,f** Radar plots comparing six diversity indices of generated epitopes using: **e**, EpitopeGen versus EpitopeGenMHC; **f**, model variants trained with different proportions of pseudo-labeled data (0%, 33%, 66%, and 100%). Metrics include Rényi diversity (*α*=2), Shannon diversity, Simpson’s diversity index, Epi-to-TCR ratio, avg repetition top 1 percent, and top 10 concentration. For visualization, unbounded metrics were normalized by their maximum values, and inverse values were used for the last two metrics. Analysis based on the 10x dataset.

We evaluated EpitopeGen using test sets (VDJdb, IEDB, PIRD, and McPAS-TCR), previously defined during the development of the Robust Affinity Predictor. Two baseline methods were implemented for comparison: RandGen, which generates random amino acid sequences based on the training set’s epitope length distribution, and BLOSUMGen, which assigns epitopes from the training set based on TCR sequence similarity. BLOSUMGen first filters candidate TCRs using Levenshtein distance when the search space exceeds 500 sequences, then identifies nearest neighbors using the BLOSUM62 substitution matrix with global sequence alignment.

EpitopeGen-generated epitopes demonstrated consistently high binding affinities across multiple test sets, with mean percentile ranks exceeding 80%. In evaluations using VDJdb, PIRD, and McPAS-TCR test sets, EpitopeGen outperformed both RandGen and BLOSUMGen while maintaining comparable performance on IEDB (**Fig. 2b**). External validation using independent experimental datasets from Glanville et al.^65^ and Nolan et al.^66^ (**Supplementary Note 1**) further confirmed EpitopeGen’s robustness, achieving mean percentile ranks of 85.8 and 84.1, respectively (**Fig. 2b**).

Leveraging the ability of GPT-2 to produce multiple, slightly varied outputs for a single input, we analyzed the top 1, 5, 10, and 20 generations. In this extended evaluation, EpitopeGen consistently showed higher mean percentile ranks on public test sets compared to RandGen and BLOSUMGen (**Extended Data Fig. 2**).

To demonstrate EpitopeGen’s predictive capabilities, we analyzed TCR sequences associated with NLVPMVATV, the most frequently occurring epitope in the VDJdb dataset, comparing TCRs that generated this epitope against experimentally validated TCRs using Logomaker^67^. The generated sequences exhibited characteristic amino acid patterns with notable variability in their central regions. Both the generated and experimentally validated sequences displayed conserved motifs: an N-terminal ‘CASS’ pattern, a central ‘LGGGGYE’ sequence, and a C-terminal ‘QFF’ motif (**Fig. 2c**). This consistent pattern alignment demonstrates EpitopeGen’s ability to generate epitope sequences from TCRs that are similar to experimentally validated ones.

We further evaluated EpitopeGen’s performance using repertoire-level test sets, specifically the 10x dataset^68^ released by 10x Genomics, Inc., which contains paired RNA and TCR sequencing of CD8^+^ T cells from a healthy donor. Therefore, this evaluation closely mirrors real-world application scenarios in which EpitopeGen is used to infer epitopes for an individual’s entire TCR repertoire. For comparative analysis, we fine-tuned ProGen2^69^, a leading protein language model, on the publicly available training set. We also developed three variants of EpitopeGen: EpitopeGenNoACF, EpitopeGenNoACFFinetune, and EpitopeGenMHC. EpitopeGenNoACF was trained on the intermediate dataset D without applying Antigen Category Filter, thus lacking inductive bias on the proper distribution of antigen categories. EpitopeGenNoACFFinetune was derived by fine-tuning EpitopeGenNoACF using C to assess whether fine-tuning could effectively learn the distribution of antigen categories. EpitopeGenMHC incorporated both TCR and MHC information for epitope sequence generation. We evaluated the models in three key aspects: distribution of antigen categories of the generated epitopes (measured by Biological Sanity Score), epitope diversity, and redundancy in multiple epitopes generated from the same TCR (**Methods**). Only models that demonstrated high performance across all three metrics were considered eligible for downstream analysis.

EpitopeGen significantly outperformed all other models, achieving the Biological Sanity Score of 0.651. The generated epitopes by EpitopeGen predominantly originated from viruses (**Fig. 2d**), with smaller proportions from tumoral, self, and bacterial sources. This distribution aligns with the immunological principles of Viral Dominance, Limited Bacterial Presence, and Endogenous Epitope Presence. In contrast, all other models (ProGen2Finetuned, EpitopeGenNoACF, and EpitopeGenNoACFFinetune) generated approximately 37.7% and 30% of epitopes from ‘Other’ (mostly Eukaryotic) and ‘Bacteria’ with notably fewer viral antigens. These skewed antigen category distributions of generated epitopes reflect the initial bias in the dataset D before the application of Antigen Category Filter, highlighting the importance of careful data balancing approach.

To assess epitope diversity, we employed six diversity indices: Shannon diversity^5, 8^, Rényi diversity (*α*=2)^7^, Simpson’s diversity index^6^, the Epi-to-TCR ratio (unique epitopes/number of TCRs), avg repetition top 1 percent, and top 10 concentration (proportion of epitopes in most frequent 10%) (**Methods**). EpitopeGen generated epitopes showed superior diversity, achieving an epitope-to-TCR ratio of 0.5 (**Fig. 2e**). In contrast, EpitopeGenMHC results showed significantly lower diversity indices, indicating the generation of redundant epitopes for different TCRs. This limitation stems from the insufficient diversity of (TCR, epitope, MHC) triplets in the currently available datasets. Furthermore, EpitopeGen exhibited lower redundancy in its top 20 generations compared to EpitopeGenMHC (**Extended Data Fig. 3a**).

To prove the effectiveness of semi-supervised learning, we evaluated models trained with different proportions of unlabeled data (pseudo-labeled by BINDSEARCH): 0%, 33%, 66%, and 100% (denoted as Pseudo-labeling 0%, 33%, 66%, and 100%, respectively). The Pseudolabeling 0% model, trained solely on the public paired dataset, showed considerable redundancy, with the top 1% epitopes repeating approximately 2,000 times on average and a top 10 concentration exceeding 0.95 (**Fig. 2f**). In contrast, Pseudo-labeling 100% achieved much greater diversity with a top 10 concentration below 0.50. Epitope diversity improved progressively with increasing proportions of pseudo-labeled data, highlighting the advantage of incorporating unlabeled data in ensuring the generation of diverse epitopes. Moreover, redundancy in the top-*k* generations decreased gradually with higher proportions of pseudolabeled data for training (**Extended Data Fig. 3b**).

Repertoire-level evaluations were instrumental in informing three key design choices: (1) implementing a semi-supervised learning approach to incorporate pseudo-labeled data for model training, (2) providing inductive bias about CD8^+^ T cell biology using Antigen Category Filter, which led to the exclusion of EpitopeGenNoACF and EpitopeGenNoACFFinetune, and (3) excluding MHC information, resulting in the removal of EpitopeGenMHC.

### EpitopeGen generates natural epitopes

We next examined the naturalness of the generated epitopes by comparing their various properties with those of naturally occurring epitopes collected from the test sets. The generated epitopes had an average length of 10.08, which aligns well with the typical length range^70^ (8 to 12 amino acids) of the epitopes loaded onto MHC class I molecules (**Fig. 3a**). Additionally, the amino acid usage patterns of the generated epitopes closely resembled those of natural epitopes (Pearson correlation = 0.911; **Fig. 3b**).

**Figure 3:**
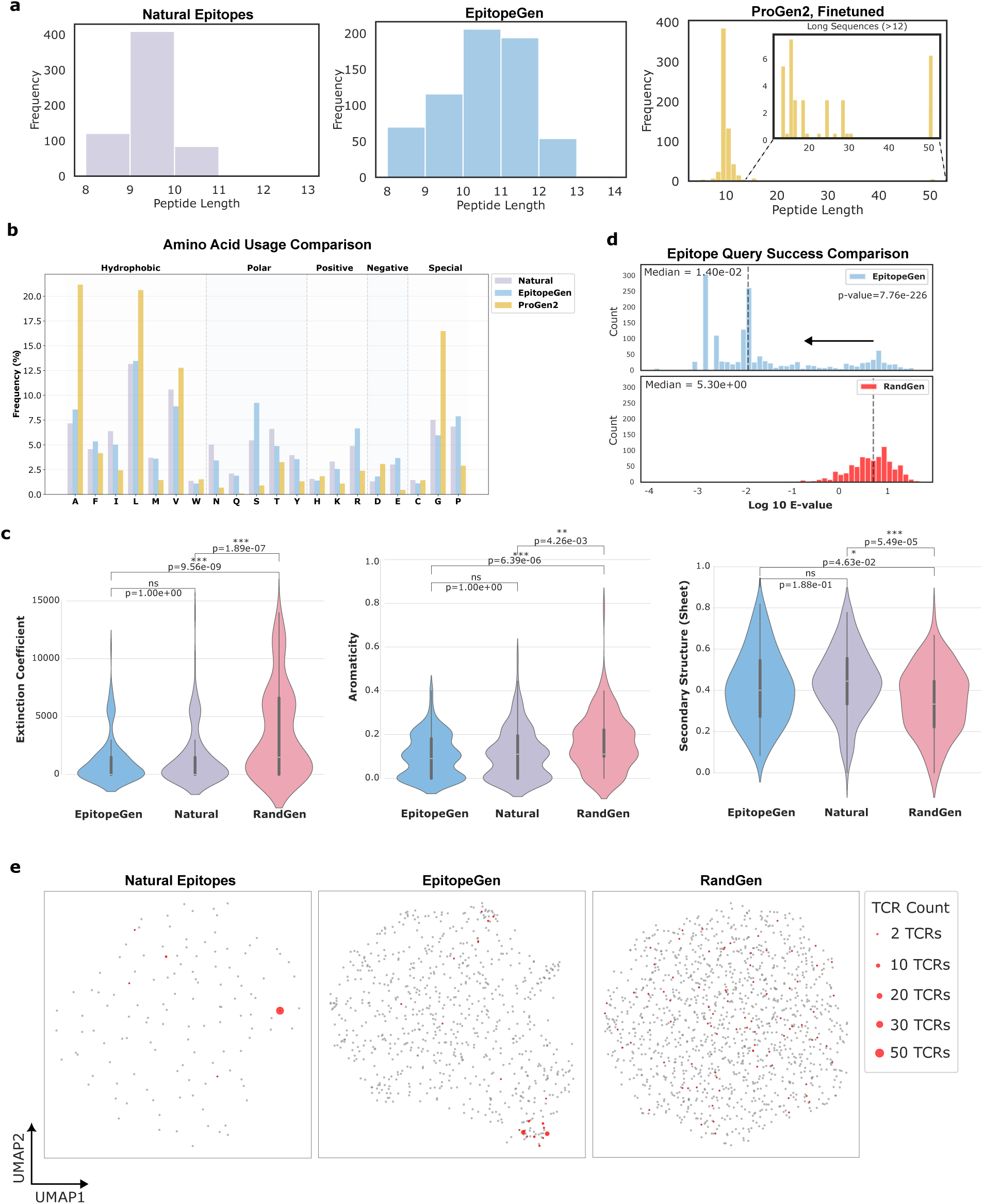
Validation of naturalness of the epitopes generated by EpitopeGen. **a.** Length distribution comparison between natural epitopes, epitopes generated by EpitopeGen, and those generated by ProGen2 fine-tuned on the public training set based on the VDJdb test set. Natural epitopes longer than 12 amino acids were excluded during preprocessing. **b.** Comparison of amino acid usage between natural epitopes (Natural), EpitopeGen-generated epitopes, and ProGen2-generated epitopes. **c.** Distribution of chemical properties for epitopes from EpitopeGen (blue), Natural (purple), and RandGen (red). Natural epitopes were sampled from the VDJdb test set (n=642). Chemical properties include extinction coefficient, aromaticity, and secondary structure (sheet). Statistical significance was assessed using two-sided Mann-Whitney U tests with Bonferroni correction for multiple comparisons. Each violin plot contains an embedded box plot displaying the median, interquartile range (IQR), and whiskers extending to 1.5 x IQR. ns means ‘not significant’, *p *<* 0.05, **p *<* 0.01, and ***p *<* 0.001. **d.** Distribution of BLASTP E-values for epitopes generated by EpitopeGen (blue, upper) and RandGen (red, lower). Each epitope was queried against the SwissProt database to identify source proteins. Lower E-values indicate higher statistical significance. Statistical comparison was performed using a one-sided Mann-Whitney U test. **e.** UMAP visualization of epitope space comparing Natural, EpitopeGen, and RandGen epitopes. Epitope similarity was computed using the BLOSUM62 matrix. Each dot represents an epitope, with dot size indicating the number of paired TCRs. Colored dots represent epitopes recognized by TCRs sharing the GLIPH2 motif “CASSIRSQETQYF”. Results are based on the PIRD test set, with Natural epitopes sampled from this set.

Recent protein language models, such as ProtGPT2^71^, ProGen^72^, and ProGen2^69^, demonstrated protein sequence generation capabilities, but lack precise control mechanisms for specialized tasks such as the generation of CD8^+^ T cell epitopes. Although ProGen incorporates conditioning through gene ontology keywords^73^ and taxonomic tags, adapting these models for specific tasks such as TCR-specific epitope generation remains non-trivial. To evaluate the utility of pre-trained models, we fine-tuned ProGen2 on public training data. Our experiments revealed that the fine-tuned model often generated epitopes exceeding the biological length constraints, exhibiting a long-tailed length distribution (**Fig. 3a**). This behavior stems from ProGen2’s pre-training on general protein sequences, which are typically longer than T-cell epitopes that naturally range between 8 and 12 amino acids. Specifically, 4.41% of the generated sequences were longer than 12, while 1.47% were shorter than 8 amino acids. The model also showed different amino acid usage patterns compared to natural epitopes (**Fig. 3b**). When tested on the 10x dataset, the fine-tuned ProGen2 model exhibited severe bias, with 83.80% of generated sequences starting with ‘G’. These findings suggest that strongly conditional generation tasks, where a single amino acid difference can significantly impact binding properties, require enhanced supervision through carefully curated training data that satisfy biological constraints.

To assess the chemical feasibility of the generated epitopes, we analyzed their several key properties (**Methods**) using the `ProtParam` package^74^. The distributions of these chemical properties in EpitopeGen-generated epitopes closely mirrored those of natural epitopes, while randomly generated epitopes showed significantly different distributions (**Fig. 3c**). These trends were consistently observed in four independent public test sets (**Extended Data Fig. 4**).

The generated epitopes were queried against the SwissProt protein database using `blastp`^75^ to assess the presence of significant matches (**Methods**). The significance of a `blastp` search is quantified by the E-value, which represents the expected number of alignments with a score equal to or larger than the observed one by chance. The generated epitopes showed significantly lower E-values (Two-Tailed Mann-Whitney *U* test, p=1.23 × 10^−244^) compared to randomly generated epitopes, suggesting a significantly higher likelihood of identification within the database (**Fig. 3d**).

Natural and generated epitopes from the same test sets were qualitatively compared through visualization. Epitope embeddings were computed using pairwise BLOSUM similarity, followed by UMAP dimensionality reduction. Within each UMAP, epitopes paired with TCRs in the same motif group (defined using `GLIPH2`^11^) were color-highlighted. We observed that similar TCRs (in the same motif group) recognize similar epitopes (aggregated colored dots) in both natural epitopes and EpitopeGen-generated epitopes (**Fig. 3e, left, middle**). The colored dots representing EpitopeGen-generated epitopes showed localized aggregation with slight variation. These observations suggest that EpitopeGen produces diverse epitopes that extend beyond the range found in publicly available datasets while preserving recognition patterns of TCR motif groups. In contrast, the colored dots representing RandGen-generated epitopes appeared widely dispersed (**Fig. 3e, right**). These patterns were consistent across multiple TCR motif groups and datasets (**Extended Data Fig. 5**).

### EpitopeGen generates biophysically stable epitopes

To analyze protein structures and interactions, we use Molecular Dynamics (MD) simulations, which compute molecular movements over time based on physical and statistical principles^50^. Specifically, we utilized `InterfaceAnalyzer`^76^, from the `Rosetta` suite^77^, to estimate TCR-pMHC complex binding affinity. For each CDR3*β*, EpitopeGen generated a corresponding epitope sequence. The complete 3D structure of the TCR-pMHC complex was then constructed using `TCRmodel2`^78^, powered by AlphaFold2^79^ (**Fig. 4a**). All simulations assumed the prevalent MHC allele HLA-A*02:01^80^ and utilized a fixed TCR alpha chain (**Methods**). **Fig. 4b** shows an example of the 3D structure featuring an EpitopeGen-generated epitope. The magnified interface between CDR3*β* (CAVSPLGGSQGNLIF) and the epitope (GILGFVFTLS) highlights a hydrogen bond between tyrosine in the epitope and glycine in the CDR3*β*, contributing to the stability of the complex.

**Figure 4:**
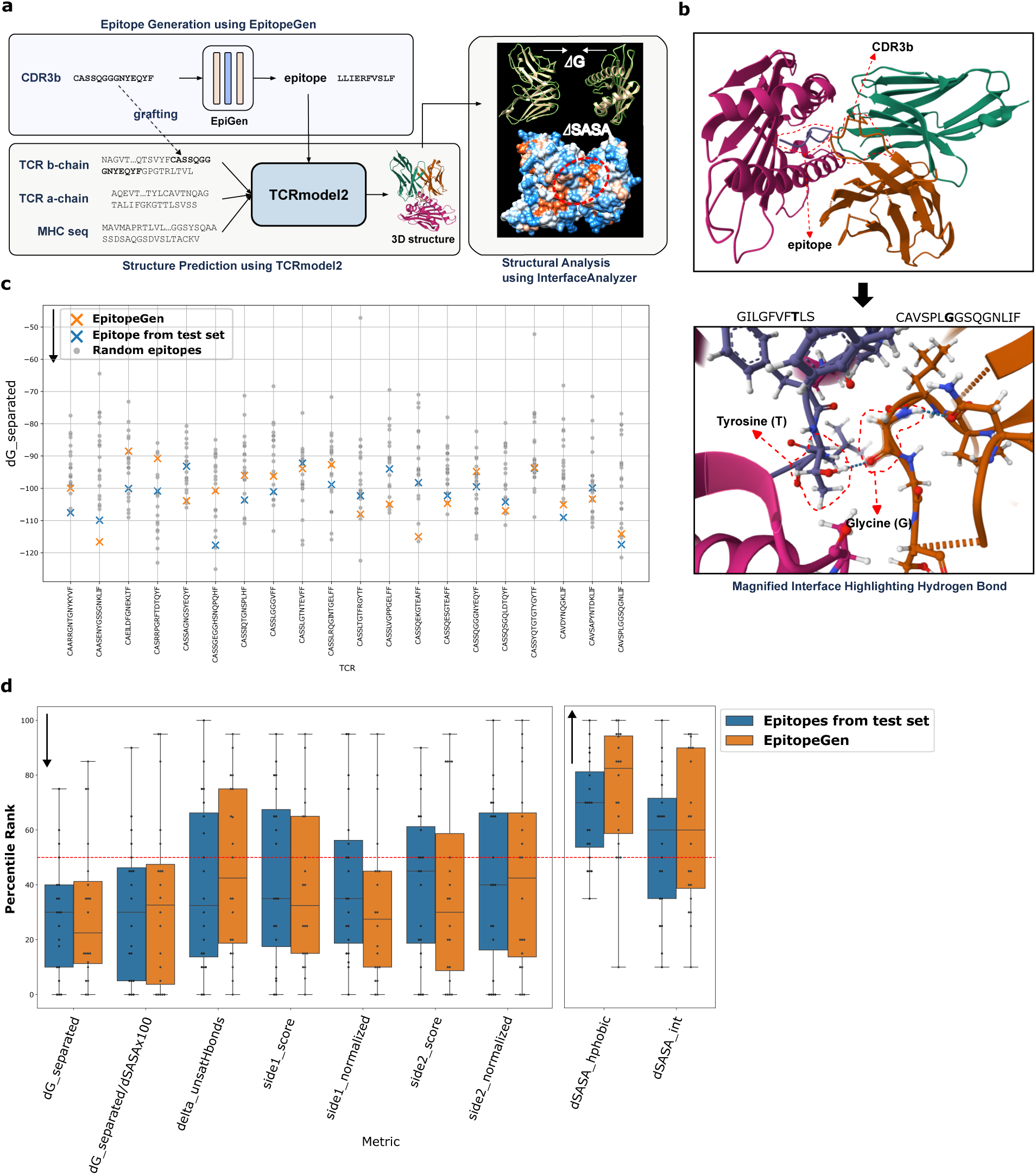
Structural analysis on the TCR-pMHC complexes with the epitopes generated by EpitopeGen using InterfaceAnalyzer. **a.** A schematic showing the structural analysis pipeline. An epitope generated by EpitopeGen is fed to TCRmodel2, which outputs the 3D structure of the TCR-pMHC complex in PDB format. The file is analyzed using Rosetta Relax and InterfaceAnalyzer, which calculate diverse physicochemical properties. The 3D structures are visualized using Mol* Viewer^83^ and Chimera^84^. **b.** Visualization of the predicted 3D structure of a TCR-pMHC complex containing an epitope generated by EpitopeGen. The epitope and CDR3*β* of the TCR are highlighted. The CDR3*β* sequence is CAVSPLGGSQGNLIF, and the epitope sequence is GILGFVFTLS. The MHC allele HLA02:01 is used. The image is a screenshot taken from Mol* viewer. The lower image shows the detailed view of the TCR-pMHC interface. A hydrogen bond between the epitope’s tyrosine and the CDR3*β*’s glycine is highlighted, illustrating a potential binding mechanism for the generated epitope. **c.** Binding stability analysis of TCR-pMHC complexes. A scatter plot depicting dG separated values (y-axis), for complexes containing EpitopeGen-generated epitopes (orange) or natural epitopes from the VDJdb test set (blue). dG separated means the change in Rosetta energy when the interface-forming chains are separated, versus when they are complexed. Analysis is conducted across 20 TCRs (x-axis), with 20 random epitopes per TCR serving as controls (grey). Lower dG separated values indicate higher structural stability in the energy context. dG separated was calculated using Rosetta’s InterfaceAnalyzer. **d.** Comprehensive analysis of key physicochemical properties in TCR-pMHC complexes. Box plots depict the percentile ranks of nine physicochemical properties related to binding affinity for EpitopeGen-generated epitopes (orange) and natural epitopes from the test set (blue). Each dot represents the percentile rank of a property compared to 20 randomly sampled epitopes. The center red line (50%) indicates the performance of a random baseline. Properties were calculated using Rosetta’s InterfaceAnalyzer. Lower values of the first seven properties (dG separated, dG separated/dSASAx100, delta unsatHbonds, side1 score, side1 normalized, side2 score, side2 normalized) and higher values of the last two properties (dSASA hphobic, dSASA int) indicate more stable structures or potentially stronger binding.

We next evaluated the binding affinity of these complexes using two key metrics. Gibbs free energy (dG separated) quantifies the energy difference before and after TCR-pMHC binding, providing a measure of binding affinity in an energy context^81^. Additionally, hydrophobic interactions, crucial forces in protein folding and docking^82^, were also evaluated. For comparison, we sampled 20 random epitopes from the VDJdb test set as controls. EpitopeGen-generated epitopes showed lower Gibbs free energy compared to randomly sampled epitopes (**Fig. 4c, orange**), with a median percentile rank of 22.5%. This suggests that EpitopeGen-generated epitopes form more energetically stable complexes compared to randomly sampled ones. Furthermore, these epitopes exhibited pronounced hydrophobic interactions (**Extended Data Fig. 6**), with a mean percentile rank of 82.5%. This observation supports the idea that the generated epitopes form stronger hydrophobic interactions with the CDR3*β* region, potentially burying hydrophobic regions and contributing to binding stability. The natural epitopes from the test set showed a median percentile rank of 30.0% for Gibbs free energy, indicating that they also form more stable TCR-pMHC complexes than random control epitopes.

We analyzed seven additional properties and visualized the aggregated results (**Fig. 4d**). These properties include: binding free energy per unit area (dG separated/dSASAx100), the number of unsatisfied hydrogen bonds at the interface (delta unsatHbonds), interface energy scores for TCR and pMHC sides (side1 score and side2 score), normalized side scores (side1 normalized and side2 normalized), and total change in Solvent Accessible Surface Area (dSASA int) (**Supplementary Note 3**).

As a sanity check, epitopes from the test set shows mean percentile ranks below 50% on all energy-related properties (dG separated, dG separated/dSASAx100, side1 score, side1 normalized, side2 score, and side2 normalized) and delta unsatHbonds. This aligns with our hypothesis that natural epitopes from the test set form more stable structures compared to randomly sampled epitopes. Notably, EpitopeGen-generated epitopes demonstrated similar trends across all properties, generally showing lower energy-related metrics. In contrast, both natural and generated epitopes displayed higher percentile ranks (≥50%) on properties related to the interaction surface (dSASA hphobic and dSASA int). These results suggest that both natural and generated epitopes exhibit stronger interactions at the interface compared to those of random epitopes.

These findings indicate that EpitopeGen successfully captures key biophysical characteristics of natural epitopes, particularly in terms of energetic stability and hydrophobic interactions.

### EpitopeGen discovers tumor-associated CD8^+^ T cells

CD8^+^ tumor-infiltrating lymphocytes that recognize tumor-associated epitopes are central to cancer immunotherapy research^85^. To identify these tumor-reactive T cells, we performed computational inference on CDR3*β* sequences of CD8^+^ T cells using EpitopeGen to predict their target epitopes. These predicted epitopes were then cross-referenced against a curated database of known tumor-associated epitopes. T cells whose predicted epitopes matched entities in the database were classified as Phenotype-Associated (PA), while those without matches were designated Not-Associated (NA). For robustness, we took an ensemble of the predictions by three independently trained EpitopeGen models (**Supplementary Note 4**). Our underlying hypothesis was that PA T cells, compared to NA T cells, would exhibit enhanced activation signatures in tumor tissues due to their potential recognition of tumor-associated antigens. This EpitopeGen-based stratification of T cells into PA and NA populations enabled comprehensive differential analyses (**Fig. 5a**).

**Figure 5:**
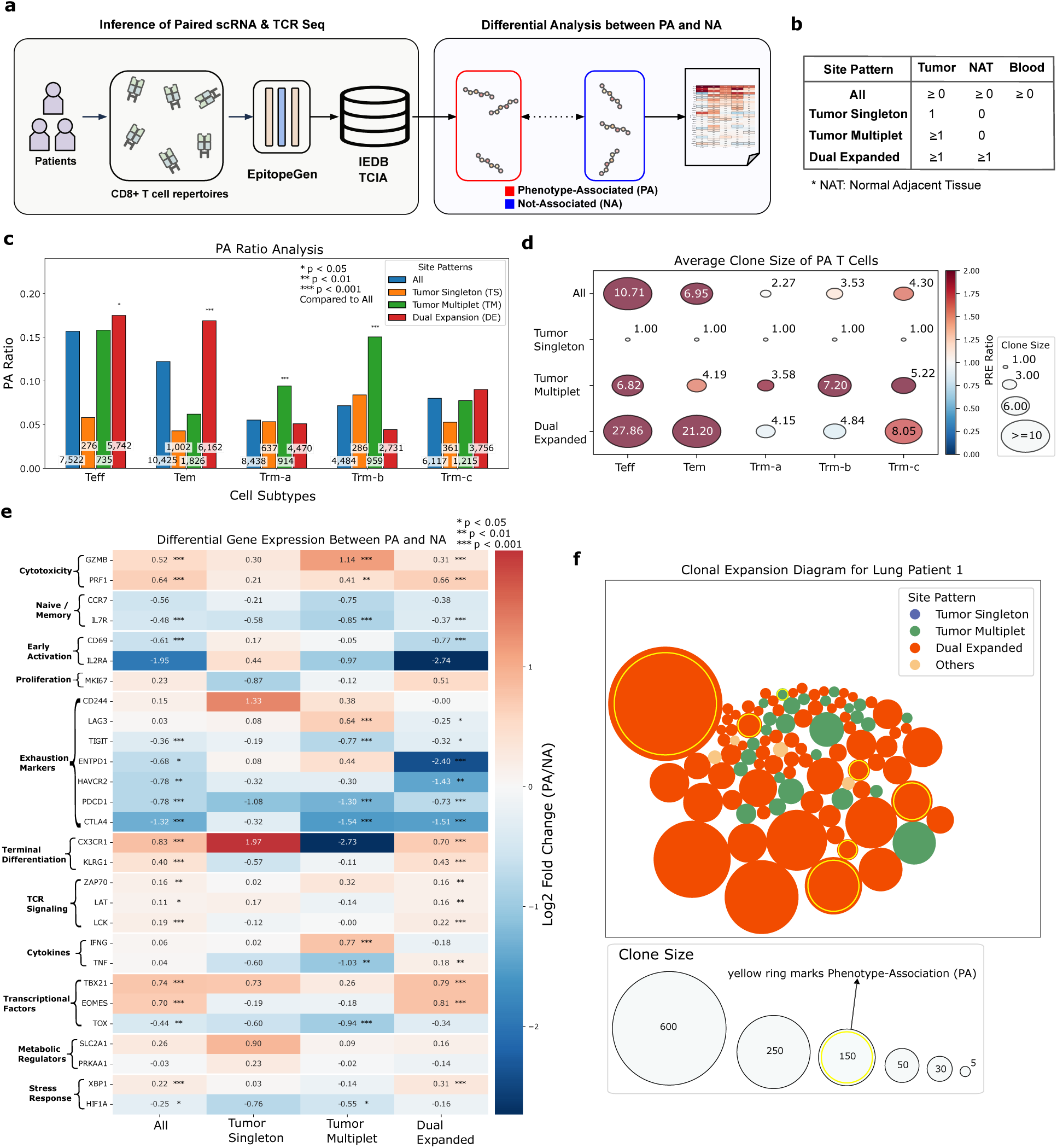
Comparative analysis of Phenotype-Associated (PA) and Not-Associated (NA) T cells by different site patterns in cancer patients. **a.** Schematic of the analysis pipeline. EpitopeGen generates potential epitopes for CD8^+^ cells from paired single-cell RNA and TCR sequencing data. Epitopes are queried against IEDB and TCIA databases to assess tumor association. TCRs and corresponding epitopes are then classified as Phenotype-Associated (PA) or Not-Associated (NA), followed by differential analyses between these groups. **b.** Summary of T cell clone distribution patterns across Tumor, Normal Adjacent Tissue (NAT), and Blood. Three patterns were defined based on clone size: Tumor Singleton, Tumor Multiplet, and Dual Expanded. An additional category, ‘All,’ was included where all cells were considered, regardless of their site patterns. Modified from Wu et al. **c.** Proportions of Phenotype-Associated (PA) TCRs within T cell repertoires, stratified by cell subtype (Teff, Tem, Trm-a, Trm-b, Trm-c) and site pattern (All, Tumor Singleton, Tumor Multiplet, Dual Expanded). One-sided Fisher’s exact tests were performed to test the enrichment of PA T cells in each site pattern to the ‘All’ category within each cell subtype. P-values were adjusted using the Benjamini–Hochberg method (*p *<* 0.05, **p *<* 0.01, ***p *<* 0.001). Sample sizes (n) for each site pattern and cell subtype are shown in each bar. **d.** Mean clone size of Phenotype-Associated (PA) TCRs by cell subtype (x-axis) and site pattern (y-axis). Color intensity indicates the Phenotype-Relative Expansion (PRE) ratio, which is the mean clone size of PA T cells divided by that of NA T cells. Red indicates a larger PRE ratio. Circle size represents the mean clone size of PA T cells. Significant differences between clone sizes of PA and NA T cells were assessed using a one-sided Mann-Whitney *U* test, with p-values corrected for multiple testing via the Benjamini–Hochberg method. Only significant p-values are shown. Sample sizes (n) for site patterns [All, Tumor Singleton, Tumor Multiplet, Dual Expanded]: Teff (n = 7522, 276, 735, 5742), Tem (n = 10425, 1002, 1826, 6162), Trm-a (n = 8438, 637, 914, 4470), Trm-b (n = 4484, 286, 959, 2731), and Trm-c (n = 6117, 361, 1215, 3756). The numbers of PA T cells for site patterns [All, Tumor Singleton, Tumor Multiplet, Dual Expanded]: Teff (1178, 16, 116, 1003), Tem (1272, 43, 113, 1039), Trm-a, (466, 34, 86, 228), Trm-b (321, 24, 144, 121), and Trm-c (490, 19, 94, 338). **e.** Heatmap of differential gene expression between Phenotype-Associated (PA) and Not-Associated (NA) TCRs, ranked by magnitude of difference. Statistical tests were performed using a two-sided Wilcoxon rank-sum test from Scanpy, and p-values were adjusted for multiple testing using the Benjamini-Hochberg method (*p *<* 0.05, **p *<* 0.01, ***p *<* 0.001). Analysis was performed for four site patterns (x-axis): All, Tumor Singleton, Tumor Multiplet, and Dual Expanded. The curated gene list was obtained from Wu et al., containing genes related to CD8^+^ T cells subtypes, states, and activation. **f.** CD8^+^ T cell composition in Lung Patient 6, combining data from lung tumor tissue, Normal Adjacent Tissue (NAT), and blood data. The top 100 expanded clones are shown, with circle size representing clone size and color denoting cell subtype. Phenotype-Associated (PA) T cells are marked with yellow rings. The plot provides an overview of the clonal expansion patterns and tumor-associated T cells.

We used a dataset published by Wu et al.^52^ that comprises paired single-cell RNA and TCR sequencing measurements for four types of cancer from three sites: tumor tissue, normal adjacent tissue (NAT), and peripheral blood. The study identified five CD8^+^ T cell subsets: Teff (effector), Tem (effector memory), Trm-a, Trm-b, and Trm-c (tissue-resident memory cells, with cluster names shortened for brevity). Additionally, the study defined site patterns of TCRs based on their clonal expansion patterns in tumor tissue, NAT, and blood. We focused on three site patterns: Tumor Singleton (TCR observed only once in tumor tissue), Tumor Multiplet (TCR observed to be clonally expanded in tumor tissue), and Dual Expanded (TCR observed at least once in both tumor tissue and NAT) (**Fig. 5b**). ‘All’ means all cells were considered, regardless of their site patterns.

We first analyzed the prevalence of PA T cells in different cell subtypes and site patterns by calculating their proportion (PA ratio). PA ratios exhibited marked heterogeneity by cell subtypes: effector T cells (Teff) showed the highest ratio of 0.157, followed by effector memory T cells (Tem) with 0.122, while three subsets of tissue-resident memory T cells displayed lower ratios (Trm-a: 0.055, Trm-b: 0.072, Trm-c: 0.080) (**Fig. 5c, blue**). Analysis of tissue distribution patterns revealed that Dual Expanded cells exhibited notably elevated PA ratios in Teff and Tem populations (**Fig. 5c, red**), suggesting significant enrichment of putative effector-like tumor-associated CD8^+^ T cell clones. The Tumor Multiplet pattern showed elevated PA ratios in Trm-a and Trm-b populations, though at lower levels than Teff and Tem cells. Trm-a and Trm-b cells show lower cytotoxic gene expression compared to Teff and Tem cells^52^, which suggests potentially diminished anti-tumor efficacy by Tumor Multiplet than by Dual Expanded T cells. Notably, the enrichment of PA T cells within Dual Expanded populations supports the idea of recruitment of tumor antigen-specific CD8^+^ T cells from peripheral sites. Consistent with their lack of clonal expansion, Tumor Singleton T cells showed the lowest PA ratios in Teff, Tem, and Trm-c subtypes. Based on these observations, EpitopeGen enables direct computational assessment of PA T cell abundance, providing an efficient alternative to traditional, labor-intensive biological assays.

We next investigated the differential clonal expansion patterns between PA and NA T cells by introducing the Phenotype-Relative Expansion (PRE) ratio, defined as the ratio of average clone sizes between PA and NA TCRs. A PRE ratio of 1.0 indicates equivalent clonal expansion between PA and NA populations. PRE ratios were calculated across cell subtypes and site patterns (**Fig. 5d**). Notably, all T cell subtypes exhibited PRE ratios exceeding 1.0 in the aggregate analysis (**Fig. 5d, All**), consistent with the established principle that antigen-specific CD8^+^ cells undergo preferential clonal expansion after antigen recognition^86^. Although single tumor cells showed uniform PRE ratios of 1.0 by definition, Dual Expanded cells (especially Teff and Tem) showed markedly elevated PRE ratios, suggesting coordinated clonal expansion in both tumor tissue and normal adjacent tissue in response to tumor antigens. Tumor Multiplet cells also showed high PRE ratios across all cell subtypes. Patient-specific analysis revealed substantial heterogeneity in clonal expansion patterns (**Extended Data Fig. 7**). Several patients (Lung Patient 1, Lung Patient 2, Renal Patient 2, and Lung Patient 4) exhibited pronounced PRE ratios in Dual Expanded cells, with distinct distributions across T cell subtypes (**Extended Data Fig. 7a-d**). For example, Lung Patient 1 showed high PRE ratios in Teff, Tem, and Trm-a cells, while Lung Patient 4 showed elevated ratios in Teff and Trm-c cells. In contrast, Renal Patient 1 and Lung Patient 6 showed minimal clonal expansion across all cell subtypes (**Extended Data Fig. 7e,f**). These results reveal that T cell populations vary widely in both their clonal expansion patterns and antigen recognition capabilities, aligned with previous findings by Yost et al.^51^. This quantitative approach to measuring antigen recognition heterogeneity provides a valuable new tool for assessing disease status and stratifying patients.

Differential gene expression analysis between PA and NA CD8^+^ T cell clones revealed distinct transcriptional signatures associated with antigen specificity (**Fig. 5e**). In the aggregate analysis (**Fig. 5e, All**), PA T cells exhibited enhanced cytotoxic potential, characterized by upregulation of genes encoding cytolytic effector molecules (*GZMB*, *PRF1*), TCR signaling components (*ZAP70*, *LAT*, *LCK*), and effector-associated transcription factors (*TBX21*, *EOMES*), while showing decreased expression of multiple exhaustion markers (*TIGIT*, *ENTPD1*, *HAVCR2*, *PDCD1*, *CTLA4*). This molecular signature was similarly observed in Dual Expanded T cells - populations showing clonal expansion in both tumor and normal adjacent tissue - which displayed heightened expression of cytotoxic effector molecules (*GZMB*, *PRF1*, *TNF*) and terminal differentiation markers (*KLRG1*), coupled with reduced expression of multiple exhaustion markers. This transcriptional profile mirrors that of recently activated effector memory/effector T cells (*T*_EMRA_) described by Zhang et al.^87^, which exhibited clonal expansion across blood, normal tissue, and tumor tissue with distinctive tumor-specific TCR repertoires. In contrast, Tumor Multiplet PA T cells, while maintaining high expression of cytolytic molecules (*GZMB*, *PRF1*) and cytokines (*IFNG*), showed a mixed exhaustion phenotype (elevated *LAG3* but downregulated *TIGIT*, *PDCD1*, *CTLA4*) and decreased *TNF* expression. Unlike their Dual Expanded counterparts, these cells lacked upregulation of TCR signaling markers and key transcription factors. The enhanced activation signatures observed in Dual Expanded PA T cells may explain the superior response to immune checkpoint inhibitors in patients with prominent Dual Expanded T cell signatures^52^. These molecular insights validate EpitopeGen’s predictive capabilities and illuminate the transcriptional programs underlying effective anti-tumor T cell responses.

To characterize expanded T cell clones, we visualized the top 100 clonally expanded CD8^+^ T cells per patient, mapping their spatial distribution, clone size, and phenotype (PA TCRs marked by yellow rings). These visualizations revealed different patterns of T cell repertoire in patients. Lung Patient 6 and Renal Patient 2 showed predominantly Dual Expanded PA TCRs, indicating active replenishment of tumor-recognizing T cells from adjacent tissue (**Fig. 5f**, **Extended Data Fig. 8a**). In contrast, the repertoire of Colon Patient 2 consisted mainly of Tumor Singleton and Tumor Multiplet T cells (**Extended Data Fig. 8b**). Lung Patient 3 exhibited both patterns associated with the immunotherapy response: clonally expanded PA T cells in Dual Expanded and Tumor Multiplet patterns, suggesting complementary anti-tumor mechanisms^52^ (**Extended Data Fig. 8c**). Lung Patient 5 showed numerous Dual Expanded and Tumor Multiplet T cells but limited PA T cell expansion (**Extended Data Fig. 8d**). These varying distributions of tumor-specific and bystander T cells align with previous findings^88^. EpitopeGen-based identification of PA and NA T cells enables quantification of T cell activation states within the tumor microenvironment and captures inter-patient heterogeneity, potentially informing personalized immunotherapy strategies^31, 89, 90^.

Sensitivity analyses of EpitopeGen demonstrated the robustness of our findings across various parameter settings. Specifically, our observations from the tumor dataset remained consistent when varying the number of epitopes generated per TCR (*K*) and the relative proportions of antigen categories in the Antigen Category Filter (**Supplementary Note 4**).

### EpitopeGen characterizes COVID19-associated CD8^+^ T cells

EpitopeGen was used to characterize COVID-19-associated CD8^+^ T cells using the data from Su et al.^91^. This study provided paired RNA and TCR sequencing data of blood samples from 139 donors, classified into four groups according to the WHO Ordinal Scale (WOS): healthy controls (WOS=0, no viral infection), mild cases (WOS=1-2, ambulatory with limited symptoms), moderate cases (WOS=3-4, hospitalized, potentially requiring supplemental oxygen), and severe cases (WOS=5-7, hospitalized requiring advanced respiratory support). Our analysis focused on CD8^+^ T cells. Seven major cell clusters were identified and consolidated into four subtypes based on the gene signatures described by Su et al. (**Methods**): naive, memory, effector, and proliferative (**Fig. 6a**). The distribution of these subtypes varied with the severity of the disease. Healthy individuals exhibited a predominance of naive cells, while mild cases showed a more balanced distribution of cell subtypes. As the disease severity increased, the T cell populations shifted, with moderate cases dominated by effector cells and severe cases characterized by a prevalence of proliferative cells. These results indicate a dynamic remodeling of T cell populations in response to increasing COVID-19 severity^91^.

**Figure 6:**
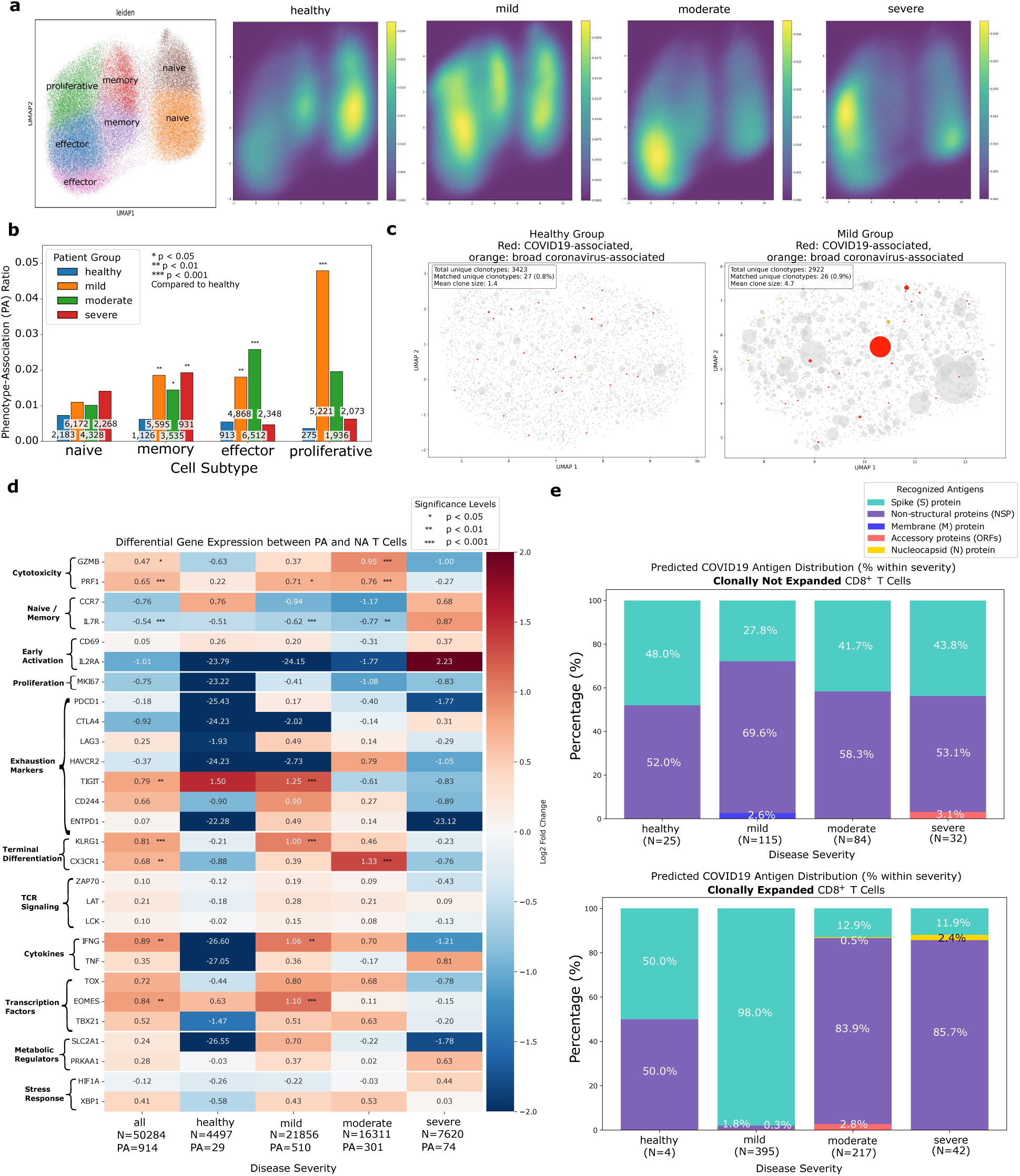
Comparative analysis of Phenotype-Associated (PA) and Not-Associated (NA) TCRs in the paired scRNA and TCR sequencing data measured from COVID-19 patients with varying severity. **a.** UMAP visualization of CD8^+^ T cells based on gene expression data, clustered using the Leiden algorithm (resolution 1.0). Colors represent distinct clusters. Four distinct cell types were identified: naive, memory, effector, and proliferative. Adjacent heatmaps illustrate cluster prevalence across patient groups: healthy, mild, moderate, and severe (from left to right). Heatmaps were generated using Gaussian kernel density estimation. **b.** Proportions of COVID-19-associated TCRs (PA ratio) across patient severity groups (healthy, mild, moderate, severe) and T cell subtypes (naive, memory, effector, proliferative). PA ratio is defined as the number of COVID-19-associated TCRs divided by the total TCR count in each group. One-sided Fisher’s exact tests were performed to compare PA proportions of each severity group to the healthy control within each cell subtype. P-values were adjusted using the Benjamini–Hochberg method (*p *<* 0.05, **p *<* 0.01, ***p *<* 0.001). The sample sizes (n) for both patient groups and cell subtypes are indicated on each corresponding bar. **c.** UMAP visualization of TCR sequences from CD8^+^ T cells in healthy and mild groups. Sequence similarity was calculated using the BLO-SUM62 substitution matrix. Predicted epitope sequences from TCRs, inferred using EpitopeGen, were queried against the curated IEDB database to determine phenotype associations. COVID-19-associated TCRs are shown in red, and broad coronavirus-associated TCRs are in orange. Both plots analyzed the same number of TCRs (4,497). The legend (upper left) displays the total number of clonotypes, COVID-19-associated clonotypes, and the mean clone size. **d.** Heatmap showing differential gene expression between Phenotype-Associated (PA) and Not-Associated (NA) TCRs, grouped by gene functions relevant to CD8^+^ T cells. The analysis was performed across five patient groups: all, healthy, mild, moderate, and severe. Differential gene expression was assessed using the two-sided Wilcoxon rank-sum test from Scanpy, and p-values were adjusted for multiple testing using the Benjamini-Hochberg method (*p *<* 0.05, **p *<* 0.01, ***p *<* 0.001), with log2 fold changes plotted in color scale. **e.** Antigen distribution of Phenotype-Associated (PA) epitopes across patient groups, as identified by EpitopeGen. The stacked bar plots show antigen proportions for COVID-19 antigens (indicated by distinct colors) in two analyses: (top) proportions based on non-expanded CD8^+^ T cells, and (bottom) proportions based on clonally expanded CD8^+^ T cells.

EpitopeGen was used to generate the eight most probable epitopes for each T cell based on its CDR3*β* sequence. T cells were labeled as PA if any of their generated epitopes matched entries in a curated database of COVID-19-associated epitopes; otherwise, they were labeled as NA. For robustness, we took an ensemble of the predictions by three independently trained EpitopeGen models. We first investigated PA ratios by disease severity. The overall frequency of PA T cells was approximately 1% of the total CD8^+^ T cell population (**Fig. 6b**). Healthy individuals exhibited consistently low PA ratios in all cell subtypes, likely due to reduced detection of unexpanded PA cell clones. In contrast, mild cases - patients who could effectively control COVID-19 without hospitalization - demonstrated elevated PA ratios in memory, effector, and proliferative populations. This suggests that the abundance of antigen-specific T cells enables efficient control of COVID-19 as observed in previous studies^92^. Similarly, moderate cases maintained high PA ratios in memory and effector populations. However, severe cases showed markedly reduced PA ratios in both effector and proliferative populations, indicating a deficiency in COVID-19-recognizing effector-like T cells (**Fig. 6b, severe**). This result is consistent with multiple previous studies that observed lymphophenia and insufficient CD8^+^ T cell responses in severe patients^93^.

We investigated the clonal dynamics of PA T cells by performing a comparative analysis of the TCR repertoires between healthy individuals and patients with mild COVID-19. To ensure unbiased comparison, we randomly sampled equal numbers of TCRs from each group. TCR similarities were quantified using the sequence alignment and the BLOSUM62 substitution matrix^94^, followed by UMAP dimensionality reduction. In the resulting visualization, each point represents a unique T cell clone, with point size proportional to the clonal size (**Fig. 6c**). The predicted TCRs to recognize COVID-19-specific epitopes were colored red, while those associated with broader coronavirus epitopes were colored orange to assess potential cross-reactivity. Our analysis revealed different patterns of PA T cell clonal expansion according to disease severity. Although healthy controls showed minimal clonal expansion, mild cases showed pronounced expansion of PA T cells, suggesting robust viral antigen-driven clonal expansion^95, 96^. The average clone size peaked in the mild group, followed by a progressive decrease in the moderate and severe groups. This indicates effective recognition of COVID-19 epitopes and subsequent clonal expansion in mild and moderate patients but not in severe patients. Notably, the severe group showed a smaller average clone size in PA T cells compared to background T cells, suggesting an impaired ability to mount CD8^+^ T cell clonal expansion against SARS-CoV-2 infection (**Extended Data Fig. 9**). In particular, T cells predicted to recognize broader coronavirus epitopes showed a similar severity-dependent expansion pattern, while background (NA) TCRs consistently maintained small average clone sizes. These findings suggest that both COVID-19-specific and cross-reactive T cell responses may contribute to disease progression through differential clonal expansion patterns^97, 98^. These results validate both the robustness and the utility of EpitopeGen in analyzing independent single-cell TCR repertoire data with minimal assumptions.

Next, we analyzed differential gene expression between PA and NA cells by severity group, visualizing log2 fold changes for genes of interest (**Fig. 6d**). In the aggregate analysis, PA T cells exhibited significantly elevated expression of cytolytic molecules (*GZMB*, *PRF1*), migration markers (*CX3CR1*), antiviral cytokines (*IFNG*), and transcription factors (*EOMES*) (**Fig. 6d, All**), indicating a distinct effector phenotype with enhanced cytotoxic characteristic of active antiviral responses. The presence of exhaustion markers suggested chronic antigenic stimulation in these cells. Mild cases maintained this molecular signature, which could explain their effective viral control. This phenotype is similar to clonally expanded and effector-like CD8^+^ T cells observed in acute COVID-19 infection associated with improved clinical outcomes^93, 99^. PA T cells in moderate cases demonstrated a high cytotoxic potential (*GZMB*, *PRF1*) with potentially reduced exhaustion compared to mild cases, along with elevated expression of the chemokine receptor gene *CX3CR1*, suggesting potential migration to inflamed lung tissue. Notably, PA T cells in severe cases lacked characteristic cytotoxic signatures and maintained expression of naive/memory markers, indicating impaired cytotoxic function, T cell dysfunction, or aberrant differentiation. Our observation further supports and details the findings of Su et al.^91^, who showed that patients with severe symptoms exhibited an increase in naive T cell clusters while showing a decrease in activated effector T cells compared to patients with moderate symptoms.

Analysis of the distributions of the generated epitopes revealed distinct antigen recognition patterns across COVID-19 severity groups. Non-expanded T cell clones maintained balanced recognition profiles between structural and non-structural proteins^100, 101^ across all severity groups, suggesting preserved immune surveillance capacity (**Fig. 6e, top**). In contrast, expanded clones showed marked differences between mild cases and moderate/severe cases. Mild cases exhibited substantial spike protein targeting (98% of expanded clones), while moderate and severe cases displayed a strong skew toward non-structural protein recognition (**Fig. 6e, bottom**). This skew could potentially serve as a predictive marker for disease progression, complementing existing findings on T cell exhaustion signatures in severe COVID-19^102, 103^. The higher proportion of spike-specific T cells in mild cases compared to moderate (12.9%) and severe cases (11.9%) highlights the importance of structural protein recognition by CD8^+^ T cells in viral control, not only by CD4^+^ T cells, addressing a gap noted in previous studies^104–106^. These findings suggest that proper clonal expansion of CD8^+^ T cells that recognize the spike protein may be crucial for optimal disease control. Based on these analyses, EpitopeGen enables systematic investigation of antigen recognition patterns from single-cell TCR sequencing data.

## 3 Discussion

Precise identification of T-cell binding partners is crucial for understanding human adaptive immunity and developing targeted therapies, including immunotherapies and personalized vaccines. We introduced EpitopeGen, a large-scale generative transformer designed to predict potential epitope sequences from TCR data, thereby enhancing the utility of single-cell TCR sequencing analysis. To address the fundamental challenge of limited paired TCR-epitope training data, we proposed a semi-supervised learning method that incorporates a large number of unpaired data complemented by a novel Antigen Category Filter algorithm that ensures biologically plausible antigen category distributions in the data for training EpitopeGen.

Our comprehensive evaluation demonstrated the necessity of each component in EpitopeGen’s development. Simply training off-the-shelf transformer models with publicly available data proved inadequate, resulting in highly redundant epitope sequences. Similarly, finetuning existing protein language models failed to capture natural amino acid distributions in the generated epitopes. The inclusion of MHC information, while biologically relevant, reduced epitope diversity due to the limited number of (TCR, epitope, MHC) triplets in the currently available data. These challenges motivated our key innovations: the Robust Affinity Predictor for reliable binding prediction, BINDSEARCH for leveraging unpaired data, and the Antigen Category Filter for maintaining biologically appropriate antigen category distributions. Together, these components enable EpitopeGen to generate epitopes that simultaneously satisfy multiple critical criteria: high binding affinity, sequence diversity, natural amino acid composition, and biophysical stability. Importantly, when applied to repertoire-scale TCR data, EpitopeGen produced epitopes with antigen category distributions aligning with established immunological principles.

Application of EpitopeGen to single-cell paired RNA and TCR sequencing datasets revealed important insights into both cancer and COVID-19 contexts. In cancer, phenotypeassociated (PA) T cells - especially those clonally expanded in both tumor and normal adjacent tissues - showed enhanced cytotoxic markers and decreased exhaustion marker expression compared to non-associated cells. These findings support the hypothesis of continuous peripheral replenishment of tumor-specific T cells^52^. In COVID-19, PA T cells from mild/-moderate patients demonstrated increased clonal expansion with elevated cytotoxic markers compared to healthy individuals, aligning with previously observed activated states^92, 95, 96^. In contrast, PA T cells in severe patients instead showed elevated naive/memory markers, suggesting ineffective viral control in this group^91^. We also identified clonal expansion of CD8^+^ T cells that recognize broader coronavirus antigens, corroborating previous studies on CD4+ T cells^97, 98, 104^.

EpitopeGen’s potential extends beyond the analyzed cancer types and COVID-19, offering promising applications in broader oncological contexts and viral infections. Its utility may encompass autoimmune conditions where CD8^+^ T cells are pivotal, such as Type 1 Diabetes^107^ and Multiple Sclerosis^108^. Furthermore, EpitopeGen could facilitate a deeper exploration of tumor microenvironments, potentially uncovering novel biomarkers to advance immunotherapy development.

Several limitations should be noted. EpitopeGen’s analyses are most reliable in the comparative analysis setting, such as comparing PA ratios across cell types, tracking immune responses, or assessing differences between patient groups. However, the absolute values of PA ratios require careful interpretation due to their dependence on the number of generated epitopes for each TCR, the completeness of the reference database, and TCR cross-reactivity. Although the current availability of TCR-epitope binding data limits EpitopeGen’s potential performance, this limitation also highlights clear avenues for improvement. Future development efforts will incorporate the analysis of class II MHC-presented peptides, enabling the study of CD4^+^ T cells and providing a more comprehensive understanding of the T cell repertoire.

To the best of our knowledge, EpitopeGen is the first tool available for epitope generation from TCR sequences, enabling the analysis of individual patient TCR repertoires. The model, its training data and strategy, as well as the evaluation framework, represent valuable resources for future advancements in this field. We envision EpitopeGen becoming an essential tool for analyzing TCR sequencing data and inspiring future efforts to develop advanced computational methods for TCR repertoire analysis.

## 4 Methods

### Robust Affinity Predictor (RAP) training

Training datasets for binding affinity predictors were compiled from four public sources: VDJdb, IEDB, PIRD, and McPAS-TCR. Data selection criteria included CDR3*β* lengths of 10-20 amino acids and epitope lengths of 8-12 amino acids. Samples containing non-standard amino acids were excluded. For VDJdb, only samples with vdjdb.score *>* 0 were considered. The resulting merged dataset comprised 116,057 TCR-epitope pairs, encompassing 98,460 unique TCRs and 1,141 unique epitopes. Detailed dataset split information was provided in **Supplementary Table 1**.

Robust Affinity Predictor, our binding affinity predictor for pseudo-labeling, was developed by modifying TABR-BERT with three architectural changes: implementing a Softmax layer in the head architecture, removing MHC-related architectures, and utilizing PyTorch’s CrossEntropyLoss instead of Contrastive loss. For the second modification, we re-trained the BERT model^53^ solely on epitope sequences, a process that took two days using two NVIDIA L40S GPUs.

A key challenge in training TCR-epitope binding predictors is the lack of confirmed nonbinding pairs (negative pairs) in public datasets. To overcome this limitation, we developed Triple Negative Sampling (TNS), which generates diverse negative training examples through three complementary strategies. First, we pair known epitopes with TCRs from a large external pool (TCRNegSet, next section). Second, we generate negative samples by pairing known TCRs with epitopes from a large external pool (EpiNegSet, next section). Third, we randomly pair TCRs and epitopes within the dataset, leveraging the assumption that random TCR-epitope pairs are unlikely to bind. This diversified negative sampling approach helps reduce potential biases arising from relying on any single sampling strategy. The final model combined the predictions of five independently trained models through ensemble averaging.

### Large-scale unpaired TCR and epitope datasets

Pseudo-labeling involves large-scale inference on randomly paired TCRs and epitopes using the Robust Affinity Predictor, selecting high-affinity samples for model training. We used the TCRdb dataset, which contained 19,701 TCR sequencing measurements as of May 2024, as the unpaired TCR sequence pool. TCRs with lengths between 10-20 amino acids and comprising only standard amino acids were selected, yielding 7,331,478 unique CDR3*β* sequences. These TCR CDR3*β* sequences were split into TCRCandidateSet (6,831,478) for pseudo-labeling and TCRNegSet (500,000) for Triple Negative Sampling.

Epitope sequences were obtained from NetMHCPanv4.0 (Binding Affinity and Eluted Ligands datasets), MHCflurryv2.0 (S1-S5 datasets), and SysteMHC (per-MHC datasets from https://systemhc.sjtu.edu.cn/download). Epitopes of 8-12 amino acid length, containing only standard amino acids, were included, resulting in 21,801,187 unique sequences. When training EpitopeGenMHC, only MHC-epitope binding pairs were extracted, with the most likely MHC selected using the mhcflurry2.ba best allele column. Inhibitory Concentration 50% (IC50) units were converted to binary labels using the formula `lambda x: 1 - log(max(x, 0.1), 50000)`^109^, with a threshold of 0.42 defining a hit. The epitope dataset was split into EpiCandidateSet (20,000,000) for pseudo-labeling and EpiNegSet (1,801,187) for Triple Negative Sampling.

### Semi-supervised learning method

The BINDSEARCH algorithm (**Algorithm 1**) generates a pseudo-labeled dataset of TCR-epitope pairs. Given sets of unpaired TCR sequences 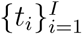 and epitope sequences 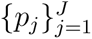, the algorithm uses a binding affinity predictor function *R* (implemented as RAP) to estimate the binding affinity between TCRs and epitopes (*I* = 6, 831, 478*, J* = 20, 000, 000 were used). For each unpaired TCR *t_i_*, the algorithm randomly samples *β* = 10, 000 candidate epitopes from 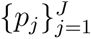. The binding affinity *a* is computed for each TCR-epitope pair (*t_i_, p*) using *R*. Subsequently, the top *n*_max___tcr_ = 32 pairs with the highest binding affinities are retained for each TCR. To mitigate redundancy in the resulting dataset, a filtering step is applied. Epitopes that occur more than *n*_max___epi_ = 100 times in all pairs of TCR-epitopes are excluded. The value of *n*_max___epi_ was determined based on the ratio of TCR to epitope observed in public datasets (specifically 116,057 epitopes to 1,141 TCR, resulting in a ratio of approximately 102). This process resulted in a pseudo-labeled dataset comprising |*D*|= 16, 909, 219 TCR-epitope pairs. The algorithm took four days to run on ten NVIDIA L40S GPUs.

#### Algorithm 1 BINDSEARCH

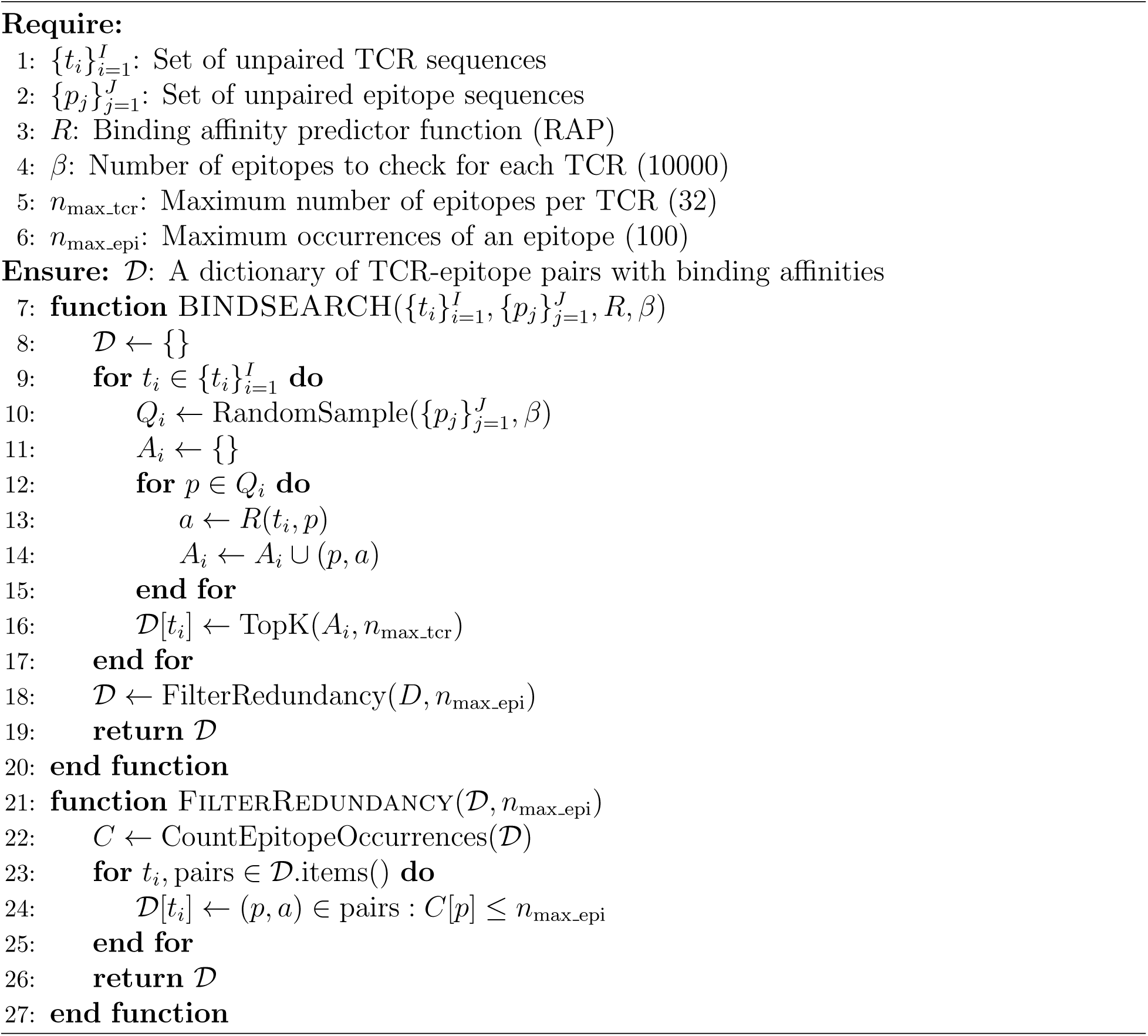

#### Algorithm 2 Antigen Category Filter (ACF)

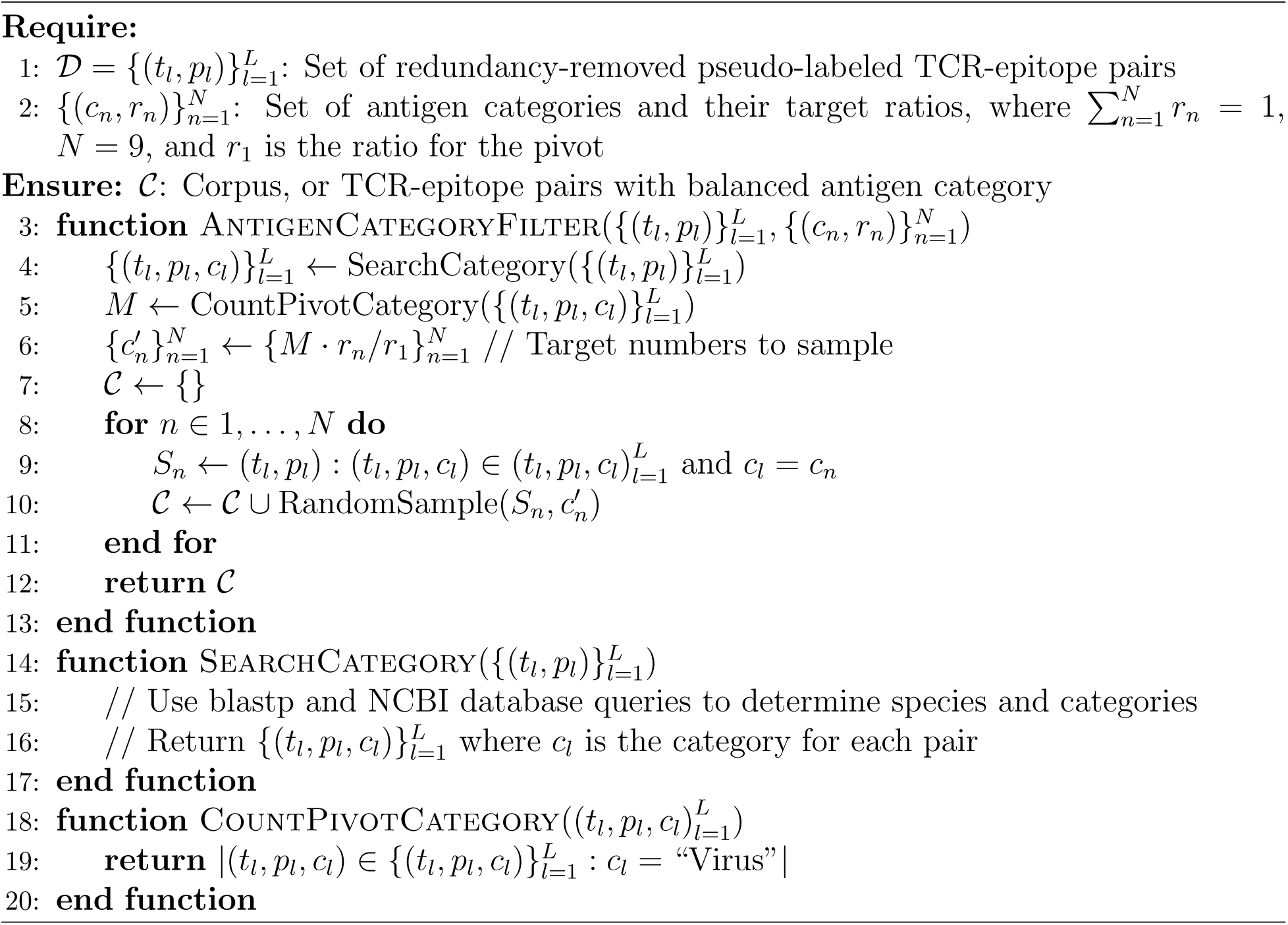

### Antigen Category Filter

The intermediate dataset D was biased towards Eukaryotic species, a consequence of the peptide collection methods used by NetMHCPanv4.0, MHCflurryv2.0, and SysteMHC. This bias likely reflects research priorities and funding rather than the biological distribution of antigens potentially recognized by CD8^+^ T cells. To correct this discrepancy, we implemented the Antigen Category Filter (ACF) algorithm (**Algorithm 2**) to impose a biologically relevant epitope distribution. ACF takes as input a set of redundancy-removed pseudo-labeled TCR-epitope pairs 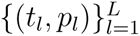 and a set of antigen categories with their target ratios 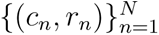.

To determine the target antigen ratios, we considered five immunological insights: (1) Viral Dominance: Studies indicate that a substantial proportion of CD8^+^ T cells recognize viral antigens, with Masopust et al.^33^ showing approximately 80% of splenic CD8^+^ T cells recognize viral epitopes during peak infection. We set the proportion of viral antigens to be P(virus) ≥ 50%. (2) Limited Bacteria: Although certain bacterial species such as *Listeria monocytogenes* and *Mycobacterium tuberculosis* can elicit CD8^+^ T cell responses through intracellular infection^41^, their contribution is limited compared to viruses. We set bacterial antigens to P(bacteria) ≤ P(virus) × 0.2. (3) Endogenous Presence: Due to thymic negative selection^39^, CD8^+^ T cells that recognize selfor tumor antigens are less frequent than virus-specific T cells. Rizzuto et al.^37^ confirmed the significantly lower frequency of self/tumor antigen-specific T cells than T cells specific for foreign antigens. We therefore constrained the self-antigen ratio in the range of 0.03-0.15 and the tumor antigen ratio in the range of 0.01-0.05. (4) Rare Fungi and Parasites: Although some fungi like *Histoplasma capsulatum*^42^ can trigger CD8^+^ T cell responses, their frequency is considerably lower, with only a few dozen fungal species causing regular human infections^43^. This led us to restrict their combined proportion to P(fungi + parasites) ≤ P(virus) × 0.1. (5) No Reported Pathogenic Archaea: Given the absence of documented archaeal pathogens causing human disease^47, 48^, we kept the proportion of archaeal antigens in our dataset very low (P(archaea)≤0.01). The target proportions for each category were sampled from their respective ranges and normalized to ensure that their sum equaled one, while maintaining the defined biological constraints. More discussion of target ratio ranges was provided in **Supplementary Note 2**.

The Antigen Category Filter algorithm begins with a species identification process for each epitope, conducted using SwissProtDB through `blastp` queries (`-task blastp-short -evalue 20000 -max_target_steps 10`), accounting for the short epitope lengths. This step is encapsulated in the `SearchCategory` function. Subsequently, the NCBI accession numbers were utilized to query proteins and species using the command: `blastdbcmd -db ncbi-blast-2.15.0+/db/swissprot -entry batch output.txt -outfmt “%a %t“`. Species lineage determination involved searching for TaxId from NCBI accession numbers and then querying lineage information from TaxId using `epost`, `esummary`, `xtract`, and `efetch` commands. The resulting XML files were parsed to categorize the species into categories. All viral epitopes were retained as pivot species (counted by the `CountPivotCategory` function).

### EpitopeGen architecture and training

EpitopeGen is a decoder-only transformer model specifically designed to generate epitope sequences while adhering to specific distributional constraints, including epitope diversity and biologically plausible antigen distributions. The model architecture is based on GPT-2, with amino acid sequences encoded using the BPETokenizer^110^ from the Hugging Face^111^ library. This tokenization method allows for the representation of single amino acids or groups of amino acids as individual tokens, potentially capturing meaningful biological motifs. The Byte-Pair Encoding (BPE) algorithm iteratively merges the most frequent pairs of tokens, capturing the recurring subsequences as single tokens. Let *V* be the vocabulary of tokens, with |*V*|= 400, which was found to efficiently encode all sequences with the optimal usage ratio (**Supplementary Note 5**). The BPETokenizer was trained using Corpus C. For a TCR-epitope pair (*t, p*), the input sequence is tokenized as:

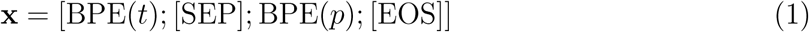

where BPE(·) denotes the BPE tokenization function, ‘;’ represents concatenation, [SEP] delineates the boundary between TCR and epitope sequences, and [EOS] marks the end of each sequence. The tokenized sequences were processed using positional embeddings, where position-specific vectors are added to the token embeddings to maintain sequence order information.

We adopted the GPT2-small architecture^28^, a decoder-only transformer architecture introduced by Vaswani et al.^112^. The model consists of 12 transformer decoder layers, each containing a multi-head self-attention mechanism followed by a position-wise feedforward network. The self-attention mechanism in each layer has 12 attention heads, allowing the model to attend to different aspects of the input sequence simultaneously. Each head projects the input into query, key, and value spaces of dimension 64 (768/12, where 768 is the dimensionality of each token embedding), allowing the model to capture various types of sequential dependencies in TCRs. The position-wise feedforward network in each layer consists of two linear transformations with a Gaussian Error Linear Unit (GELU) activation function, expanding the intermediate representation to dimension 3,072 before projecting back to 768. This architecture, which contains 124 million parameters in total, was implemented using the HuggingFace and PyTorch^113^ frameworks. The model defines a probability distribution *p****_θ_***(**x**) over a sequence of tokens **x** of length *n*, which can be factorized as:

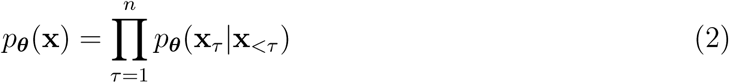

where ***θ*** represents the model parameters, **x***_τ_* is the *τ* -th token in the sequence, and **x***_<τ_* denotes all tokens before *τ* . This autoregressive formulation allows the model to generate epitope sequences token by token, conditioned on the input TCR sequence. The objective function for training is the negative log-likelihood:

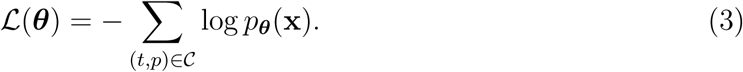

Given the narrow length distribution of the TCR and epitope sequences compared to natural language paragraphs, each batch contained a single TCR-epitope pair. The AdamW optimizer^114^ was used with parameters: initial learning rate (*α* = 1 × 10^−5^), *β*_1_ = 0.9, *β*_2_ = 0.999, *ɛ* = 1 × 10^−8^, and weight decay (*λ* = 0.01). Training took four hours for EpitopeGen and four days for EpitopeGenNoACF using four NVIDIA L40S GPUs. Additional details on EpitopeGen training can be found in **Supplementary Note 5**.

### Metrics to evaluate generated epitopes

To assess whether the predicted epitope distributions align with domain knowledge of CD8^+^ T cell recognition, we developed the Biological Sanity Score (BSS). This score quantifies the similarity between a predicted distribution *P* and a reference distribution *R*, weighted by the biological importance of different antigen categories:

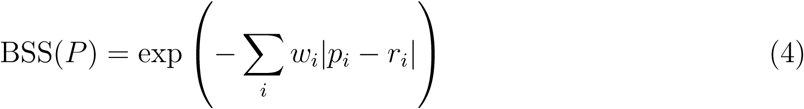

where *p_i_* and *r_i_* are the proportions of antigen category *i* in the predicted and reference distributions, respectively, and *w_i_* is the weight. The weights were given by: virus (5.0), tumor (3.0), bacteria (2.0), self-antigens (2.0), parasites (1.0), fungi (0.5), archaea (0.2), and others (0.3). The reference distribution was obtained from a healthy donor (10x dataset): virus (45%), tumor (17.5%), bacteria (12.5%), self-antigens (10%), parasites (6.5%), fungi (3%), archaea (0.5%), and others (5%). The score ranges from 0 to 1, with 1 indicating perfect alignment.

We employed multiple metrics to evaluate epitope diversity. Shannon Diversity quantifies distribution entropy, considering both unique epitopes and their abundances. Rényi and Simpson’s Diversity, which incorporate quadratic terms, reduce sensitivity to rare epitopes. The epi-to-TCR ratio is calculated as the number of unique generated epitopes divided by unique input TCRs. We also measured the average repetition frequency of top 1% epitopes and the proportion of epitopes within the most frequent 10% to assess redundancy.

Chemical feasibility was assessed using the `ProtParam` package^74^. We analyzed extinction coefficient to evaluate light-absorbing properties, aromaticity for hydrophobicity and potential TCR interactions, and secondary structure for folding tendencies. Additionally, we examined isoelectric point to understand pH-dependent charge behavior and instability index to assess stability in solution.

### Molecular Dynamics (MD) simulation

MD simulation provides ways to evaluate the generated epitopes, which is orthogonal to measuring binding affinities using Robust Affinity Predictor, a deep learning-based predictor. After generating an epitope, the first step was to predict the 3D structure of the full TCR-pMHC complex. For this task, we employed TCRmodel2, which is based on AlphaFold2^79^. The model inputs comprised the full TCR alpha chain, the TCR beta chain, the epitope, and the full MHC sequence, with MHC alleles mapped to sequences referring to the IMGT/HLA Database^115^. The TCR alpha chain was fixed, while the TCR beta chain’s CDR3*β* region was replaced with the input CDR3*β* sequence. Of the five PDB files generated by TCRmodel2, only `pdb_0` was used for subsequent analyses. After predicting the 3D structure, structure refinement and interface analysis were performed using Rosetta 3.14 (released March 7, 2024). The Rosetta `relax` application (`main/source/bin/relax.static.linuxgccrelease`) was executed with options`-relax:default repeats 5 -nstruct 1` for structure relaxation. Then, interface analysis was performed using Rosetta’s InterfaceAnalyzer (`main/source/bin/InterfaceAnalyzer. static.linuxgccrelease`) to examine protein-protein interface properties.

### Wu et al. dataset preprocessing

The dataset from Wu et al.^52^ was obtained from Gene Expression Omnibus (GEO) with access number GSE139555, containing clustering, cell type annotation, and TCR contig annotation information. We focused on cells from clusters 8.1-Teff, 8.2-Tem, 8.3a-Trm, 8.3b-Trm, epiand 8.3c-Trm, re-labeled as Teff, Tem, Trm-a, Trm-b, and Trm-c for brevity. The dataset comprised samples from 14 donors in four types of cancer (lung, renal, endometrial, and colorectal), collected from tumor tissue, normal adjacent tissue (NAT), and blood whenever available. The preprocessed data, which contain highly variable genes, UMAP coordinates, cell subtypes, and clonotype information, were initially loaded. To integrate the RNA and TCR sequencing data, we used the scirpy package^116^. We read the TCR data from files with the `contig_annotations.csv` suffix using `scirpy`’s `ir.io.read 10 vdj()` function. The resulting information was stored in a MultiModalData (MuData) object, combining both RNA and TCR data. For TCR analysis, we performed quality control using `scirpy`’s `ir.pp.index chains()` and `ir.tl.chain qc()` functions. We retained cells with chain pairings classified as ‘single pair’, ‘orphan VDJ’, ‘extra VJ’, or ‘extra VDJ’. In cases of ‘extra VDJ’, we selected the first entry. Only productive TRB chains were included for further analysis, as non-productive chains are excluded by `scirpy`. For each cell, we extracted the TRB locus, amino acid junction (CDR3 aa), and nucleotide junction (CDR3 nt) sequences. This information was then merged with the gene expression data based on cell identifiers, resulting in a dataset of 36,986 cells with matched gene expression and TCR information. This dataset contained 9,166 unique CDR3 aa sequences and 9,284 unique CDR3 nt sequences. To analyze the expression levels of genes of interest, we utilized the raw dataset of `barcodes.tsv`, `genes.tsv`, and `matrix.mtx` files, which were loaded using `scanpy`’s^117^ `sc.read 10x mtx()` function. We obtained the curated list of genes associated with CD8^+^ T cells from Wu et al., including exhaustion markers (e.g., *PDCD1*, *CTLA4*, *LAG3*), activation and proliferation markers (e.g., *CD69*, *IL2RA*, *MKI67*), TCR pathway-related genes (e.g., *ZAP70*, *LAT*, *LCK*), cytokines (e.g., *IFNG*, *TNF*), and various other markers related to T cell function and differentiation. The data were normalized to a total count of 10^4^ per cell and log-transformed. While the clones were defined based on CDR3 nt, EpitopeGen used CDR3 aa to generate epitope sequences. These epitopes were then queried against a combined database derived from the Immune Epitope Database (IEDB)^55^ and The Cancer Immunome Atlas (TCIA)^118^, focusing on cancer-related linear epitopes restricted by MHC Class I. The epitopes found in these databases were classified as Phenotype-Associated (PA). Finally, the site patterns of the TCRs were annotated on the basis of their presence in different tissue sites. Statistical analyses, including comparisons between groups, were performed using two-sided Wilcoxon rank-sum test (`scanpy`^117^). Additional details on processing Wu et al. dataset can be found in **Supplementary Note 6**.

### Su et al. dataset preprocessing

For the analysis of COVID-19-related T-cell responses, we utilized the dataset from Su et al.^91^, which comprises paired single-cell RNA and TCR sequencing data from 139 donors. We processed the data by first combining all the files with the format `heathlab_dc_9_17_pbmc_gex_library_XX_X.txt` to create a comprehensive `AnnData` object indexed by cell barcodes. TCR information was extracted from files formatted as `heathlab_dc_9_17_pbmc_cd8_tcr_library_XX_X.txt` and used to filter the `AnnData` object, retaining only cells with associated TCR data. The resulting dataset contained 50,284 unique cells, containing 26,170 unique CDR3 amino acid sequences and 26,621 unique CDR3 nucleotide sequences. We annotated the data with patient demographics and WHO Ordinal Scale scores, mapping the latter to four categories: healthy (0), mild (1 or 2), moderate (3 or 4), and severe (5, 6, or 7). For cell type annotation, we performed standard single-cell RNA sequencing analysis, including normalization, log transformation, selection of 1000 highly variable genes supplemented with the signature genes of CD8^+^ T cells, PCA dimensionality reduction (50 components), Leiden clustering (resolution 1.0), and UMAP visualization. This process identified seven major clusters, which we grouped into four subtypes based on selectively high expression of marker genes: naive (*TCF7*, *LEF1*, *SELL*, *CCR7*), memory (*GZMK*), effector (*NKG7*, *CCL4*, *CST7*, *PRF1*, *GZMA*, *GZMB*), and proliferating (proliferation markers such as *MKI67*, *TYMS*). To identify COVID-19-associated epitopes, we constructed a database using IEDB, querying for linear sequence epitopes from SARS-CoV-2, with positive assays, MHC Class I restriction, and association of COVID-19 disease, resulting in 1,357 unique epitopes. We then used EpitopeGen to generate epitopes based on the CDR3 amino acid sequences. The matches were defined by a Levenshtein distance ≤ 1 with the first match selected in the cases of multiple matches. This preprocessing pipeline allowed us to integrate transcriptomic, clonotypic, and epitope-specific information.Additional details on the processing of the Su et al. dataset can be found in **Supplementary Note 7**.

## Acknowledgements

This work was supported by a Discovery grant from the Natural Sciences and Engineering Research Council (NSERC) of Canada, and a department startup fund from the University of British Columbia (to J.D.). J.D. is a Canada Research Chair and is supported by the Canadian Institutes of Health Research through the Canada Research Chair Program. The computational resource is partially supported by the Canada Foundation for Innovation & John. R. Evans Leader Fund (to J.D.). This research was supported in part through the computational resources and services provided by Advanced Research Computing at the University of British Columbia.

## Author’s contributions

M.M. developed and implemented the model, conducted experiments, and prepared the manuscript with input from J.D. W.T. and C.V. assisted with data analysis. J.D. supervised the project. All authors reviewed and approved the manuscript.

## 5 Extended Data Figure

**Extended Data Fig. 1:**
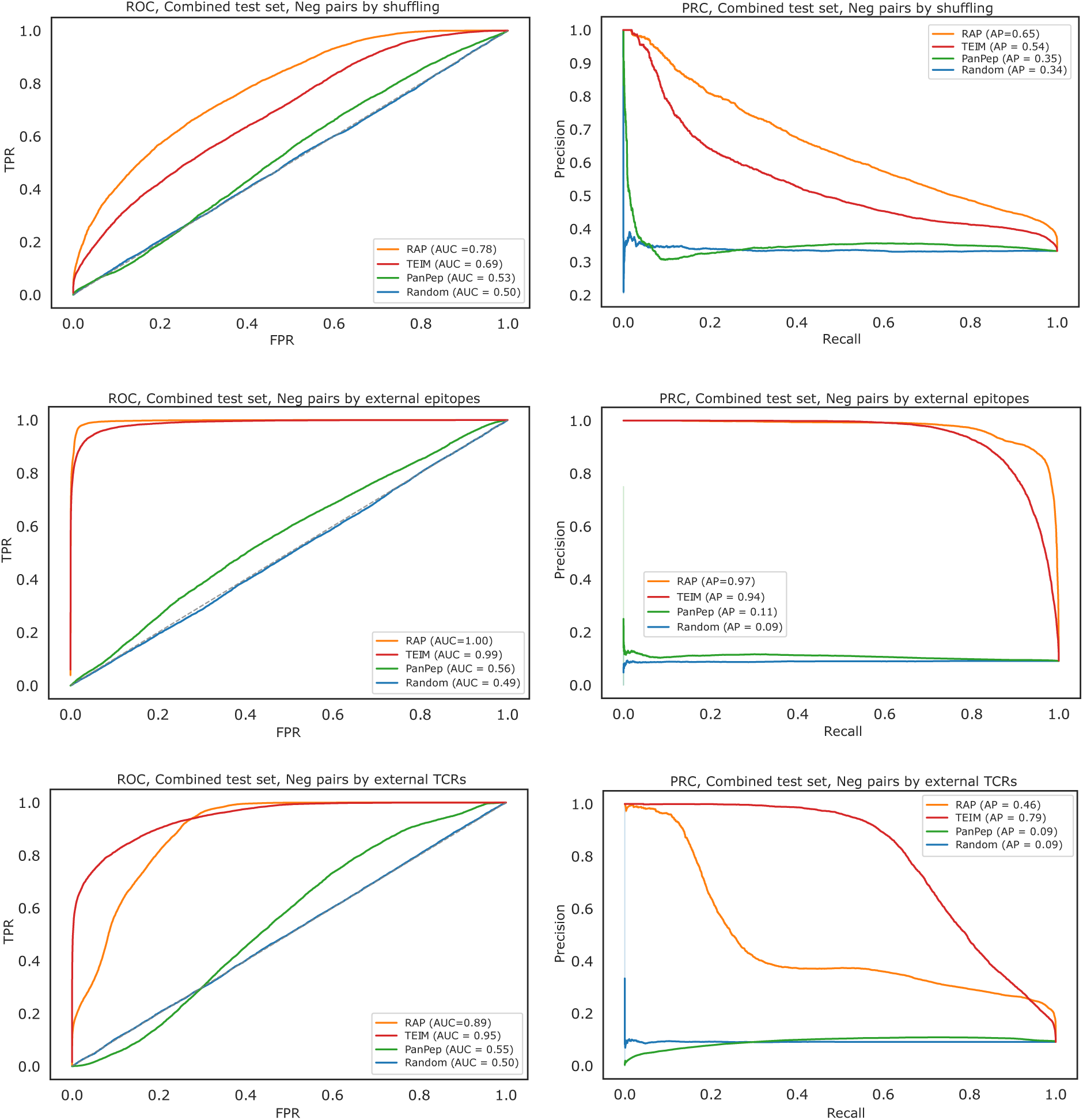
Receiver Operating Curves (ROC) and Precision-Recall Curves (PRC) of Robust Affinity Predictor (RAP) and other competing binding affinity predictors. ROC and PRC of the binding affinity predictors on the combined public test set that comes from four sources: VDJdb, IEDB, PIRD, and McPAS-TCR. Three scenarios of negative sample generation were considered: randomly pairing TCR and epitope in the test set (top), pairing the TCRs with external epitopes (middle), and pairing the epitopes with external TCRs (bottom). Four models were compared: Robust Affinity Predictor (RAP), TEIM, PanPep, and Random. AUC or AP values are denoted in the legends next to the model names.

**Extended Data Fig. 2:**
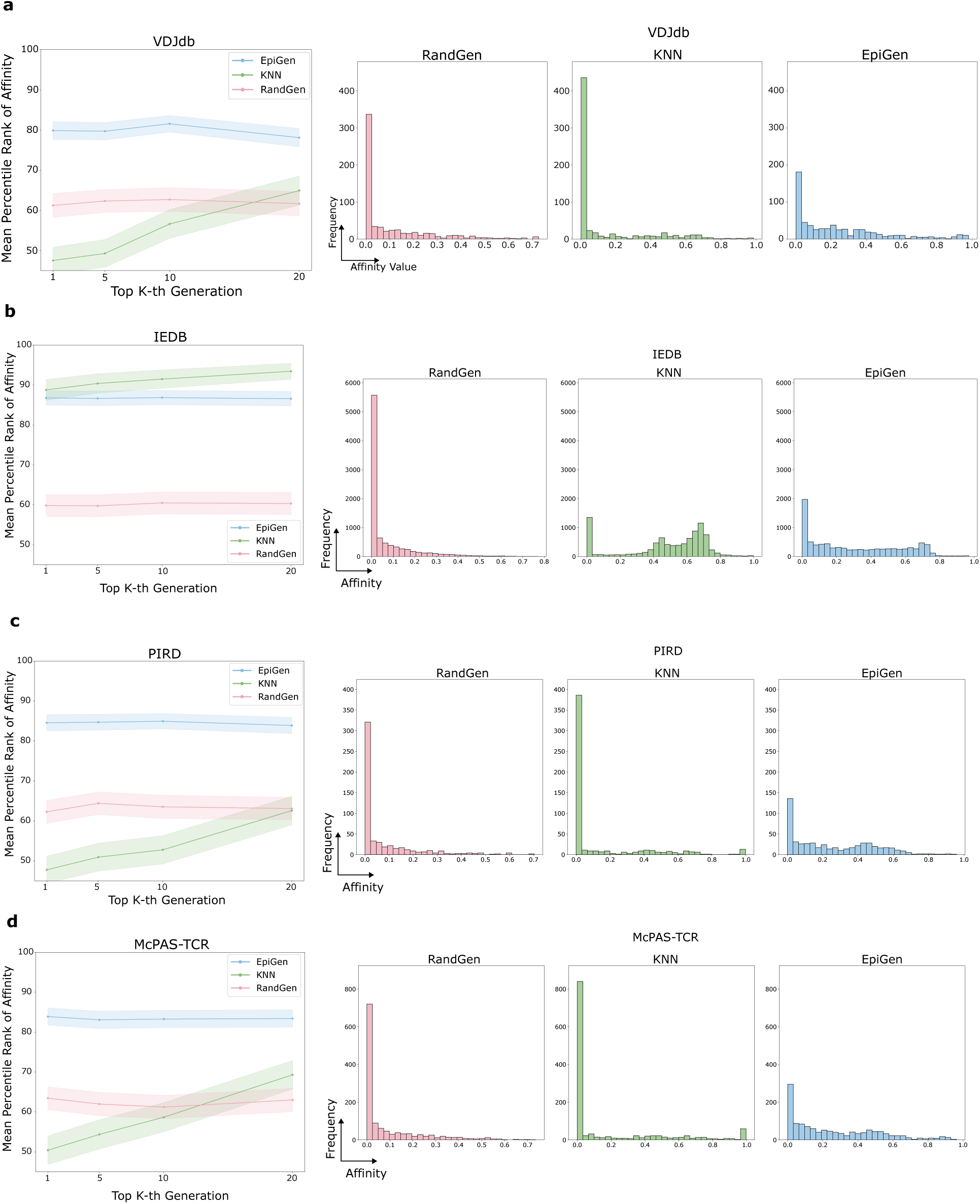
Validation of high binding affinities between EpitopeGengenerated epitopes and input TCRs. **a–d.** The validation results by the test sets: VDJdb, IEDB, PIRD, and McPAS-TCR. The plots show the mean percentile rank of the binding affinities between the top K generated epitopes and TCR_4_s_7_(left) and the predicted binding affinity distributions (right). Three methods were compared: RandGen, KNN, and EpitopeGen. Higher binding affinity values are favored.

**Extended Data Fig. 3:**
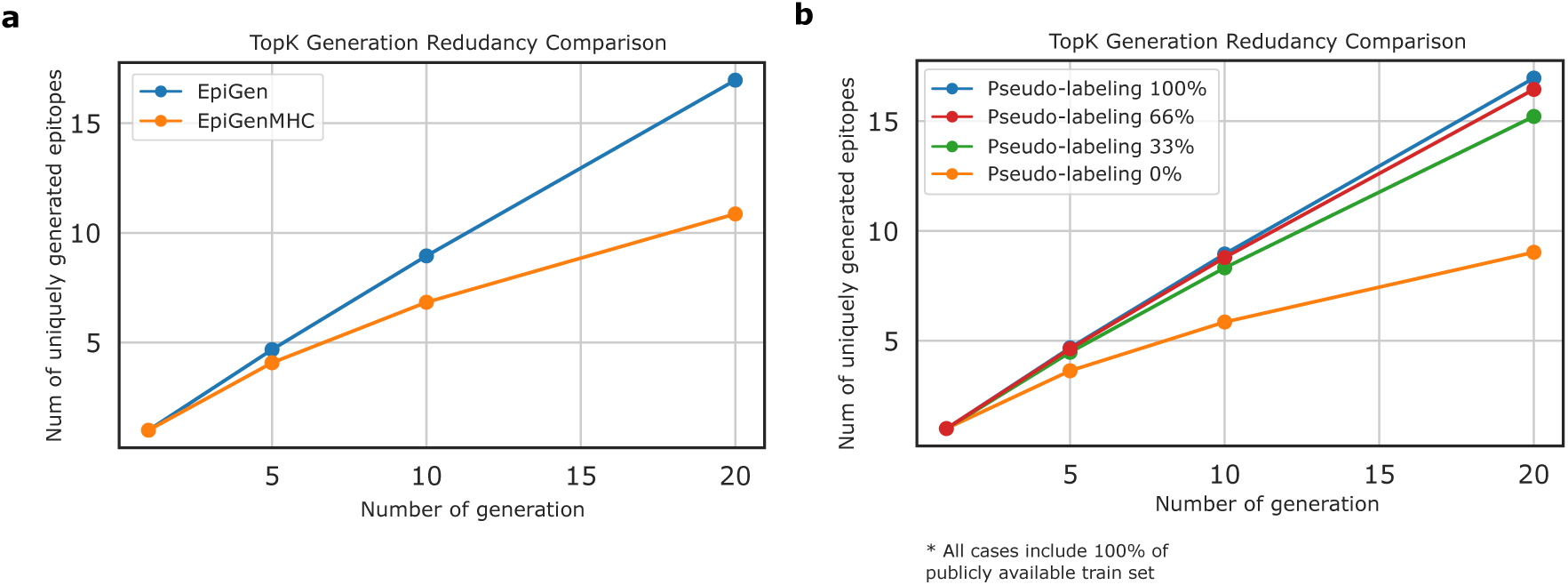
Comparison of redundancy in multiple generations on repertoire-level dataset. **a.** Plot showing the average number of uniquely generated epitopes (y-axis) by the number of generation attempt for each TCR (x-axis). The top generations may produce the same epitope sequences, leading to redundancy. The plot highlights the redundancy of EpitopeGenMHC. The dataset was the 10x dataset. **b.** Plot showing the average number of uniquely generated epitopes (y-axis) by the number of generation attempt for each TCR (x-axis). The plot compares four models trained using different amount of pseudo-labeled dataset from 0%, 33%, 66% to 100%. The dataset was the 10x dataset.

**Extended Data Fig. 4:**
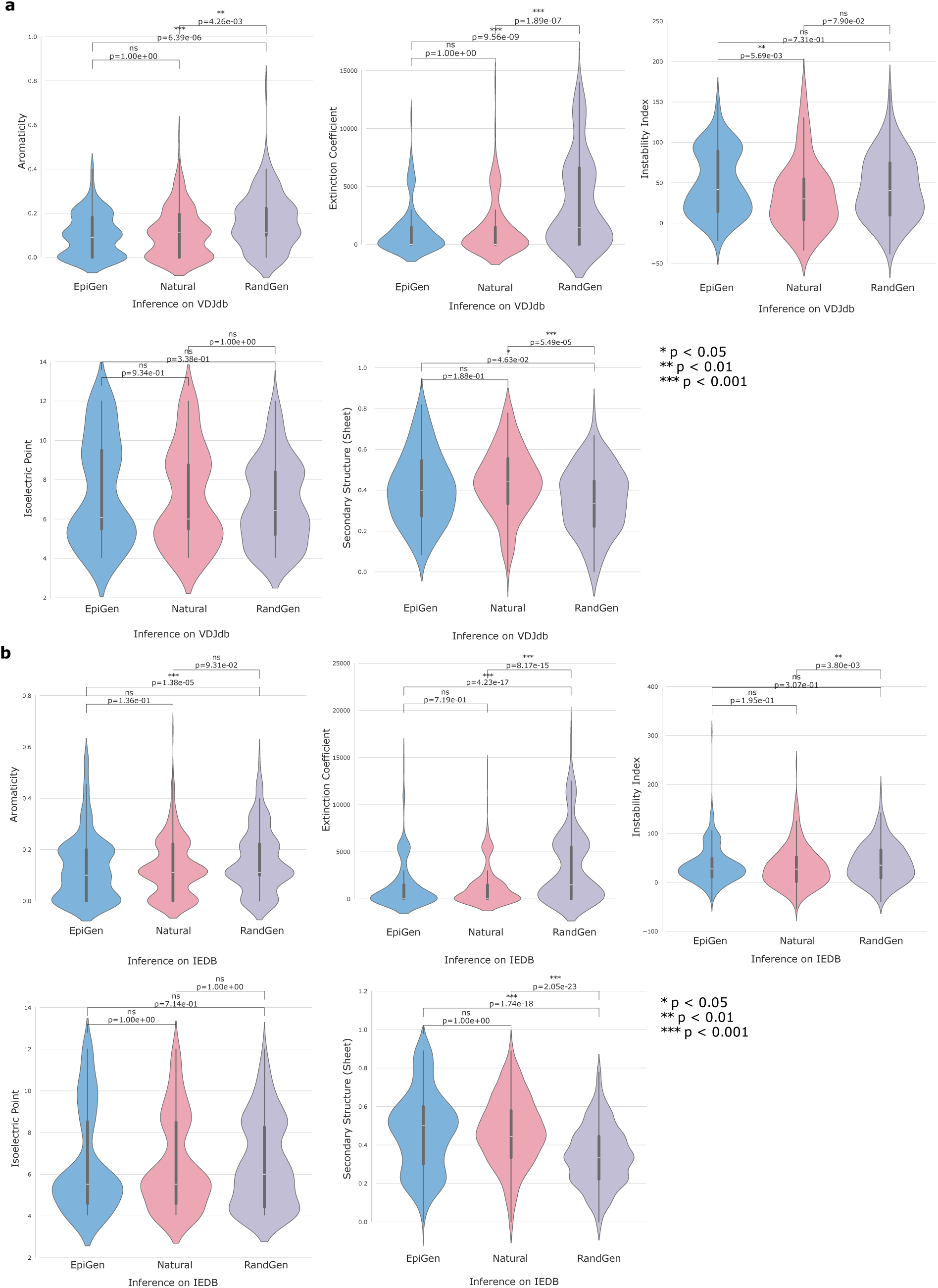
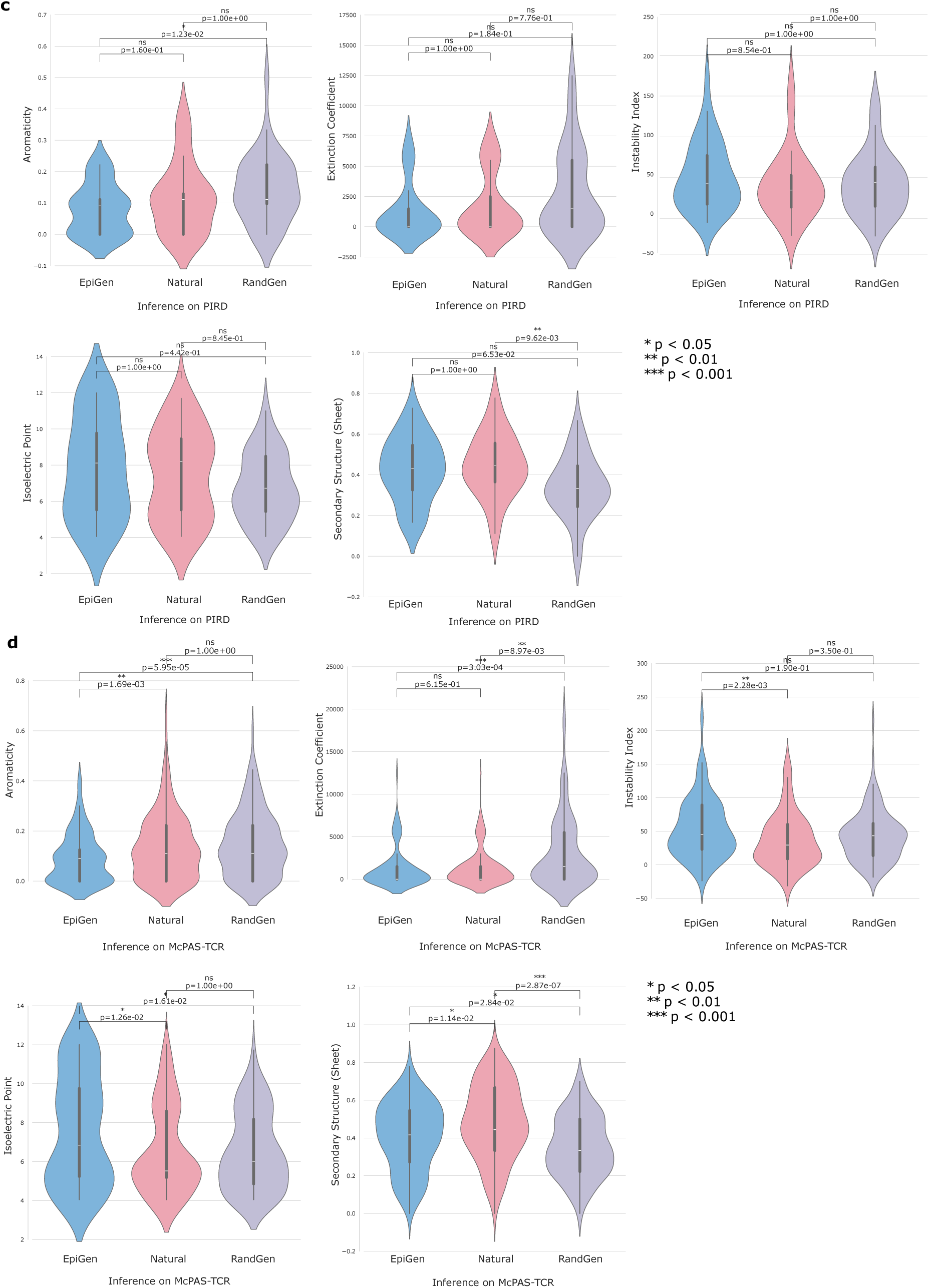
Chemical property distributions comparison. **a–d.** Distribution of physicochemical properties across epitope sources, visualized using violin plots. Five properties are analyzed: Aromaticity, Extinction Coefficient, Instability Index, Isoelectric Point, and Secondary Structure (*β*-sheet content). Epitopes are categorized by source: EpitopeGen-generated (blue), naturally occurring in the test sets (red), and randomly generated (RandGen, red). The analysis spans four test sets: VDJdb (n=642), PIRD (n=545), IEDB (n=9,030), and McPAS-TCR (n=1,276), shown in panels **a**, **b**, **c**, and **d**, respectively. Violin plots incorporate box plots showing the median, interquartile range (IQR), and whiskers (1.5 × IQR). Statistical significance was determined using two-sided Mann-Whitney U tests with Bonferroni correction (*p *<* 0.05, **p *<* 0.01, ***p *<* 0.001, ns: not significant).

**Extended Data Fig. 5:**
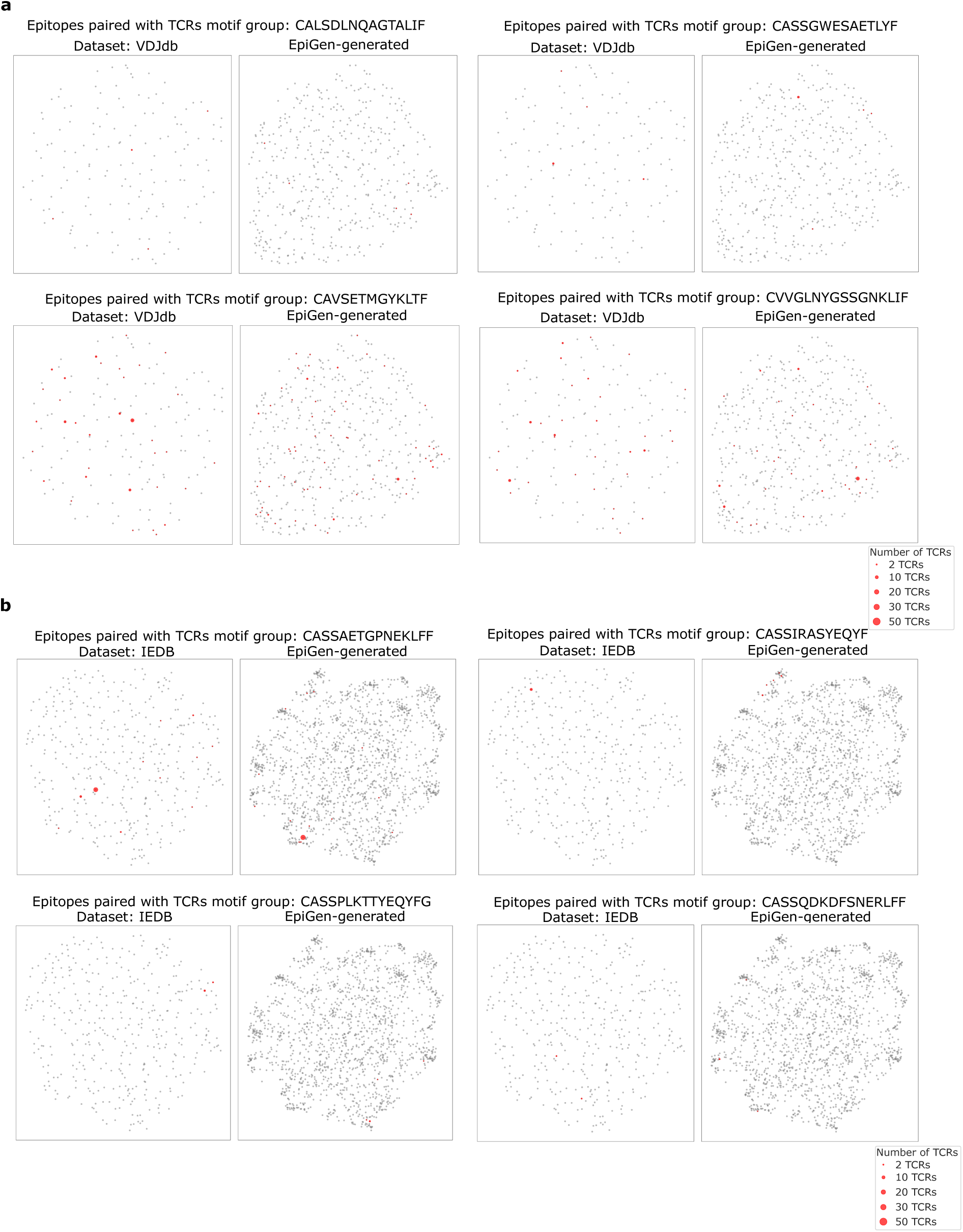
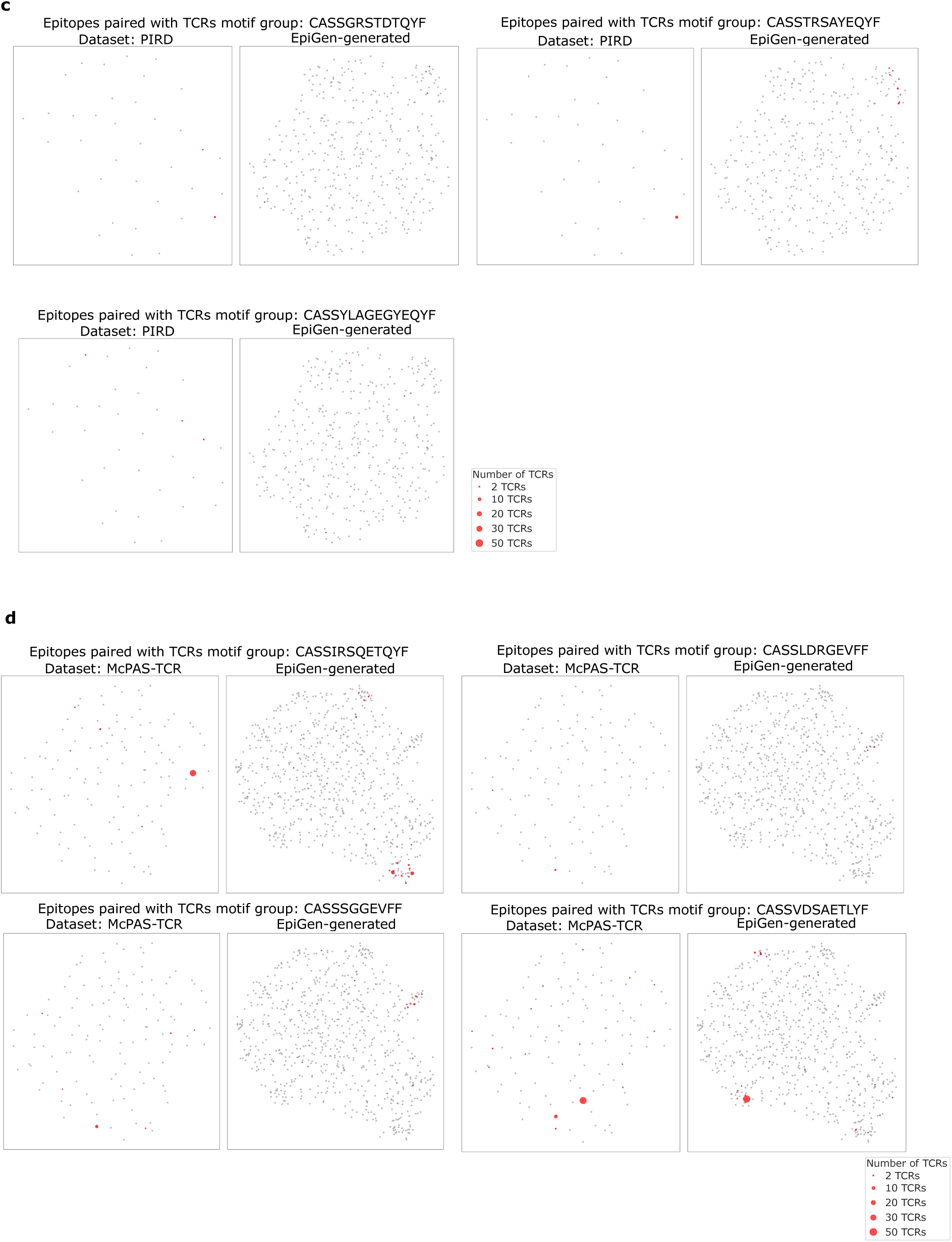
Similar epitopes generated by EpitopeGen tend to be recognized by similar TCRs. **a–d.** UMAP visualization of epitope sequence similarity spaces across four datasets: **a.** VDJdb, **b.** IEDB, **c.** PIRD, and **d.** McPAS-TCR. Sequence similarities were computed using the BLOSUM62 substitution matrix. Each point represents an epitope, with point size proportional to its paired TCR count. Epitopes recognized by TCRs sharing common motifs (identified by GLIPH2 analysis of test sets) are highlighted in distinct colors, while other epitopes are shown in grey. Three to four representative TCR motif groups are displayed per dataset. The visualization demonstrates both the broad distribution of generated epitopes and the clustering of similar epitopes recognized by TCRs with shared motifs.

**Extended Data Fig. 6:**
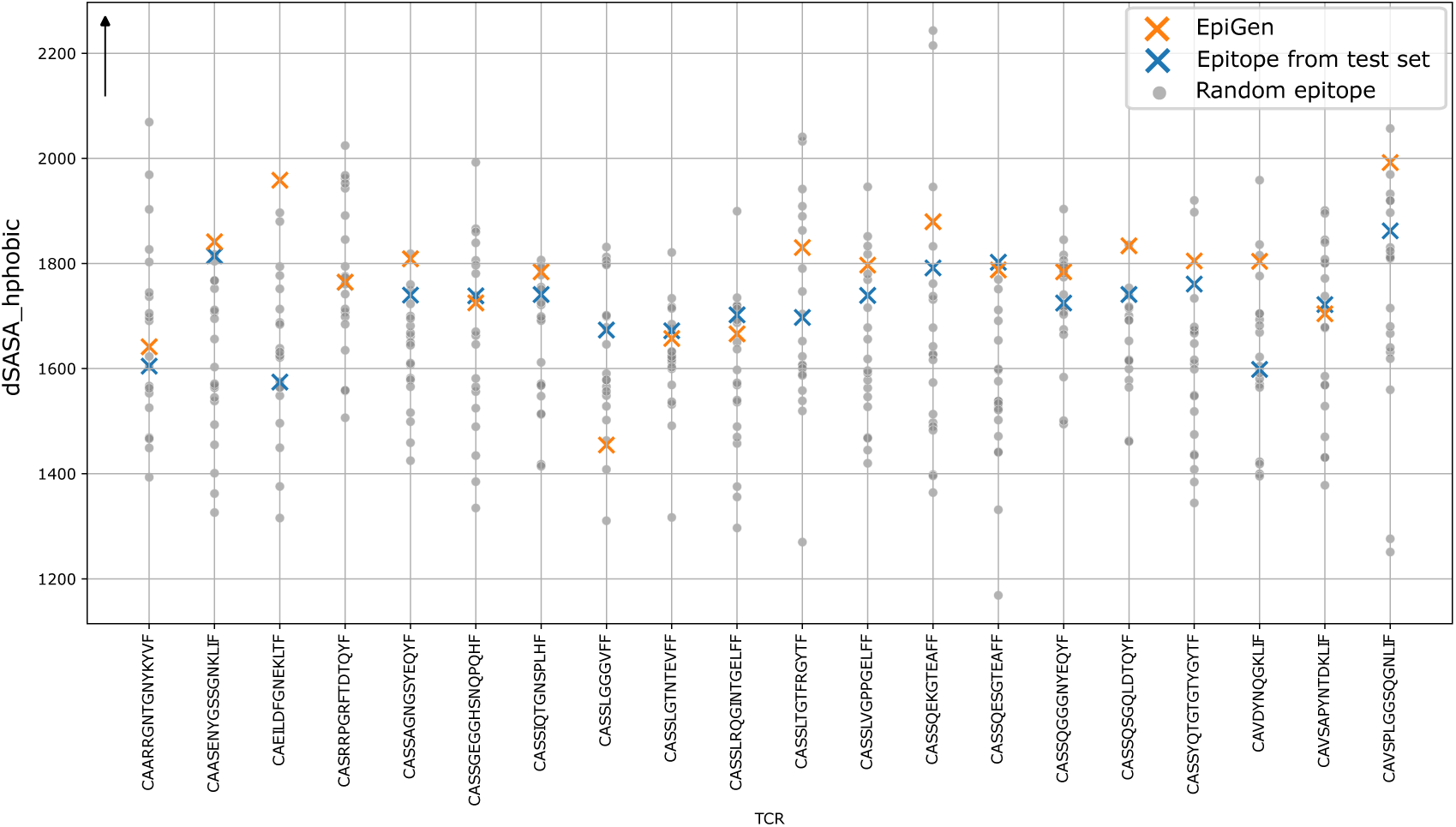
Structural analysis using InterfaceAnalyzer with the generated epitopes. Hydrophobic interaction analysis of TCR-pMHC complexes. Scatter plot showing dSASA hphobic values (y-axis) for complexes containing EpitopeGen-generated epitopes (orange) or natural epitopes from the VDJdb test set (blue). Analysis conducted across 20 TCRs (x-axis), with 20 random epitopes per TCR serving as controls (grey). dSASA hphobic measures the change in hydrophobic surface area that are hidden inside where dSASA stands for ‘delta Solvent Accessible Surface Area’. Higher dSASA hphobic values indicate higher structural stability. dSASA hphobic was calculated using Rosetta’s InterfaceAnalyzer.

**Extended Data Fig. 7:**
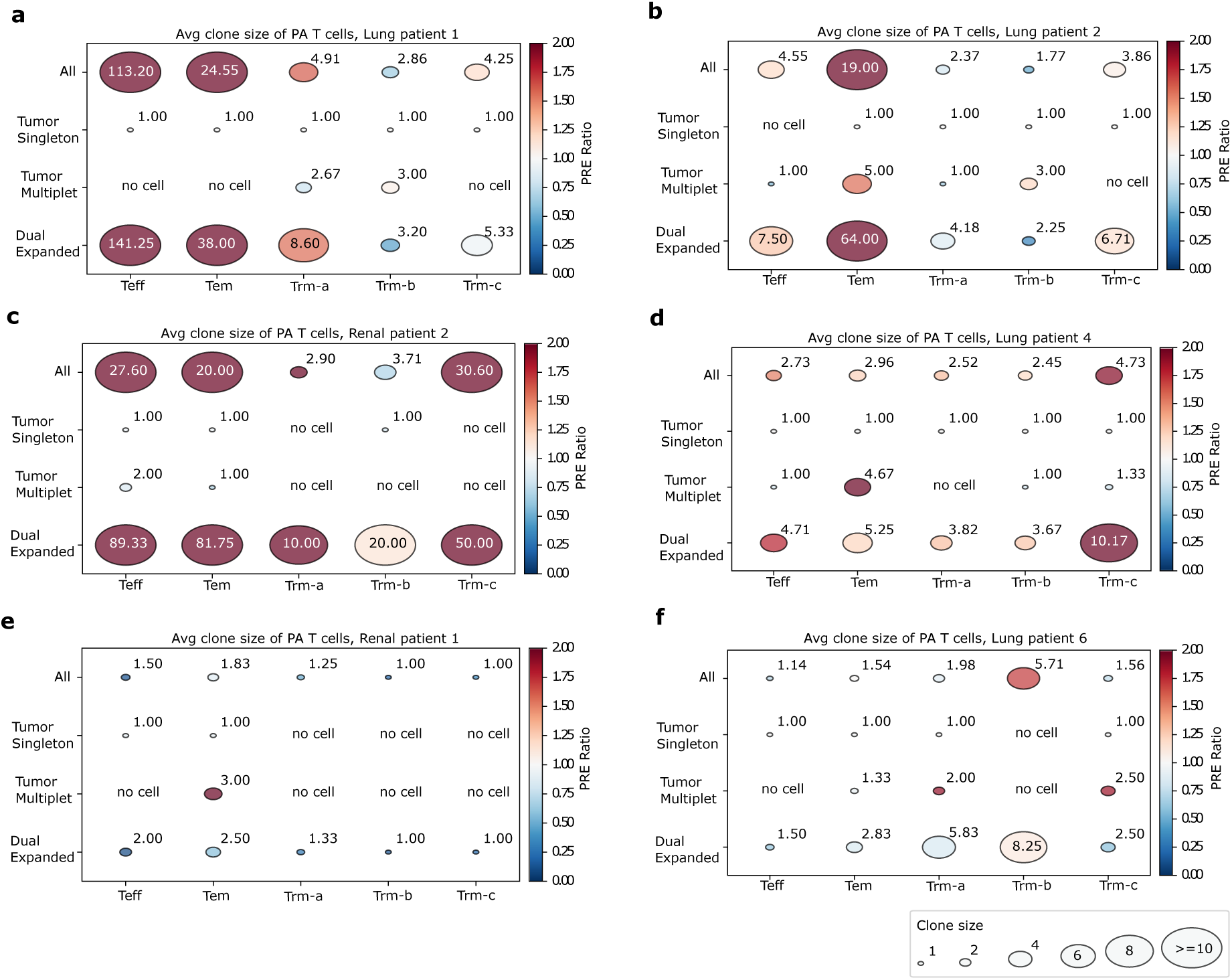
Clonal expansion patterns of Phenotype-Associated T cells across patients. **a-f.** Clonal expansion patterns across four site patterns (All, Tumor Singleton, Tumor Multiplet, and Dual Expanded) and five cell subtypes (Teff, Tem, Trm-a, Trm-b, and Trm-c) in six cancer patients (Lung 1, 2, 4, 6; Renal 1, 2). Color intensity indicates the Phenotype-Relative Expansion (PRE) ratio which means the mean clone size of PA T cells divided by that of NA T cells. Red indicates larger PRE ratio. Circle size represents mean clone size of PA T cells. Significant differences between clone sizes of PA and NA T cells were assessed using a one-sided Mann-Whitney U test, with p-values corrected for multiple testing via the Benjamini–Hochberg method. Only significant p-values are shown. “No cell” indicates absence of specific cell populations in that compartment. The plot highlights the heterogeneity of the clonal expansion patterns of PA T cells across patients.

**Extended Data Fig. 8:**
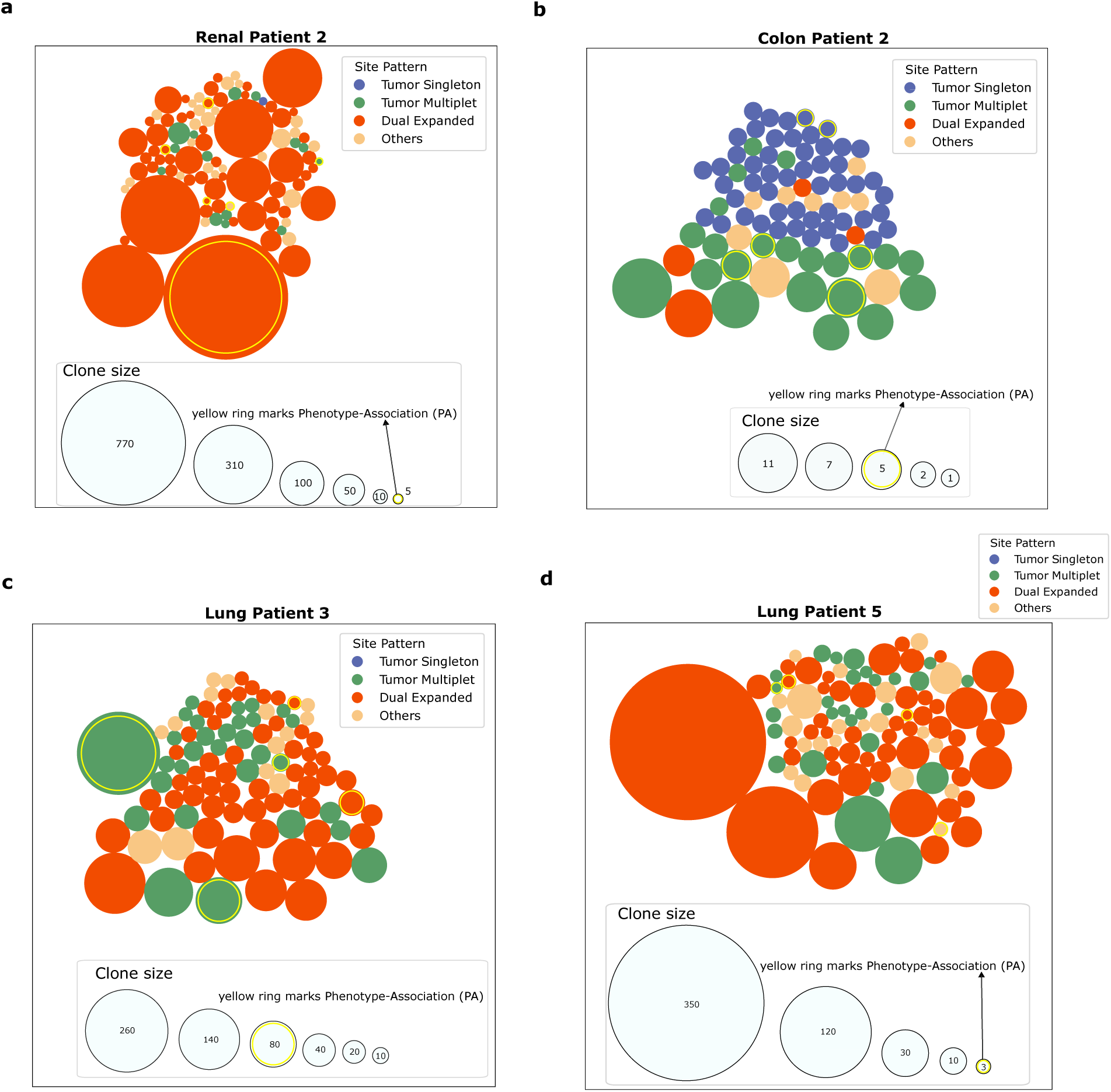
Distribution of clonally expanded T cells in tumor and normal adjacent tissues. **a-d.** Bubble plots illustrating the top 100 clonally expanded T cell clones in four patients: Lung patient 4, Lung patient 3, Endometrium patient 1, and Endometrium patient 3. Two site patterns are highlighted: Tumor Multiplet and Dual Expanded. Circle size represents clone size, while color denotes cell subtype. Phenotype-Associated (PA) T cells are marked by yellow rings. For Tumor Multiplet T cells, the Normal Adjacent Tissue (NAT) region is grayed out, indicating absence of these clones in that area. Dual Expanded T cells are present in both tumor and NAT regions.

**Extended Data Fig. 9:**
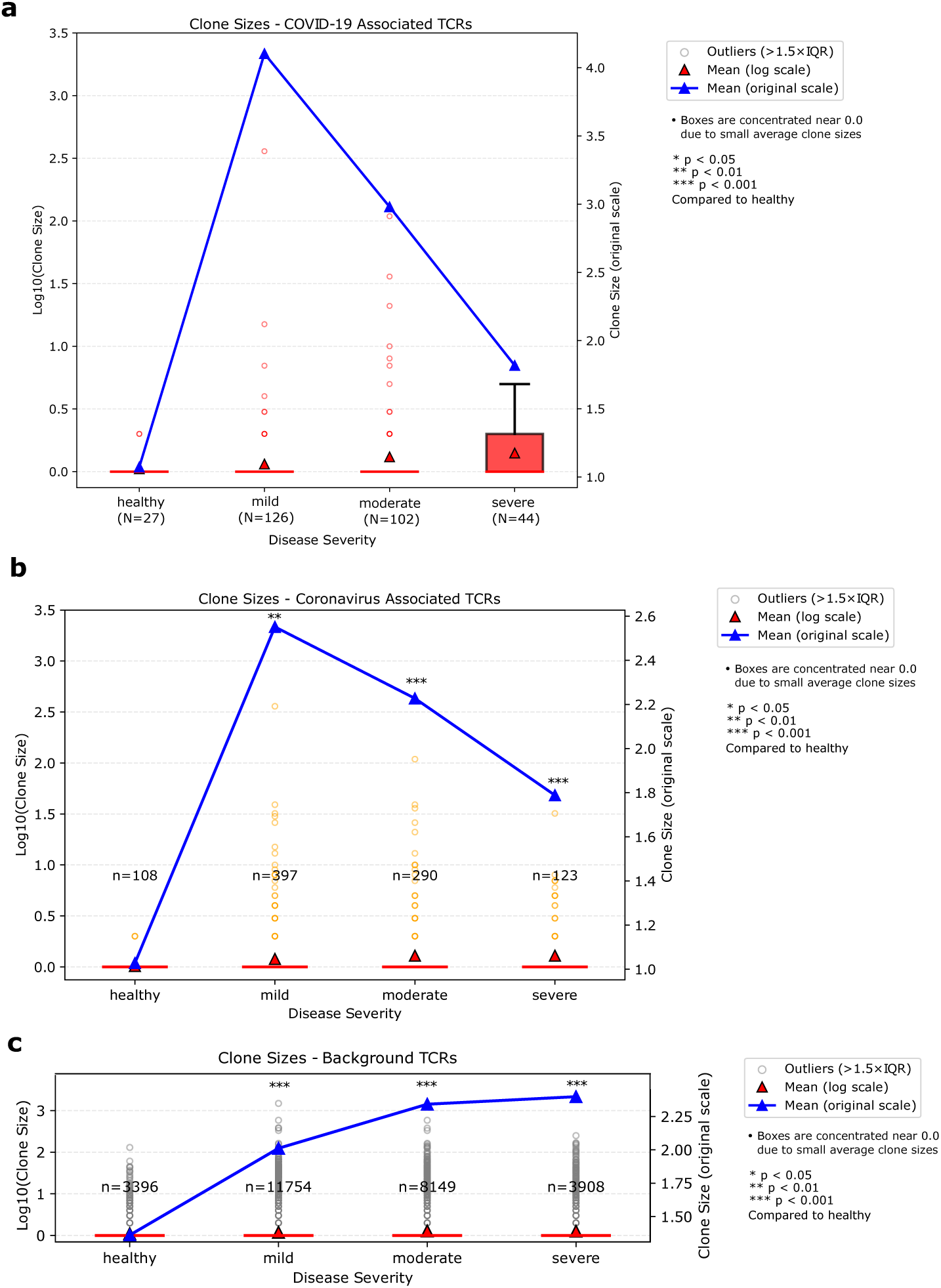
Clonal expansion of COVID-19-Associated, coronavirusassociated, and background T cells by severity group. **a-c.** T cell clonal expansion analysis across COVID-19 disease severity (healthy, mild, moderate, and severe) for three T cell categories: **a.** COVID-19-specific, **b.** broad coronavirus-associated, and **c.** non-COVID-19-associated T cells. T cells were labeled based on EpitopeGen-predicted epitopes followed by database querying to determine phenotype associations. Box plots display clone size distribution (left y-axis, log10 scale), with outliers (colored dots) representing significantly expanded clones. Mean clone sizes are indicated by red triangles (log scale, left y-axis) and blue triangles (original scale, right y-axis). Statistically significant differences compared to the healthy group were determined by one-sided Mann-Whitney U tests with p-values corrected using the Benjamini-Hochberg method.

## References

1. Chaplin, D. D. Overview of the immune response. J Allergy Clin Immunol 125, S3–23 (2010).

2. Andersen, M. H., Schrama, D., Thor Straten, P. & Becker, J. C. Cytotoxic T cells. J Invest Dermatol 126, 32–41 (2006).

3. Tonegawa, S. Somatic generation of antibody diversity. Nature 302, 575–581 (1983).

4. Parham, P. & Ohta, T. Population biology of antigen presentation by MHC class I molecules. Science 272, 67–74 (1996).

5. Shannon, C. E. A mathematical theory of communication. The Bell System Technical Journal 27, 379–423 (1948).

6. Simpson, E. H. Measurement of diversity. Nature 163, 688–688 (1949).

7. Rényi, A. On measures of entropy and information. In Proceedings of the Fourth Berkeley Symposium on Mathematical Statistics and Probability, Volume 1: Contributions to the Theory of Statistics, 547–561 (University of California Press, Berkeley, California, 1961).

8. Greiff, V. et al. A bioinformatic framework for immune repertoire diversity profiling enables detection of immunological status. Genome Med 7, 49 (2015).

9. Janarthanam, R. et al. Bulk t-cell receptor sequencing confirms clonality in pediatric eosinophilic esophagitis and identifies a food-specific repertoire. Allergy 78, 2487–2496 (2023).

10. Vujović, M., Marcatili, P., Chain, B., Kaplinsky, J. & Andresen, T. L. Signatures of T cell immunity revealed using sequence similarity with TCRDivER algorithm. Communications Biology 6, 357 (2023).

11. Huang, H., Wang, C., Rubelt, F., Scriba, T. J. & Davis, M. M. Analyzing the mycobacterium tuberculosis immune response by t-cell receptor clustering with GLIPH2 and genome-wide antigen screening. Nature Biotechnology 38, 1194–1202 (2020).

12. Mayer-Blackwell, K. et al. Tcr meta-clonotypes for biomarker discovery with *tcrdist3* enabled identification of public, hla-restricted clusters of sars-cov-2 tcrs. eLife 10, e68605 (2021).

13. Sidhom, J.-W., Larman, H. B., Pardoll, D. M. & Baras, A. S. DeepTCR is a deep learning framework for revealing sequence concepts within t-cell repertoires. Nature Communications 12, 1605 (2021).

14. Zhang, H., Zhan, X. & Li, B. GIANA allows computationally-efficient TCR clustering and multi-disease repertoire classification by isometric transformation. Nature Communications 12, 4699 (2021).

15. Lu, T. et al. Deep learning-based prediction of the T cell receptor–antigen binding specificity. Nature Machine Intelligence 3, 864–875 (2021).

16. Gao, Y. et al. Pan-Peptide meta learning for t-cell receptor–antigen binding recognition. Nature Machine Intelligence 5, 236–249 (2023).

17. Chronister, W. D. et al. TCRMatch: Predicting T-Cell receptor specificity based on sequence similarity to previously characterized receptors. Front Immunol 12, 640725 (2021).

18. Weber, A., Born, J. & Rodriguez Martínez, M. TITAN: T-cell receptor specificity prediction with bimodal attention networks. Bioinformatics 37, i237–i244 (2021).

19. Jokinen, E., Huuhtanen, J., Mustjoki, S., Heinonen, M. & Lähdesmäki, H. Predicting recognition between T cell receptors and epitopes with TCRGP. PLoS Comput Biol 17, e1008814 (2021).

20. Montemurro, A. et al. NetTCR-2.0 enables accurate prediction of TCR-peptide binding by using paired TCR*α* and *β* sequence data. Communications Biology 4, 1060 (2021).

21. Zhang, W. et al. A framework for highly multiplexed dextramer mapping and prediction of t cell receptor sequences to antigen specificity. Science Advances 7, eabf5835 (2021).

22. Tong, Y. et al. SETE: Sequence-based ensemble learning approach for TCR epitope binding prediction. Comput Biol Chem 87, 107281 (2020).

23. Springer, I., Besser, H., Tickotsky-Moskovitz, N., Dvorkin, S. & Louzoun, Y. Prediction of specific tcr-peptide binding from large dictionaries of tcr-peptide pairs. Frontiers in Immunology 11 (2020).

24. Zhang, J., Ma, W. & Yao, H. Accurate TCR-pMHC interaction prediction using a BERT-based transfer learning method. Briefings in Bioinformatics 25, bbad436 (2023).

25. Peng, X. et al. Characterizing the interaction conformation between t-cell receptors and epitopes with deep learning. Nature Machine Intelligence 5, 395–407 (2023).

26. Cai, M., Bang, S., Zhang, P. & Lee, H. ATM-TCR: TCR-Epitope binding affinity prediction using a Multi-Head Self-Attention model. Front Immunol 13, 893247 (2022).

27. Moris, P. et al. Current challenges for unseen-epitope TCR interaction prediction and a new perspective derived from image classification. Briefings in Bioinformatics 22, bbaa318 (2020).

28. Radford, A. et al. Language models are unsupervised multitask learners (2019).

29. Lewis, M. et al. BART: Denoising sequence-to-sequence pre-training for natural language generation, translation, and comprehension. In Jurafsky, D., Chai, J., Schluter, N. & Tetreault, J. (eds.) Proceedings of the 58th Annual Meeting of the Association for Computational Linguistics, 7871–7880 (Association for Computational Linguistics, Online, 2020).

30. Raffel, C. et al. Exploring the limits of transfer learning with a unified text-to-text transformer. J. Mach. Learn. Res. 21 (2020).

31. Hudson, D., Fernandes, R. A., Basham, M., Ogg, G. & Koohy, H. Can we predict T cell specificity with digital biology and machine learning? Nature Reviews Immunology 23, 511–521 (2023).

32. Grifoni, A. et al. Targets of T cell responses to SARS-CoV-2 coronavirus in humans with COVID-19 disease and unexposed individuals. Cell 181, 1489–1501.e15 (2020).

33. Masopust, D., Murali-Krishna, K. & Ahmed, R. Quantitating the magnitude of the lymphocytic choriomeningitis virus-specific cd8 t-cell response: It is even bigger than we thought. Journal of Virology 81, 2002–2011 (2007).

34. Moutaftsi, M. et al. A consensus epitope prediction approach identifies the breadth of murine TCD8+-cell responses to vaccinia virus. Nature Biotechnology 24, 817–819 (2006).

35. Addo, M. M. et al. Comprehensive epitope analysis of human immunodeficiency virus type 1 (HIV-1)-specific t-cell responses directed against the entire expressed HIV-1 genome demonstrate broadly directed responses, but no correlation to viral load. J Virol 77, 2081–2092 (2003).

36. Pittet, M. J. et al. High frequencies of naive Melan-A/MART-1-specific CD8(+) T cells in a large proportion of human histocompatibility leukocyte antigen (HLA)-A2 individuals. J Exp Med 190, 705–715 (1999).

37. Rizzuto, G. A. et al. Self-antigen-specific CD8+ T cell precursor frequency determines the quality of the antitumor immune response. J Exp Med 206, 849–866 (2009).

38. Nelson, C. E. et al. Robust iterative stimulation with Self-Antigens overcomes CD8(+) T cell tolerance to self- and tumor antigens. Cell Rep 28, 3092–3104.e5 (2019).

39. Kenison, J. E., Stevens, N. A. & Quintana, F. J. Therapeutic induction of antigenspecific immune tolerance. Nature Reviews Immunology 24, 338–357 (2024).

40. Friot, A. et al. Antigen specific activation of cytotoxic CD8(+) T cells by staphylococcus aureus infected dendritic cells. Front Cell Infect Microbiol 13, 1245299 (2023).

41. Shepherd, F. R. & McLaren, J. E. T cell immunity to bacterial pathogens: Mechanisms of immune control and bacterial evasion. Int J Mol Sci 21 (2020).

42. Mittal, J., Ponce, M. G., Gendlina, I. & Nosanchuk, J. D. Histoplasma capsulatum: Mechanisms for pathogenesis. Curr Top Microbiol Immunol 422, 157–191 (2019).

43. Stuckey Peter V. & Santiago-Tirado Felipe H. Fungal mechanisms of intracellular survival: what can we learn from bacterial pathogens? Infection and Immunity 91, e00434–22 (2023).

44. Walker, D. M. et al. Mechanisms of cellular invasion by intracellular parasites. Cell Mol Life Sci 71, 1245–1263 (2013).

45. Morrison, D. A. Evolution of the apicomplexa: where are we now? Trends in Parasitology 25, 375–382 (2009).

46. Stuart, K. et al. Kinetoplastids: related protozoan pathogens, different diseases. The Journal of Clinical Investigation 118, 1301–1310 (2008).

47. Cavicchioli, R., Curmi, P. M., Saunders, N. & Thomas, T. Pathogenic archaea: do they exist? BioEssays 25, 1119–1128 (2003).

48. Gill, E. E. & Brinkman, F. S. L. The proportional lack of archaeal pathogens: Do viruses/phages hold the key? BioEssays 33, 248–254 (2011).

49. UniProt Consortium. UniProt: the universal protein knowledgebase in 2021. Nucleic Acids Res 49, D480–D489 (2021).

50. McCammon, J. A., Gelin, B. R. & Karplus, M. Dynamics of folded proteins. Nature 267, 585–590 (1977).

51. Yost, K. E. et al. Clonal replacement of tumor-specific T cells following PD-1 blockade. Nature Medicine 25, 1251–1259 (2019).

52. Wu, T. D. et al. Peripheral T cell expansion predicts tumour infiltration and clinical response. Nature 579, 274–278 (2020).

53. Devlin, J., Chang, M.-W., Lee, K. & Toutanova, K. Bert: Pre-training of deep bidirectional transformers for language understanding. In North American Chapter of the Association for Computational Linguistics (2019).

54. Shugay, M. et al. VDJdb: a curated database of t-cell receptor sequences with known antigen specificity. Nucleic Acids Res 46, D419–D427 (2018).

55. Vita, R. et al. The immune epitope database (iedb): 2018 update. Nucleic Acids Research 47 (2019).

56. Zhang, W. et al. PIRD: Pan immune repertoire database. Bioinformatics 36, 897–903 (2019).

57. Tickotsky, N., Sagiv, T., Prilusky, J., Shifrut, E. & Friedman, N. McPAS-TCR: a manually curated catalogue of pathology-associated T cell receptor sequences. Bioinformatics 33, 2924–2929 (2017).

58. Chen, S.-Y., Yue, T., Lei, Q. & Guo, A.-Y. TCRdb: a comprehensive database for t-cell receptor sequences with powerful search function. Nucleic Acids Res 49, D468–D474 (2021).

59. Jurtz, V. et al. NetMHCpan-4.0: Improved Peptide-MHC class I interaction predictions integrating eluted ligand and peptide binding affinity data. J Immunol 199, 3360–3368 (2017).

60. O’Donnell, T. J., Rubinsteyn, A. & Laserson, U. Mhcflurry 2.0: Improved pan-allele prediction of mhc class i-presented peptides by incorporating antigen processing. Cell Systems 11, 42–48.e7 (2020).

61. Huang, X. et al. The SysteMHC Atlas v2.0, an updated resource for mass spectrometrybased immunopeptidomics. Nucleic Acids Research 52, D1062–D1071 (2023).

62. Neefjes, J., Jongsma, M. L. M., Paul, P. & Bakke, O. Towards a systems understanding of MHC class I and MHC class II antigen presentation. Nature Reviews Immunology 11, 823–836 (2011).

63. Fan, A., Lewis, M. & Dauphin, Y. Hierarchical neural story generation. In Gurevych, I. & Miyao, Y. (eds.) Proceedings of the 56th Annual Meeting of the Association for Computational Linguistics *(Volume* 1*: Long Papers)*, 889–898 (Association for Computational Linguistics, Melbourne, Australia, 2018).

64. Holtzman, A., Buys, J., Du, L., Forbes, M. & Choi, Y. The curious case of neural text degeneration. In ICLR (OpenReview.net, 2020).

65. Glanville, J. et al. Identifying specificity groups in the T cell receptor repertoire. Nature 547, 94–98 (2017).

66. Nolan, S. et al. A large-scale database of t-cell receptor beta (TCRβ) sequences and binding associations from natural and synthetic exposure to SARS-CoV-2 (2020).

67. Tareen, A. & Kinney, J. B. Logomaker: beautiful sequence logos in python. Bioinformatics 36, 2272–2274 (2020).

68. 10x Genomics. A new way of exploring immunity: Linking highly multiplexed antigen recognition to immune repertoire and phenotype. Application Note, 10x Genomics (2022).

69. Nijkamp, E., Ruffolo, J. A., Weinstein, E. N., Naik, N. & Madani, A. ProGen2: Exploring the boundaries of protein language models. Cell Systems 14, 968–978.e3 (2023).

70. Trolle, T. et al. The length distribution of class i–restricted t cell epitopes is determined by both peptide supply and mhc allele–specific binding preference. The Journal of Immunology 196, 1480–1487 (2016).

71. Ferruz, N., Schmidt, S. & Höcker, B. ProtGPT2 is a deep unsupervised language model for protein design. Nature Communications 13, 4348 (2022).

72. Madani, A. et al. Large language models generate functional protein sequences across diverse families. Nature Biotechnology 41, 1099–1106 (2023).

73. Ashburner, M. et al. Gene ontology: tool for the unification of biology. Nature Genetics 25, 25–29 (2000).

74. Wilkins, M. R. et al. Protein identification and analysis tools in the ExPASy server. Methods Mol Biol 112, 531–552 (1999).

75. Camacho, C. et al. BLAST+: architecture and applications. BMC Bioinformatics 10, 421 (2009).

76. Stranges, P. B. & Kuhlman, B. A comparison of successful and failed protein interface designs highlights the challenges of designing buried hydrogen bonds. Protein Sci 22, 74–82 (2012).

77. Leaver-Fay, A. et al. ROSETTA3: an object-oriented software suite for the simulation and design of macromolecules. Methods Enzymol 487, 545–574 (2011).

78. Yin, R. et al. TCRmodel2: high-resolution modeling of T cell receptor recognition using deep learning. Nucleic Acids Res 51, W569–W576 (2023).

79. Jumper, J. et al. Highly accurate protein structure prediction with AlphaFold. Nature 596, 583–589 (2021).

80. Gonzalez-Galarza, F. F., Christmas, S., Middleton, D. & Jones, A. R. Allele frequency net: a database and online repository for immune gene frequencies in worldwide populations. Nucleic Acids Res 39, D913–9 (2010).

81. Alford, R. F. et al. The rosetta All-Atom energy function for macromolecular modeling and design. J Chem Theory Comput 13, 3031–3048 (2017).

82. Dill, K. A. Dominant forces in protein folding. Biochemistry 29, 7133–7155 (1990).

83. Sehnal, D. et al. Mol* Viewer: modern web app for 3D visualization and analysis of large biomolecular structures. Nucleic Acids Research 49, W431–W437 (2021).

84. Pettersen, E. F. et al. UCSF chimera–a visualization system for exploratory research and analysis. J Comput Chem 25, 1605–1612 (2004).

85. Fridman, W. H., Pagès, F., Sautès-Fridman, C. & Galon, J. The immune contexture in human tumours: impact on clinical outcome. Nature Reviews Cancer 12, 298–306 (2012).

86. Zhang, N. & Bevan, M. J. CD8(+) T cells: foot soldiers of the immune system. Immunity 35, 161–168 (2011).

87. Zhang, L. et al. Lineage tracking reveals dynamic relationships of T cells in colorectal cancer. Nature 564, 268–272 (2018).

88. Simoni, Y. et al. Bystander CD8+ T cells are abundant and phenotypically distinct in human tumour infiltrates. Nature 557, 575–579 (2018).

89. Waldman, A. D., Fritz, J. M. & Lenardo, M. J. A guide to cancer immunotherapy: from T cell basic science to clinical practice. Nature Reviews Immunology 20, 651–668 (2020).

90. Esfahani, K. et al. A review of cancer immunotherapy: From the past, to the present, to the future. Current Oncology 27, 87–97 (2020).

91. Su, Y. et al. Multi-omics resolves a sharp disease-state shift between mild and moderate covid-19. Cell 183, 1479–1495.e20 (2020).

92. Rydyznski Moderbacher, C., et al. Antigen-Specific adaptive immunity to SARS-CoV-2 in acute COVID-19 and associations with age and disease severity. Cell 183, 996–1012.e19 (2020).

93. Chen, Z. & John Wherry, E. T cell responses in patients with COVID-19. Nature Reviews Immunology 20, 529–536 (2020).

94. Henikoff, S. & Henikoff, J. G. Amino acid substitution matrices from protein blocks. Proc Natl Acad Sci U S A 89, 10915–10919 (1992).

95. Sekine, T. et al. Robust T cell immunity in convalescent individuals with asymptomatic or mild COVID-19. Cell 183, 158–168.e14 (2020).

96. Stephenson, E. et al. Single-cell multi-omics analysis of the immune response in COVID-19. Nature Medicine 27, 904–916 (2021).

97. Nelde, A. et al. SARS-CoV-2-derived peptides define heterologous and COVID-19-induced T cell recognition. Nature Immunology 22, 74–85 (2021).

98. Mateus, J. et al. Selective and cross-reactive sars-cov-2 t cell epitopes in unexposed humans. Science 370, 89–94 (2020).

99. Moss, P. The T cell immune response against SARS-CoV-2. Nature Immunology 23, 186–193 (2022).

100. Kared, H. et al. Sars-cov-2–specific cd8+ t cell responses in convalescent covid-19 individuals. The Journal of Clinical Investigation 131 (2021).

101. Gangaev, A. et al. Identification and characterization of a SARS-CoV-2 specific CD8+ T cell response with immunodominant features. Nature Communications 12, 2593 (2021).

102. Zheng, H.-Y. et al. Elevated exhaustion levels and reduced functional diversity of T cells in peripheral blood may predict severe progression in COVID-19 patients. Cellular & Molecular Immunology 17, 541–543 (2020).

103. Functional exhaustion of antiviral lymphocytes in COVID-19 patients. Cellular & Molecular Immunology 17, 533–535 (2020).

104. Braun, J. et al. SARS-CoV-2-reactive T cells in healthy donors and patients with COVID-19. Nature 587, 270–274 (2020).

105. Peng, Y. et al. Broad and strong memory CD4+ and CD8+ T cells induced by SARS-CoV-2 in UK convalescent individuals following COVID-19. Nature Immunology 21, 1336–1345 (2020).

106. Early induction of functional SARS-CoV-2-specific T cells associates with rapid viral clearance and mild disease in COVID-19 patients. Cell Reports 34 (2021).

107. Bluestone, J. A., Herold, K. & Eisenbarth, G. Genetics, pathogenesis and clinical interventions in type 1 diabetes. Nature 464, 1293–1300 (2010).

108. Huseby, E. S., Huseby, P. G., Shah, S., Smith, R. & Stadinski, B. D. Pathogenic CD8 T cells in multiple sclerosis and its experimental models. Frontiers in Immunology 3 (2012).

109. Nielsen, M. et al. NetMHCpan, a method for quantitative predictions of peptide binding to any HLA-A and -B locus protein of known sequence. PLoS One 2, e796 (2007).

110. Sennrich, R., Haddow, B. & Birch, A. Neural machine translation of rare words with subword units. In Erk, K. & Smith, N. A. (eds.) Proceedings of the 54th Annual Meeting of the Association for Computational Linguistics *(Volume* 1*: Long Papers)*, 1715–1725 (Association for Computational Linguistics, Berlin, Germany, 2016).

111. Wolf, T. et al. Transformers: State-of-the-art natural language processing. In Liu, Q. & Schlangen, D. (eds.) Proceedings of the 2020 Conference on Empirical Methods in Natural Language Processing: System Demonstrations, 38–45 (Association for Computational Linguistics, Online, 2020).

112. Vaswani, A., et al. Attention is all you need. In Guyon, I. et al. (eds.) Advances in Neural Information Processing Systems, vol. 30 (Curran Associates, Inc., 2017).

113. Paszke, A., et al. PyTorch: an imperative style, high-performance deep learning library (Curran Associates Inc., Red Hook, NY, USA, 2019).

114. Loshchilov, I. & Hutter, F. Decoupled weight decay regularization. In International Conference on Learning Representations (2017).

115. Barker, D. J. et al. The IPD-IMGT/HLA Database. Nucleic Acids Research 51, D1053–D1060 (2022).

116. Sturm, G. et al. Scirpy: a scanpy extension for analyzing single-cell t-cell receptorsequencing data. Bioinformatics 36, 4817–4818 (2020).

117. Wolf, F. A., Angerer, P. & Theis, F. J. SCANPY: large-scale single-cell gene expression data analysis. Genome Biology 19, 15 (2018).

118. Charoentong, P. et al. Pan-cancer immunogenomic analyses reveal genotypeimmunophenotype relationships and predictors of response to checkpoint blockade. Cell Reports 18, 248–262 (2017).

