## Supplementary Information for "Repertoire-level generation of T-cell epitopes with a large-scale generative transformer"

This document is Supplementary Information.

### 1 Supplementary Note 1. Datasets

**Public paired datasets** Four publicly available datasets of VDJdb[1], PIRD[2], IEDB[3], and McPAS-TCR[4] were used to develop the Robust Affinity Predictor (RAP) and evaluate the generated epitopes. The dataset sizes and basic statistics per split after preprocessing are summarized in **Supplementary Table 1**.

VDJdb is a curated database of T-cell receptor (TCR) sequences with known antigen specificities. This resource aggregates data from previously published studies, providing a comprehensive collection of TCR-antigen interactions. The dataset contains rich metadata that offers detailed information about each TCR-antigen pair, including TCR information (gene, cdr3, v.segm, j.segm, v.end, and j.start), antigen information (antigen.epitope, antigen.gene, and antigen.species), MHC context (mhc.a, mhc.b, mhc.class, and complex.id), quality control and others (species, reference.id, vdjdb.score, and label). Among these, vdjdb.score is a confidence score (0-3) assigned to each entry, reflecting the reliability of the association between the TCR and the antigen in that entry.

The Immune Epitope Database (IEDB) is a comprehensive repository of experimentally determined immune epitope data, encompassing information on both B-cell and T-cell epitopes from human and animal model studies. The database covers various immunological contexts, including infections, allergies, autoimmune diseases, and transplantations. We utilized the IEDB database export v3 (`https://www.iedb.org/database_export_v3.php`), which provides detailed information on epitope sequences (trimmed_seq, original_seq), immune receptor characteristics (receptor_group), antigen source details (source_organisms, source_antigens), labels, and epitopes.

The Pan Immune Repertoire Database (PIRD) was developed to provide a structured repository for sequencing data of the T-cell receptor (TCR) and B-cell receptor (BCR). Within PIRD, the T and B cell Antigen database (TBAdb) is a manually curated dataset of TCRs and BCRs with known antigen specificity. The dataset contains the following columns:disease information (ICDname, Disease.name, Category), antigen details (Antigen, Antigen.sequence, HLA), receptor sequences (Locus, CDR3.alpha.aa, CDR3.beta.aa, CDR3.alpha.nt, CDR3.beta.nt), gene usage (Valpha, Jalpha, Vbeta, Dbeta, Jbeta), experimental methods (Seq.platform, Species, Origin, Nucleotide.type, Cell.subtype, Prepare.method, Evaluate.method), study design (Case.num, Control.type, Control.num, Filteration), publication information (Journal, Pubmed.id), data quality (grade), and label. We used the CDR3.beta.aa and Antigen.sequence columns to collect the binding pairs of TCRs and epitopes.

McPAS-TCR is a manually curated database of TCR sequences associated with various pathological conditions (including pathogen infections, cancer, and autoimmunity) and their respective antigens in humans and mice. The dataset contains comprehensive information organized into the following categories: TCR sequence data (CDR3.alpha.aa, CDR3.beta.aa, CDR3.alpha.nt, CDR3.beta.nt, TRAV, TRAJ, TRBV, TRBD, TRBJ, Reconstructed.J.annotation), study characteristics (Species, Category, Pathology, Pathology.Mesh.ID, Additional.study.details, Antigen.identification.method, Single.cell, NGS, PubMed.ID), antigen and epitope information (Antigen.protein, Protein.ID, Epitope.epitope, Epitope.ID, MHC), T cell properties (Tissue, T.Cell.Type, T.cell.characteristics), and additional information (Mouse.strain, Remarks, label).

**Unpaired datasets** Four unpaired datasets (TCRdb[5], NetMHCPan v4.0[6], MHCFlurry v2.0[7], and SysteMHC[8]) were utilized to collect large-scale unpaired CDR3$\beta$ sequences and epitope sequences for semi-supervised learning. A subset of these datasets was also used for Triple Negative Sampling (TNS) to construct diverse negative samples for the training and testing of the Robust Affinity Predictor (RAP). Dataset sizes and basic statistics per split after preprocessing are summarized in **Supplementary Table 2**.

TCRdb, a comprehensive structured collection of TCR sequencing experiments, contains 131 TCR-seq projects, 8,265 TCR-seq samples, and 277,439,349 TCR CDR3 sequences as of August 2024. From the downloadable dataset (`https://guolab.wchscu.cn/TCRdb/#/download`), we obtained 7,331,478 unique CDR3$\beta$ sequences after preprocessing. The dataset was split into TCRNetSet for training the Robust Affinity Predictor and TCRCandidateSet for BINDSEARCH.

NetMHCPan v4.0 is a predictive model for interactions between class I MHC alleles (represented as MHC pseudo sequences) and epitopes. The project's dataset (`https://services.healthtech.dtu.dk/suppl/immunology/NetMHCpan-4.0`) comprises two components: Binding Affinity (BA) and Eluted Ligand (EL) data. BA data, derived from *in vitro* binding assays, provide quantitative IC50 values that measure the binding strength between epitopes and MHC molecules. EL data, obtained through mass spectrometry (MS) experiments, provide information on naturally processed and presented epitopes, capturing aspects of antigen processing and presentation beyond binding affinity.

MHCFlurry v2.0 is another predictive model for epitope-MHC interactions, which divided the task into antigen presentation prediction and binding affinity prediction. The study compiled datasets from affinity measurements and mass spectrometry experiments, establishing several benchmarks as detailed in the Methods section of the original paper[7].

SysteMHC is an archive of MS-based immunopeptidomics data. Regarding the class I MHC interaction, the atlas includes more than 4,680 MS raw files, 154 allele types, 1 million epitopes, and 457,360 HLA-bound epitopes. The per-allele data in SpectraST format were downloaded from `https://systemhc.sjtu.edu.cn/download`.

The epitopes from NetMHCPan v4.0, MHCFlurry v2.0, and SysteMHC were merged and split into EpiNegSet for training the Robust Affinity Predictor and EpiCandidateSet for BINDSEARCH.

**Pseudo-labeled datasets** The sizes of the pseudo-labeled datasets per split were summarized in **Supplementary Table 3**. These datasets were constructed by evaluating the binding affinities between TCRs from TCRCandidateSet ($I = 6,831,478$) and epitopes from EpiCandidateSet ($J = 20,000,000$). For each TCR, $\beta = 10,000$ epitopes were randomly sampled without replacement from EpiCandidateSet and ranked by binding affinity. The top $n_{\text{max\_tcr}} = 32$ epitopes with the highest binding affinities were selected for each TCR, yielding $I \times n_{\text{max\_tcr}} = 218,607,296$ pairs. Pairs containing redundant epitopes occurring more than $n_{\text{max\_epi}} = 100$ times in the dataset were excluded, resulting in the dataset in **Supplementary Table 3**. While the application of the Antigen Category Filter decreased the size of the pseudo-labeled dataset, it was a necessary quality control step for applying EpitopeGen to repertoire-level datasets.

**External test sets** We utilized two external datasets for additional validation, the statistics of which after preprocessing are summarized in **Supplementary Table 4**.

The dataset released by Glanville et al.[9] comprises TCR sequences from antigen-specific T cells isolated from multiple individuals. Their training set, which we used as an external test set, includes 2,068 unique TCRs of known specificity, derived from T cells sorted using eight different epitope-MHC tetramers in 33 donors.

The dataset released by Nolan et al.[10] provides a large-scale collection of T-cell receptor beta (TCR$\beta$) sequences and their binding associations to the SARS-CoV-2 epitopes. We specifically used their Multiplex Identification of Antigen-Specific T-Cell Receptors Assay (MIRA) dataset, which contains over 135,000 high-confidence SARS-CoV-2-specific TCRs. This dataset maps TCRs binding to SARS-CoV-2 virus epitopes from exposed subjects and naïve controls, offering a comprehensive view of TCR-epitope interactions in the context of COVID-19. We constructed a test set by sampling 2,000 pairs of TCR and epitope by processing `peptide-detail-ci.csv` of `ImmuneCODE-MIRA-Release002.1`.

**10x CD8$^+$ datasets** 10x Genomics Inc. published many single-cell sequencing datasets using their platforms. We used a dataset containing CD8$^+$ T cells from a healthy donor[11], obtained using the Single Cell Immune Profiling platform and analyzed using Cell Ranger 3.0.2. This dataset serves as a test set to simulate a real-world application scenario, distinct from those used in EpitopeGen's training. The model is challenged to generate epitopes for CD8$^+$ T cells based on single-cell TCR sequencing data from an individual patient with specific MHC alleles. This approach assesses the model's ability to generate diverse epitopes with a natural antigen distribution.

**More analysis on the pseudo-labeled data** Here, we provide additional examples of TCRs and the binding affinities of the selected epitopes during the BINDSEARCH process. The epitopes in the pseudo-labeled dataset exhibited consistently high binding affinities to their paired TCRs, while the randomly sampled epitopes showed significantly lower affinities (**Supplementary Fig. 1**). This observation confirms the effectiveness of our selection criteria during pseudo-labeling.

BINDSEARCH Algorithm involves two critical hyperparameters: $n_{\text{max\_tcr}}$ and $n_{\text{max\_epi}}$. The parameter $n_{\text{max\_tcr}}$ specifies the number of high-affinity epitopes selected for each TCR. We set $n_{\text{max\_tcr}} = 32$ in our final implementation, balancing two key considerations: (1) TCR cross-reactivity: TCRs naturally recognize multiple epitopes, necessitating the inclusion of multiple epitopes, and (2) Computational feasibility: the size of the intermediate dataset scales linearly with $n_{\text{max\_tcr}}$. Our initial experiments with $n_{\text{max\_tcr}} = 1$ proved ineffective for generating a large-scale dataset and failed to capture TCR cross-reactivity.

The second parameter, $n_{\text{max\_epi}}$, controls epitope redundancy by excluding epitopes that appear more than $n_{\text{max\_epi}}$ (default value: 100) times in the generated dataset. This filtering step is crucial for ensuring the appropriate epitope diversity. Without this filter, we observed unique epitope ratios as low as 0.0064, leading to excessive redundancy, where the same epitopes were paired with multiple TCRs (**Supplementary Table 5**). Such redundancy is particularly problematic for repertoire-level analysis.

Publicly available datasets showed an average TCR-to-epitope ratio of 86.28 (**Supplementary Table 1**). This high ratio reflects the current limitations of experimental methods such as tetramer assays[12], where a limited set of epitopes is tested against numerous TCRs. Setting $n_{\text{max\_epi}} = 100$, we achieved more balanced ratios of 6.76, 3.18, and 2.36 for Train, Validation, and Test sets, respectively (**Supplementary Table 3**). This controlled redundancy contributes to EpitopeGen's ability to generate more diverse epitopes compared to those found in public datasets, as shown in **Fig. 3e** of the manuscript.

#### 2 Supplementary Note 2. More details on Antigen Category Filter

**Justification for antigen target ratios in Antigen Category Filter** The distribution of antigen categories in Antigen Category Filter (ACF) significantly influences EpitopeGen's characteristics. Although the exact ratio of antigens recognized by $CD8^+$ T cells *in vivo* remains undefined, we can estimate this distribution based on the current immunological understanding.

$CD8^+$ T cells respond primarily to endogenous antigens presented via MHC class I molecules, with viruses representing the predominant category due to their intracellular replication cycle. This intracellular lifecycle leads to extensive endogenous epitope generation and presentation, making viruses the primary targets for $CD8^+$ T cell surveillance. Several studies support this viral dominance: Masopust et al.[13] demonstrated that approximately 80% of splenic $CD8^+$ T cells recognized lymphocytic choriomeningitis virus (LCMV)-derived epitopes during peak primary infection. Moutaftsi et al.[14] found that 29.6% of $CD8^+$ T cells produced IFN-$\gamma$ when exposed to vaccinia virus Western Reserve strain (VACV-WR)-infected cells. Furthermore, Addo et al.[15] reported robust responses of 10,640 spot-forming cells per million PBMC in untreated chronically infected individuals.

Bacteria primarily elicit $CD4^+$ T cell and B cell responses due to their predominantly extracellular nature. However, certain bacterial species have evolved intracellular survival strategies. As Shepherd et al.[16] documented, bacteria such as *Listeria monocytogenes* and *Shigella flexneri* directly target the host cell cytosol, while *Mycobacterium tuberculosis* and *Salmonella* persist in vacuolar compartments. Even traditional 'extracellular pathogens such as *Staphylococcus aureus* can invade intracellular spaces[17], although they represent a smaller proportion of targets of $CD8^+$ T cells compared to viruses.

$CD8^+$ T cells that respond to self-antigens and tumor-associated antigens occur at significantly lower frequencies compared to those that target external pathogens, mainly due to thymic selection mechanisms that maintain immune tolerance[18]. Rizzuto et al.[19] demonstrated that self- / tumor antigen-specific T cells exist at frequencies significantly lower than those specific for foreign antigens, attributing this difference to negative selection in the thymus, a crucial process for preventing autoimmunity. Nelson et al.[20] reported that initial self-reactive T cells are exceptionally rare and difficult to detect prior to antigenic boost, contrasting with the robust responses generated against viral antigens. In quantitative terms, Pittet et al.[21] determined that approximately 1 in 2,500 naive $CD8^+$ T cells recognize tumor (melanoma) antigens, with remarkably similar proportions observed in healthy donors, suggesting that this may represent a baseline frequency for self-antigen recognition.

Fungi and protozoa constitute a more limited set of potential antigens, despite some species causing intracellular infections. Mittal et al.[22] described how *Histoplasma capsulatum* can infect human cells, while Stuckey et al.[23] estimated that among the 3–5 million fungal species thought to exist, only a few dozen regularly cause human infections.

Notably, archaea, despite their presence as human commensals, have not demonstrated pathogenic capabilities and generally do not trigger $CD8^+$ T cell responses[24,25].

This evidence supports five key immunological principles that govern ACF design: (1) Viral Dominance, (2) Limited Bacterial Presence, (3) Endogenous Epitope Presence (Self-Antigen and Tumor-Associated Antigen), (4) Rare Fungi and Parasites, and (5) Absence of Pathogenic Archaea.

**Species identification and categorization protocol** The identification and categorization of species of the generated epitopes follows a systematic process using BLAST[26] (Basic Local Alignment Search Tool) and NCBI taxonomy databases[27]. We performed BLASTP searches on epitope sequences using optimized parameters for short peptides:

```
blastp -task "blastp-short"
       -query FASTA_PATH
       -db DB_PATH
       -out OUT_PATH
       -outfmt "6"
       -evalue "20000"
       -max_target_seqs "10"
       -num_threads "8"
```

The BLASTP output was processed to extract essential information, including the epitope sequence, accession numbers, match details (length, gaps, start/end positions), and statistical metrics (E-value, score). For each epitope, we selected the match with the lowest E-value to ensure the highest confidence in species identification. From NCBI accession numbers, we retrieved TaxIDs using the E-utilities of the Entrez system:

```
cat {accession} | epost -db protein |
    esummary -db protein |
    xtract -pattern DocumentSummary -element Caption,TaxId
```

Subsequently, we obtained detailed taxonomy information in XML format:

```
efetch -db taxonomy -id TAX_IDS -format xml
```

The resulting XML files were parsed to extract lineage information, which was categorized using hierarchical taxonomic classification rules:

1. **Viral Classification**: Species are classified as viruses if they:
   - Belong to viral families (ending in 'viridae')
   - Contain specific viral markers (e.g., HIV, SARS, coronavirus)
   - Include retroviruses and bacteriophages, with special handling for exclusion cases
2. **Superkingdom-Based Classification**: Non-viral species are initially categorized by their superkingdom:

- Bacteria: Classified directly as bacterial species
- Archaea: Classified as archaeal species
- Eukaryota: Further classified based on kingdom and specific characteristics

3. **Eukaryotic Sub-classification**:

   - Fungi: Identified by kingdom markers ('fungi' or 'mycota')
   - Parasites: Recognized through specific genera, families, or parasitic phyla
   - Other eukaryotes: Classified as 'others'

The keywords in **Supplementary Table 6** were used for categorization. Species that cannot be definitively categorized by these rules were labeled as 'others'. This comprehensive approach ensured accurate species identification and categorization, crucial to implementing Antigen Category Filter.

#### 3 Supplementary Note 3. More details on molecular dynamics simulation

**Molecular Dynamics (MD) Simulation Protocol** To provide an orthogonal validation of EpitopeGen's predictions, we used MD simulation to assess the structural stability of TCR-pMHC complexes formed with generated epitopes. This evaluation was particularly important, as it is based on fundamental physicochemical principles, offering an independent validation approach distinct from deep learning. The structural analysis pipeline consisted of three main components: structure prediction, relaxation, and interface analysis.

We used TCRmodel2[28], an AlphaFold2-based model[29] specifically optimized for TCR-pMHC structure prediction. The model requires complete sequences of four components: TCR$\alpha$ chain, TCR$\beta$ chain, MHC molecule, and epitope. To ensure consistent structural context in all simulations, we used fixed template sequences for TCR chains (**Supplementary Table 7**). For each simulation, the CDR3$\beta$ region of the template (CASSYSIRGSRGEQF) was replaced with the input CDR3$\beta$ sequence. HLA-A*02:01 was selected as the MHC allele due to its high prevalence in human populations. We constructed a test set comprising:

1. 20 CDR3$\beta$ sequences randomly sampled from the VDJdb test set
2. Their corresponding ground truth epitopes from VDJdb
3. EpitopeGen-generated epitopes for each CDR3$\beta$ sequence
4. 20 randomly selected epitopes from EpiCandidateSet for each CDR3$\beta$ as controls

This design yielded $20 \times (20+1+1) = 440$ unique TCR-pMHC complexes for comparative analysis, comprising 20 control epitopes, one ground truth epitope, and one EpitopeGen-generated epitope for each of the 20 CDR3$\beta$ sequences. This enables evaluating whether EpitopeGen-generated epitopes demonstrate superior binding characteristics compared to random epitopes and whether they compare with experimentally validated epitopes.

The computational resources to perform the analysis were as follows:

1. Structure Prediction:
   - Tool: TCRmodel2 (AlphaFold2-based)
   - Hardware: NVIDIA V100 GPU
   - Runtime: ≈30 minutes per complex
2. Structure Refinement:
   - Tool: Rosetta relax application
   - Hardware: 64GB RAM, 8 Intel Xeon Gold 6130
   - Runtime: ≈20 minutes per complex
   - Parameters:

```
-relax:default repeats 5 -nstruct 1
```

The structural stability and binding characteristics of each complex were evaluated using `InterfaceAnalyzer`[30] of `Rosetta`. We focused on key properties directly related to binding affinity and interface characteristics. The binding free energy of separated chains (dG_separated) provides a fundamental measure of complex stability. To normalize this measurement across different interfaces, we calculated the binding energy per unit interface area (dG_separated / dSASAx100). Changes in the Solvent Accessible Surface Area (SASA) were examined through two metrics: the change in hydrophobic SASA (dSASA_hphobic) and the total change in interface SASA (dSASA_int). The quality of hydrogen bonding networks was assessed through the number of unsatisfied hydrogen bonds (delta_unsatHbonds). Interface energetics were evaluated separately for both the T-cell receptor (TCR) and peptide-MHC (pMHC) sides of the interface. For each side, we calculated both the raw interface energy score (side1_score for TCR, side2_score for pMHC) and its normalized version (side1_normalized for TCR, side2_normalized for pMHC). These normalized scores account for differences in interface size and composition, enabling more meaningful comparisons between different complexes. More details on these metrics can be found at `https://docs.rosettacommons.org/docs/latest/application_documentation/analysis/interface-analyzer`.

#### 4 Supplementary Note 4. Sensitivity Analysis

**Caveat of interpreting the downstream analysis result** The interpretation of the downstream analysis using EpitopeGen can be influenced by key parameters, most notably the number of generated epitopes ($K$) per TCR. Although higher $K$ values yield increased Phenotype-Associated (PA) ratios, these results require interpretation within their appropriate biological context.

For our analysis of the cancer dataset, we generated the most likely epitope per TCR. However, EpitopeGen's architecture supports the generation of multiple potential epitopes per TCR—a capability that reflects the biological reality of T cell cross-reactivity. In analyzing the COVID-19 dataset, we set $K = 8$, recognizing that COVID-19 represents a single pathogen species with relatively few TCRs that respond specifically to COVID-19-related antigens. Thus, $K$ effectively serves as a sensitivity parameter that modulates the recall of our analysis in detecting phenotype-associated cells.

This parametric sensitivity has important implications for the interpretation of EpitopeGen results. EpitopeGen's insights are most robust when applied comparatively within consistent $K$ values, for instance, when comparing PA ratios across cell types, tracking temporal patterns in immune responses, or evaluating inter-patient group differences. Interpreting absolute PA ratios requires careful consideration, as these values are directly shaped by multiple factors: the chosen $K$ value, reference database completeness, and inherent TCR cross-reactivity. Consequently, while EpitopeGen offers valuable insight into T cell specificity patterns, its results are interpreted more meaningfully within a comparative framework than as absolute measures of antigen specificity.

**Agreement between different model checkpoints** EpitopeGen models and their phenotype-associated (PA) and non-associated (NA) classifications inherently contain multiple sources of variability. To ensure robust and reproducible downstream analysis, we developed an ensemble approach using multiple independently trained models for TCR phenotype association prediction. We propose this ensembling strategy as it is non-trivial to take an ensemble in the prediction space where EpitopeGen was inherently trained to generate diverse epitope sequences from the same input TCRs. Our ensemble framework combines PA predictions from three independently trained models with a consensus threshold. With a threshold of 0.5, a TCR is labeled as PA when at least two models agree, while a threshold of 1.0 requires unanimous agreement across all three models. To generate these ensemble models (`ens_model_1`, `ens_model_2`, and `ens_model_3`), we trained each component using different random seeds in the Antigen Category Filter, resulting in distinct yet balanced training sets that maintain the prescribed antigen category ratio ranges.

Clone size analyses of the Wu et al. cancer dataset using ensembled models revealed consistent patterns across all models: PA T cells, specifically effector (Teff) and effector memory (Tem) subtypes within the Dual Expanded group, exhibited significantly larger clone sizes compared to their non-associated (NA) counterparts. This clonal expansion pattern was notably attenuated in tissue-resident memory subtypes (Trm-a, Trm-b, and Trm-c) (**Supplementary Fig. 2, a–c**). Differential gene expression analysis between PA and NA T cells demonstrated reproducible signatures across all models: PA T cells displayed elevated cytotoxic markers (*GZMB*, *PRF1*), downregulated naive/memory markers, reduced exhaustion markers, enhanced terminal differentiation markers, and increased expression of key transcription factors (*EOMES*, *TBX21*) (**Supplementary Fig. 2, d–f**). These molecular signatures strongly suggest that PA T cells respond actively to tumor-associated antigens.

In the Su et al. COVID-19 dataset, differential gene expression analysis using ensembled models consistently revealed distinct molecular signatures of PA T cells compared to NA T cells: enhanced cytotoxic markers, diminished naive/memory markers, elevated *TIGIT* expression with heterogeneous exhaustion marker patterns, increased terminal differentiation markers, upregulated cytokine expression (*IFNG*), and enhanced transcription factor levels (*EOMES*). This molecular profile indicates an activated state of PA T cells, likely resulting from recognition of COVID-19-associated epitopes (**Supplementary Fig. 3**).

**Sensitivity analysis of the epitope generation count parameter ($K$) in downstream analyses** EpitopeGen's application to single-cell TCR sequencing datasets generates potential cognate epitopes for individual TCR sequences, with the capability to predict multiple distinct epitopes per TCR. The parameter $K$, representing the number of generated epitopes per TCR, influences the identification of Phenotype-Associated (PA) T cells, as each predicted epitope is evaluated against a protein database for phenotype association. While our primary analyses employed $K = 1$ for the cancer dataset and $K = 8$ for the COVID-19 dataset, we conducted comprehensive sensitivity analyses across various $K$ values to assess the robustness of our findings.

In analyzing the cancer dataset, we consistently observed elevated PA ratios (proportion of PA cells within each repertoire group) in effector T cells (Teff) and effector memory T cells (Tem) characterized by the Dual Expanded pattern, which exhibited gene signatures previously correlated with immunotherapy response rates[31]. These key findings remained stable across multiple $K$ values (1–6), validating the robustness of our analytical pipeline (**Supplementary Fig. 4**).

For the COVID-19 dataset, differential gene expression analysis between PA and Non-Associated (NA) cells across disease severity groups revealed distinct molecular signatures. PA T cells across all patients, compared to NA T cells, demonstrated upregulation of cytotoxic markers, terminal differentiation markers, cytokines, and transcription factors, accompanied by downregulation of naive/memory markers. Notably, PA T cells in severe cases did not exhibit enhanced cytotoxicity signatures. These molecular patterns remained consistent across varying $K$ values (7–10) (**Supplementary Fig. 5**). While the magnitude of $\text{Log}_2$ Fold Changes varied with different $K$ values, the preservation of the core biological signatures by different conditions demonstrates the robustness of our differential gene expression analysis to $K$.

#### 5 Supplementary Note 5. More details when training EpitopeGen

**Triple Negative Sampling for Robust Training** Training an effective binding affinity predictor requires both positive and negative training examples. While public datasets provide TCR-epitope pairs with confirmed binding (positive pairs), they lack explicit non-binding pairs (negative pairs). Previous works have addressed this limitation through different approaches. pMTnet[32] generated negative pairs by randomly shuffling positive pairs, assuming that arbitrary TCR-epitope pairs would exhibit low binding affinity. TABR-BERT[33] introduced an alternative approach by pairing pMHC sequences from positive samples with TCR sequences from healthy donors.

We proposed Triple Negative Sampling (TNS), a strategy to generate negative pairs from three distinct sources to enhance model robustness. The first source, internal shuffling, randomly permutes TCR-epitope pairs within the positive dataset, comprising 50% of negative samples in each training batch. The second source, external epitope sampling, pairs TCRs in the positive dataset with epitopes from EpiNegSet, constituting 25% of negative samples and introducing diversity in epitope space while maintaining TCRs. The third source, external TCR sampling, pairs epitopes in the positive dataset with TCRs from TCRNegSet, accounting for the remaining 25% of negative samples and expanding TCR diversity while preserving epitopes. The sampling ratios (50:25:25) were empirically determined to balance between maintaining dataset distributions and introducing external diversity, with the negative pair generation process repeating every five epochs during training. This three-pronged approach offers several advantages: it reduces potential biases from any single sampling strategy and maintains realistic TCR and epitope distributions. The effectiveness of Triple Negative Sampling was demonstrated through improved performance metrics compared to single-source negative sampling strategies (**Extended Data Fig. 1**).

**Sequence Tokenization Strategy** The tokenization scheme for biological sequences should capture meaningful amino acid combinations and efficiently encode variable-length sequences. For TCR sequences, particularly the CDR3$\beta$ region, certain amino acid motifs play crucial roles in antigen recognition. These include conserved triplet motifs associated with antigen specificity, common N-terminal sequences such as 'CASS' in CDR3$\beta$, and frequent C-terminal patterns that mark junction regions.

We used a Byte-Pair Encoding (BPE)[34] tokenization scheme that captured these patterns. The vocabulary size was a hyperparameter to be chosen. We evaluated vocabulary sizes ranging from 50 to 12,800 tokens using two key metrics:

1. Reconstruction ratio ($R_r$): percentage of sequences that can be perfectly reconstructed

$$R_r = \frac{\text{Number of perfectly reconstructed sequences}}{\text{Total number of sequences}} \times 100\% \tag{1}$$

2. Vocabulary utilization ($U_v$): frequency distribution of token usage

$$U_v = \frac{\text{Number of tokens used at least once}}{\text{Total vocabulary size}} \times 100\% \tag{2}$$

Our experiments revealed that a vocabulary size of 400 achieved optimal performance (**Supplementary Fig. 6**): $R_r = 99.98\%$ for CDR3$\beta$ sequences, $R_r = 99.97\%$ for epitope sequences, and $U_v = 92.5\%$ across the training corpus, indicating an efficient representation of biological motifs.

The tokenization process was implemented using the following configuration using the `tokenizers` package in huggingface[35]:

```
tokenization_params = {
    'padding': 'max_length',
    'max_length': 12,
    'truncation': True,
    'return_tensors': 'pt'
}
tokenized_sequence = tokenizer(sequence, **tokenization_params)
```

#### 6 Supplementary Note 6. Additional information on Wu et al. dataset

**Site pattern of $CD8^+$ T cells from the Wu et al. dataset** When processing the dataset from Wu et al.[31], we encoded the site pattern of each TCR using a three-character code. The first character represented the clonal expansion pattern in the tumor tissue, the second character represented the expansion pattern in Normal Adjacent Tissue (NAT), and the third character represented the expansion pattern in peripheral blood. If a TCR was clonally expanded (observed more than once), a capital letter was used. If a TCR was observed only once, a lowercase letter was used. If a TCR was not found on the site, the character 'x' was used. For example, 'TNx' indicated that the TCR was clonally expanded in tumor tissue and NAT, but not observed in blood. Based on this rule, we could find the four site patterns: All (all possible codes), Tumor Singleton ('txb', 'txB', 'txx'), Tumor Multiplet ('Txb', 'TxB', 'Txx'), and Dual Expanded ('tnb', 'tnB', 'tNb', 'tNB', 'Tnb', 'TnB', 'TNb', 'TNB', 'tNx', 'tnx', 'Tnx', 'TNx'). The number of each site pattern in the dataset was summarized in **Supplementary Fig. 7**.

**A curated database of tumor-associated epitopes** To label generated epitopes by Phenotype-Associated or Not-Associated, we constructed a database of tumor-associated epitopes based on IEDB[?] and TCIA[36]. From IEDB, we searched entries with the following filters: Epitope Structure: Linear Sequence, Include Positive Assays, No B cell assays, MHC Restriction Type: Class I, Host: Homo sapiens (human), and Disease Data: cancer (ID:DOID:162), anus cancer (ID:DOID:14110, anal cancer), bone cancer (ID:DOID:184), Reference Type: Journal Article, and Disease Data: cancer (ID:DOID:162), anus cancer (ID:DOID:14110, anal cancer), bone cancer (ID:DOID:184), lung cancer (ID:DOID:1324), skin cancer (ID:DOID:4159), hematologic cancer (ID:DOID:2531, blood cancer), brain cancer (ID:DOID:1319), cecum carcinoma (ID:DOID:1519, Cecal cancer), cecum cancer (ID:DOID:1521), colon cancer (ID:DOID:219), liver cancer (ID:DOID:3571), kidney cancer (ID:DOID:263, renal cancer), uveal cancer (ID:DOID:3479), breast cancer (ID:DOID:1612), cervical cancer (ID:DOID:4362, cervix cancer), muscle cancer (ID:DOID:4045), ocular cancer (ID:DOID:2174), rectum cancer (ID:DOID:1993, rectal cancer). The exported data contained 339,908 entries (June 2024). From TCIA (`https://tcia.at/neoantigens`), we exported all neoantigens data without filtering, which contained 1,011,037 rows. The merge of two data contained 1,350,946 entries.

**Differential gene expression analysis by cell subtypes on the Wu et al. dataset** Building upon our primary analysis of the cancer dataset, which examined differential gene expression between Phenotype-Associated (PA) and Non-Associated (NA) T cells across four site patterns (All, Tumor Singleton, Tumor Multiplet, and Dual Expanded), we conducted a detailed investigation of gene expression patterns across distinct T cell subtypes. PA effector T cells (Teff) exhibited markedly elevated expression of multiple functional markers compared to their NA counterparts, including genes encoding cytotoxic effector molecules (*GZMB*, *PRF1*), T cell signaling and activation markers (*LCK*, *TNF*, *IFNG*), transcription factors (*EOMES*, *TBX21*), and stress response elements (*XBP1*) (**Supplementary Fig. 8**). This comprehensive molecular signature indicates a highly activated effector state with robust cytotoxic potential, characteristic of tumor-reactive T cells.

PA effector memory T cells (Tem) displayed a distinct molecular profile, maintaining highly elevated cytotoxic markers but lacking key cytokine signatures (*IFNG*, *TNF*). Instead, these cells exhibited an increased expression of terminal differentiation markers (*CX3CR1*). These subtype-specific molecular patterns align with the established functional characteristics of antigen-recognizing T cell populations, which additionally validates EpitopeGen's ability to identify tumor-reactive T cells. Notably, these molecular signatures were substantially attenuated in tissue-resident memory subtypes (Trm-a, Trm-b, and Trm-c).

#### 7 Supplementary Note 7. Additional information on the Su et al. dataset

**Cluster annotation of the Su et al. dataset** We characterized each cluster based on their gene expression signatures to assign functional annotations. Clusters 1 and 5 exhibited distinctly high expression of naive T cell markers (*TCF7*, *LEF1*, *SELL*, and *CCR7*), supporting their classification as naive T cells. Clusters 3 and 4 showed elevated expression of memory-associated genes, particularly *GZMK*, along with memory markers *AQP3* and *CD69*, leading to their designation as memory T cells. Clusters 0 and 6 displayed strong cytotoxic signatures characterized by high expression of *NKG7*, *CST7*, *PRF1*, *GZMA*, and *GZMB*, and were thus classified as effector T cells. Cluster 2 exhibited a complex phenotype: while showing elevated proliferation markers (*MKI67*, *TYMS*), it also demonstrated high cytotoxic gene expression, substantial exhaustion markers (*PDCD1*, *TIGIT*, *LAG3*), and intermediate memory signatures. Due to its distinctive proliferative signature, we designated it as proliferative, though this classification represents its most distinguishing features rather than its sole characteristic (**Supplementary Fig. 9**).

**A curated database of COVID-19-associated epitopes** To compile a comprehensive dataset of SARS-CoV-2 epitopes, we integrated data from two major sources: the Immune Epitope Database (IEDB) and MIRA. From IEDB, we retrieved entries using stringent filtering criteria: Linear Sequence epitopes from SARS-CoV-2 (ID:2697049), restricted to positive assays involving MHC Class I presentation, human hosts, and infectious disease contexts, excluding B cell assays and limiting to peer-reviewed journal articles. This initial query yielded 1,780 unique entries. Additionally, we extracted Class I epitopes from MIRA and systematically annotated their protein origins by querying all 546 epitopes against NCBI's non-redundant (nr) protein database using BLASTP. After merging these datasets and removing redundant entries, we obtained a final dataset of 1,587 unique epitopes. These epitopes were subsequently arranged in increasing order of antigen frequency, prioritizing structural proteins that are in smaller quantities compared to non-structural proteins.

For broader coronavirus epitopes, from IEDB, we searched entries with the following filter: Epitope Structure: Linear Sequence, Include Positive Assays, No B cell assays, Host: Homo sapiens (human), Disease Data: Infectious Disease, Reference Type: Journal Article, and Organism: SARS-CoV2 (ID:2697049, COVID-19 virus), Betacoronavirus (ID:694002, Coronavirus), Coronavirus (ID:11118), Alphacoronavirus (ID:693996, Coronavirus), Gammacoronavirus (ID:694013, Coronavirus), Bat coronavirus (ID:1508220), Yak coronavirus (ID:2501420), Human coronavirus 229E (Coronavirus 229E) (ID:11137, Coronavirus 229E), Human coronavirus NL63 (Coronavirus NL63) (ID:277944, Coronavirus NL63), Middle East respiratory syndrome-related coronavirus (MERS coronavirus) (ID:1335626, MERS coronavirus), Severe acute respiratory syndrome coronavirus (SARS coronavirus) (ID:2901879, SARS coronavirus), Avian coronavirus (ID:694014), Betacoronavirus 1 (ID:694003), Coronavirus HKU15 (ID:1965089), Other Coronavirus (ID:11118-other), Alphacoronavirus 1 (Alphacoronavirus-1) (ID:693997, Alphacoronavirus-1), Murine coronavirus (ID:694005), Rodent coronavirus (ID:2050018). In this dataset, we observed that several entries exceeded the canonical MHC Class I epitope length of 10 amino acids. This presented a methodological challenge for our analysis, which employed Levenshtein distance (threshold = 1.0) to query 9-mer epitopes. To address this, we implemented a sliding window approach: for any epitope exceeding 10 amino acids in length, we systematically generated all possible 10-mer subsequences through a sliding window. This preprocessing step ensured consistent length distributions for subsequent sequence similarity analyses while preserving the biological relevance of the epitope sequences.

#### 8 Supplementary Table

| Public Paired Datasets | | | | | |
|---|---|---|---|---|---|
| | **IEDB** | **VDJdb** | **McPAS-TCR** | **PIRD** | **Total** |
| Unique TCR | 86,519 | 6,131 | 10,081 | 5,174 | 98,460 |
| Unique epitope | 914 | 361 | 313 | 54 | 1,141 |
| Total rows | 91,201 | 6,479 | 12,879 | 5,498 | 116,057 |
| TCR-to-epitope ratio | 94.66 | 16.98 | 32.21 | 95.81 | 86.29 |
| **Training Set** | | | | | |
| Unique TCR | 61,393 | 4,334 | 7,290 | 3,644 | 71,263 |
| Unique epitope | 804 | 340 | 288 | 50 | 1,044 |
| Total rows | 63,840 | 4,535 | 9,015 | 3,848 | 81,238 |
| TCR-to-epitope ratio | 76.36 | 12.75 | 25.31 | 72.88 | 68.26 |
| **Validation Set** | | | | | |
| Unique TCR | 18,082 | 1,280 | 2,299 | 1,081 | 22,106 |
| Unique epitope | 512 | 204 | 185 | 43 | 694 |
| Total rows | 18,331 | 1,302 | 2,588 | 1,105 | 23,326 |
| TCR-to-epitope ratio | 35.32 | 6.27 | 12.43 | 25.14 | 31.85 |
| **Test Set** | | | | | |
| Unique TCR | 8,948 | 638 | 1,134 | 538 | 11,092 |
| Unique epitope | 388 | 146 | 125 | 36 | 518 |
| Total rows | 9,030 | 642 | 1,276 | 545 | 11,493 |
| TCR-to-epitope ratio | 23.06 | 4.37 | 9.07 | 14.94 | 21.41 |

**Supplementary Table** 1: **Summary of publicly available paired TCR and epitope datasets.** Unique TCR and unique epitope denote the number of unique TCRs and epitopes in the dataset. Total rows denote the number of pairs included. TCR-to-epitope ratio is unique TCR divided by unique epitope. The Train:Val:Test was split into the 7:2:1 ratio. The datasets were used to train Robust Affinity Predictor. The train set was merged with the pseudo-labeled dataset to train EpitopeGen. The test set was used to assess the binding affinities and naturalness of the generated epitopes.

| Unpaired Peptide Dataset Sources | | | |
|---|---|---|---|
| **Type** | **NetMHCPan** | **MHCFlurry** | **SysteMHC** |
| Unique peptide | 12,081,588 | 13,195,789 | 449,852 |
| Unique class I peptide | 9,147,682 | 13,178,655 | 442,746 |
| **Merge of NetMHCPan, MHCFlurry, and SysteMHC** | **EpiNegSet (for training Robust Affinity Predictor)** | **EpiCandidateSet (for pseudo-labeling and training EpitopeGen)** | **Total** |
| Train | 1,601,187 | | |
| Val | | | |
| Test | 200,000 | | |
| Total | 1,801,187 | 20,000,000 | 21,801,187 |
| **Unpaired TCR Dataset Sources** | | | |
| **TCRdb** | **TCRNegSet (for training Robust Affinity Predictor)** | **TCRCandidateSet (for pseudo-labeling and training EpitopeGen)** | **Total** |
| Train | 380,000 | | |
| Val | | | |
| Test | 120,000 | | |
| Total | 500,000 | 6,831,478 | 7,331,478 |

**Supplementary Table** 2: **Summary of the unpaired datasets of TCR and peptide sequences.** The peptide datasets are from three sources: NetMHCPan v4.0, MHCFlurry v2.0, and SysteMHC, which were preprocessed, merged, and split into two: EpiNegSet and EpiCandidateSet. The unpaired TCRs were obtained from the TCRdb dataset and were split into two: TCRNegSet and TCRCandidateSet.

| $(n_{\text{max\_tcr}}, n_{\text{max\_epi}}) = (32, 100)$, **after ACF** | **Train** | **Val** | **Test** |
|---|---|---|---|
| Unique TCR | 480,423 | 120,563 | 59,815 |
| Unique epitope | 71,028 | 37,876 | 25,332 |
| Total rows | 527,828 | 123,602 | 60,556 |
| TCR-to-epitope ratio | 6.76 | 3.18 | 2.36 |

| $(n_{\text{max\_tcr}}, n_{\text{max\_epi}}) = (32, 100)$, **before ACF** | **Train** | **Val** | **Test** |
|---|---|---|---|
| Unique TCR | 3,928,055 | 2,044,980 | 1,251,077 |
| Unique epitope | 1,037,034 | 710,506 | 546,014 |
| Total rows | 11,836,453 | 3,398,753 | 1,674,013 |
| TCR-to-epitope ratio | 3.79 | 2.88 | 2.29 |

**Supplementary Table** 3: **Summary of the pseudo-labeled datasets per split.** Here, $n_{\text{max\_tcr}}$ denotes the maximum number of selected epitopes for a TCR and $n_{\text{max\_epi}}$ denotes the maximum occurrences of an epitope. The TCR-to-epitope ratio is the number of unique TCRs divided by the number of unique epitopes.

| **Glanville** | **Total (Test)** |
|---|---|
| Unique TCR | 1,966 |
| Unique epitope | 7 |
| Total rows | 4,120 |
| TCR-to-epitope ratio | 280.86 |
| **MIRA** | **Total (Test)** |
| Unique TCR | 136,463 |
| Unique epitope | 419 |
| Total rows | 154,320 |
| TCR-to-epitope ratio | 325.69 |

**Supplementary Table** 4: **Summary of external datasets.** The external datasets are from two sources: the Glanville dataset published by Glanville et al.[9] and the MIRA dataset published by Nolan et al.[10] The datasets show a high TCR-to-epitope ratio, indicating relatively low diverse peptides relative to TCRs.

| $n_{\text{max\_epi}}$ | **Inf** | **800** | **400** | **200** | **100** |
|---|---|---|---|---|---|
| Total rows | 224,526,336 | 81,575,641 | 50,091,251 | 29,489,056 | 16,909,219 |
| Unique tcr | 6,762,496 | 6,757,435 | 6,577,021 | 5,787,948 | 4,514,443 |
| Unique epitope | 1,437,906 | 1,354,151 | 1,298,733 | 1,226,493 | 1,138,726 |
| TCR-to-epitope ratio | 4.7 | 4.99 | 5.06 | 4.72 | 3.96 |
| Unique epitope ratio | 0.00640 | 0.0165 | 0.0259 | 0.0415 | 0.0673 |
| Reduction ratio | - | 0.636 | 0.777 | 0.869 | 0.925 |

**Supplementary Table** 5: **Summary of the pseudo-labeled datasets by** $n_{\text{max\_tcr}}$**.** Here, $n_{\text{max\_epi}}$ denotes the maximum occurrences of an epitope, TCR-to-epitope ratio is given by the number of unique TCRs divided by the number of unique epitopes, unique epitope ratio is given by the number of unique epitopes divided by total rows, and reduction ratio is given by (224,526,336 - total rows)/224,526,336. The table highlights that the data before the Redundancy Filter contains a large number of redundant epitopes.

| Category | Classification Criteria |
|---|---|
| Virus | **Families:** Viridae suffixes<br>**Special markers:** HIV, immunodeficiency, SARS, coronavirus<br>**Retroviral markers:** Retroviridae, retrovirus<br>**Phage families:** Caudovirales, Myoviridae, Siphoviridae, Podoviridae<br>**Exclusions:** Bacteriophage, provirus |
| Parasite | **Genera:** *Plasmodium*, *Trypanosoma*, *Leishmania*, *Toxoplasma*, *Cryptosporidium*, *Giardia*, *Entamoeba*, *Trichomonas*, *Babesia*, *Theileria*, *Schistosoma*, *Fasciola*, *Taenia*, *Echinococcus*, and others<br>**Families:** Plasmodiidae, Trypanosomatidae, Toxoplasmatidae, Cryptosporidiidae, Giardiidae, Entamoebidae, Trichomonadidae, and others<br>**Phyla:** Apicomplexa, Platyhelminthes, Nematoda, Arthropoda<br>**Protozoan phyla:** Apicomplexa, Ciliophora, Euglenozoa, Amoebozoa, Microsporidia |
| Fungi | **Kingdom markers:** fungi, mycota |
| Bacteria | **Superkingdom:** Bacteria |
| Archaea | **Superkingdom:** Archaea |

**Supplementary Table** 6: **Taxonomic classification criteria for antigen categories.** The classification criteria for each category used to categorize the species according to lineage information. Bacteriophages were excluded because they infect bacteria and proviruses were excluded as they behave like self-antigens.

| Protein | Amino Acid Sequence |
|---|---|
| TCR Alpha Chain Template | AQEVTQIPAALSVPEGENLVLNCSFTDSAIYNLQWFRQDPGK<br>GLTSLLLIQSSQREQTSGRLNASLDKSSGRSTLYIAASQPGD<br>SATYLCAVTNQAGTALIFGKGTTLSVSS |
| TCR Beta Chain Template | NAGVTQTPKFQVLKTGQSMTLQCSQDMNHEYMSWYRQDPGM<br>GLRLIHYSVGAGITDQGEVPNGYNVSRSTTEDFPLRLLSAA<br>PSQTSVYFCASSYSIRGSRGEQFFGPGTRLTVL |
| HLA-A*02:01 Sequence | MAVMAPRTLVLLLSGALALTQTWAGSHSMRYFFTSVSRPGRGEPRFIAV<br>GYVDDTQFVRFDSDAASQRMEPRAPWIEQEGPEYWDGETRKVKAHSQTH<br>RVDLGTLRGYYNQSEAGSHTVQRMYGCDVGSDWRFLRGYHQYAYDGKDY<br>IALKEDLRSWTAADMAAQTTKHKWEAAHVAEQLRAYLEGTCVEWLRRYL<br>ENGKETLQRTDAPKTHMTHHAVSDHEATLRCWALSFYPAEITLTWQRDG<br>EDQTQDTELVETRPAGDGTFQKWAAVVVPSGQEQRYTCHVQHEGLPKPL<br>TLRWEPSSQPTIPIVGIIAGLVLFGAVITGAVVAAVMWRRKSSDRKGGS<br>YSQAASSDSAQGSDVSLTACKV |

**Supplementary Table 7**: **Protein sequences used in molecular dynamics (MD) simulations.** The table presents the amino acid sequences of three proteins used in MD simulations and subsequently processed through TCRmodel2 for structure prediction. The TCR alpha chain and the HLA-A02:01 sequences remained constant throughout the simulations. To evaluate structural stability, the CDR3$\beta$ region of the TCR beta chain template was substituted with the generated epitope sequences. HLA-A02:01 was chosen for this analysis because it is a predominant human HLA allele.

### 9 Supplementary Figure

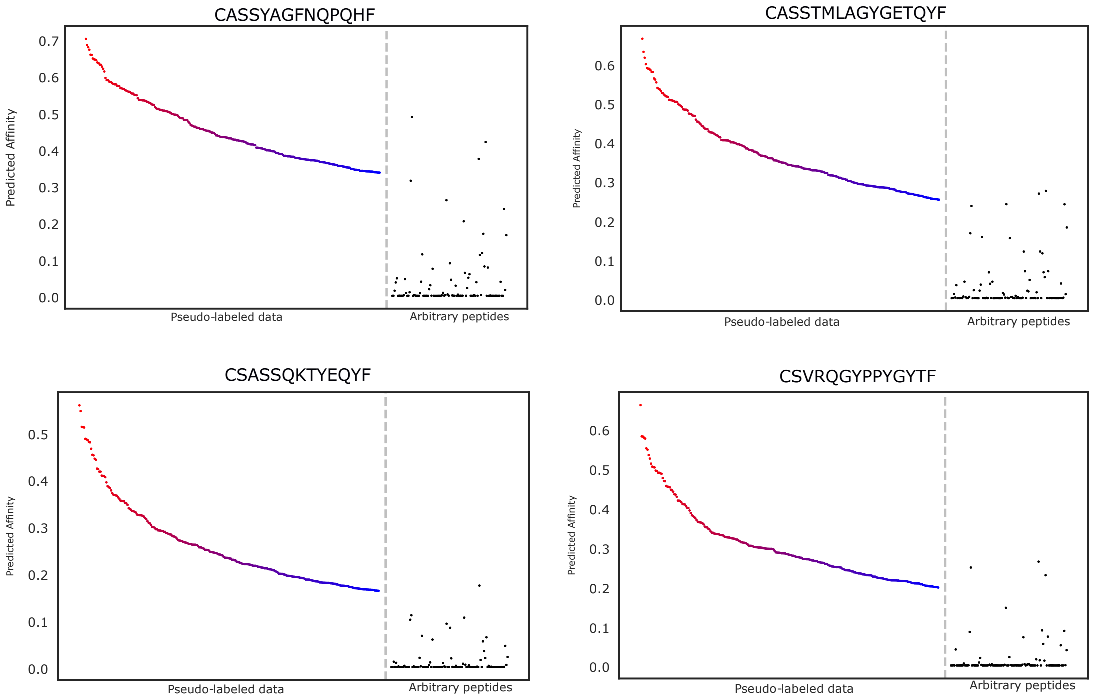

**Supplementary Fig.** 1: **Binding affinity distribution for example TCRs against peptides from pseudo-labeled and random datasets.** The y-axis represents binding affinity, while the x-axis represents peptide sources. In each figure, the left part shows the binding affinity between TCRs and the peptides from pseudo-labeled data, ordered by binding affinity, while the right part shows the binding affinity between TCRs and arbitrary peptides.

**a.**

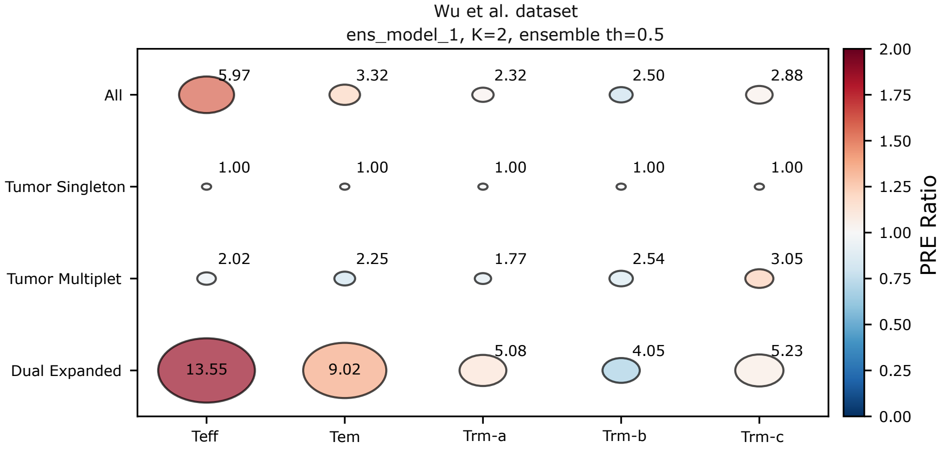

Wu et al. dataset
ens_model_1, K=2, ensemble th=0.5
All
Tumor Singleton
Tumor Multiplet
Dual Expanded
5.97
3.32
2.32
2.50
2.88
1.00
1.00
1.00
1.00
1.00
2.02
2.25
1.77
2.54
3.05
13.55
9.02
5.08
4.05
5.23
Teff
Tem
Trm-a
Trm-b
Trm-c
2.00
1.75
1.50
1.25
1.00
0.75
0.50
0.25
0.00
PRE Ratio

**b.**

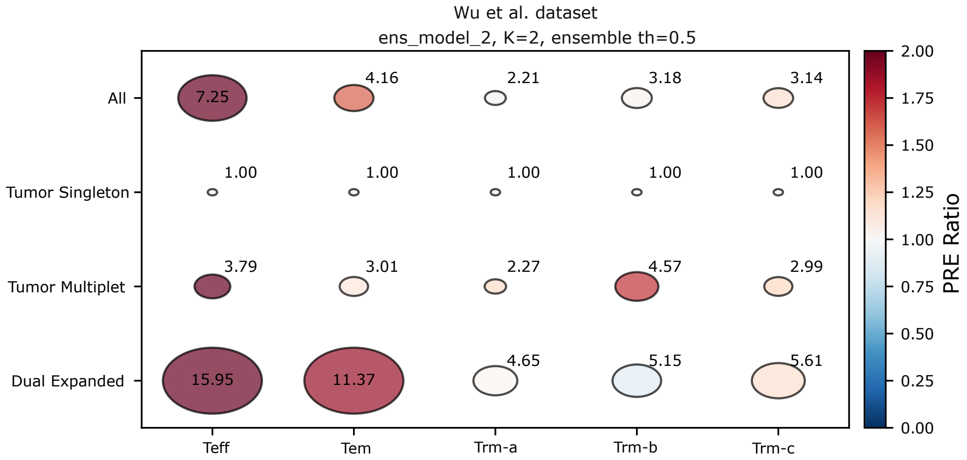

Wu et al. dataset
ens_model_2, K=2, ensemble th=0.5
All
Tumor Singleton
Tumor Multiplet
Dual Expanded
7.25
4.16
2.21
3.18
3.14
1.00
1.00
1.00
1.00
1.00
3.79
3.01
2.27
4.57
2.99
15.95
11.37
4.65
5.15
5.61
Teff
Tem
Trm-a
Trm-b
Trm-c
2.00
1.75
1.50
1.25
1.00
0.75
0.50
0.25
0.00
PRE Ratio

**c.**

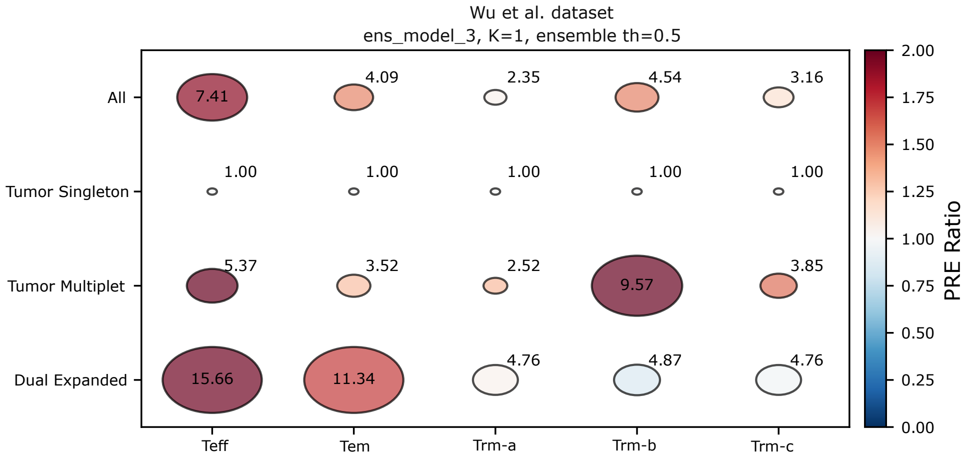

Wu et al. dataset
ens_model_3, K=1, ensemble th=0.5
All
Tumor Singleton
Tumor Multiplet
Dual Expanded
7.41
4.09
2.35
4.54
3.16
1.00
1.00
1.00
1.00
1.00
5.37
3.52
2.52
9.57
3.85
15.66
11.34
4.76
4.87
4.76
Teff
Tem
Trm-a
Trm-b
Trm-c
2.00
1.75
1.50
1.25
1.00
0.75
0.50
0.25
0.00
PRE Ratio

**d.**

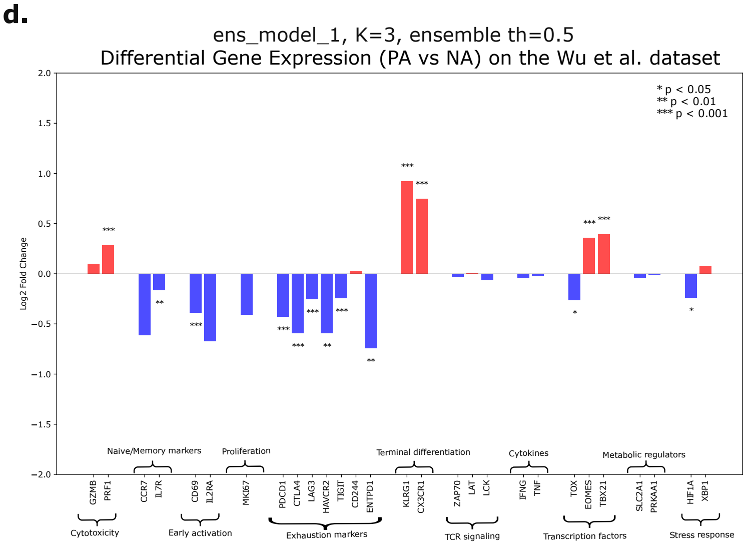
ens_model_1, K=3, ensemble th=0.5
Differential Gene Expression (PA vs NA) on the Wu et al. dataset
* p < 0.05
** p < 0.01
*** p < 0.001
Log2 Fold Change
Naive/Memory markers
Proliferation
Terminal differentiation
Cytokines
Metabolic regulators
GZMB
PRF1
CCR7
IL7R
CD69
IL2RA
MKI67
PDCD1
CTLA4
LAG3
HAVCR2
TIGIT
CD244
ENTPD1
KLRG1
CX3CR1
ZAP70
LAT
LCK
IFNG
TNF
TOX
EOMES
TBX21
SLC2A1
PRKAA1
HIF1A
XBP1
Cytotoxicity
Early activation
Exhaustion markers
TCR signaling
Transcription factors
Stress response

**e.**

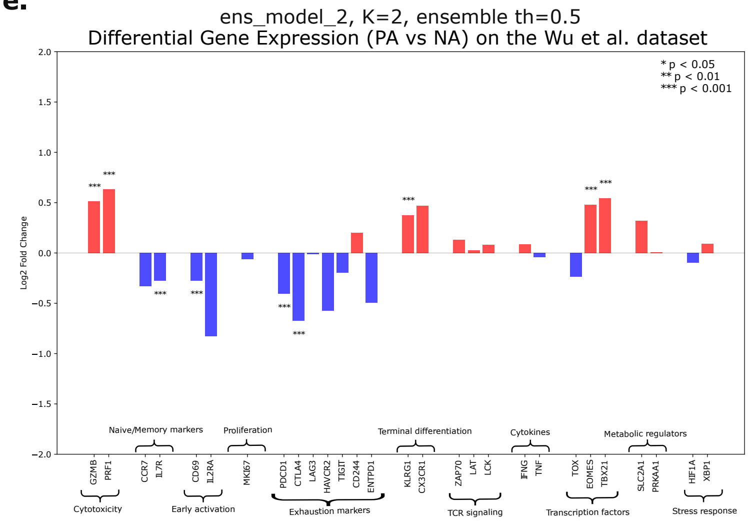
ens_model_2, K=2, ensemble th=0.5
Differential Gene Expression (PA vs NA) on the Wu et al. dataset
* p < 0.05
** p < 0.01
*** p < 0.001
Log2 Fold Change
Naive/Memory markers
Proliferation
Terminal differentiation
Cytokines
Metabolic regulators
GZMB
PRF1
CCR7
IL7R
CD69
IL2RA
MKI67
PDCD1
CTLA4
LAG3
HAVCR2
TIGIT
CD244
ENTPD1
KLRG1
CX3CR1
ZAP70
LAT
LCK
IFNG
TNF
TOX
EOMES
TBX21
SLC2A1
PRKAA1
HIF1A
XBP1
Cytotoxicity
Early activation
Exhaustion markers
TCR signaling
Transcription factors
Stress response

**f.**

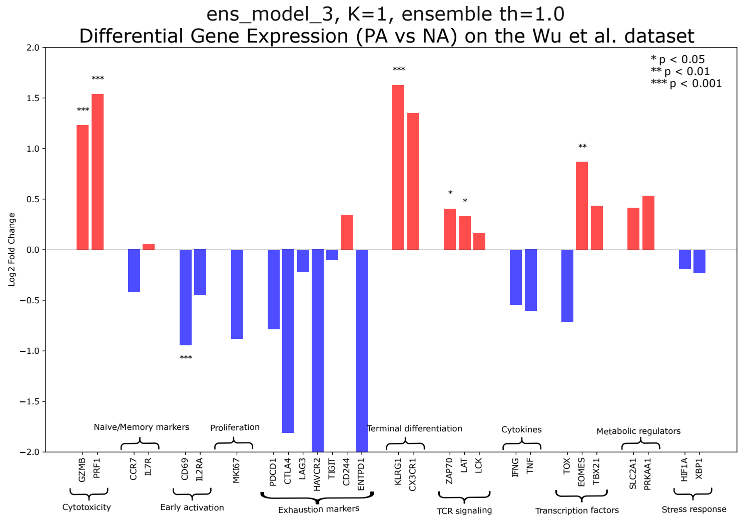
ens_model_3, K=1, ensemble th=1.0
Differential Gene Expression (PA vs NA) on the Wu et al. dataset
* p < 0.05
** p < 0.01
*** p < 0.001
Log2 Fold Change
Naive/Memory markers
Proliferation
Terminal differentiation
Cytokines
Metabolic regulators
GZMB
PRF1
CCR7
IL7R
CD69
IL2RA
MKI67
PDCD1
CTLA4
LAG3
HAVCR2
TIGIT
CD244
ENTPD1
KLRG1
CX3CR1
ZAP70
LAT
LCK
IFNG
TNF
TOX
EOMES
TBX21
SLC2A1
PRKAA1
HIF1A
XBP1
Cytotoxicity
Early activation
Exhaustion markers
TCR signaling
Transcription factors
Stress response

**Supplementary Fig.** 2: **Consistency analysis of three ensemble models on the Wu et al. dataset**. **a–c.** Clonal expansion analysis across four site patterns (All, Tumor Singleton, Tumor Multiplet, and Dual Expanded) and five T cell subtypes (Teff, Tem, Trm-a, Trm-b, and Trm-c) using three ensemble models: (**a**) `ens_model_1` with generation number $K = 2$, (**b**) `ens_model_2` with $K = 2$, and (**c**) `ens_model_3` with $K = 1$. The ensemble threshold was set to 0.5, labeling a TCR as Phenotype-Associated (PA) if at least two models agreed (average PA label $> 0.5$). The color scale represents the Phenotype-Relative Expansion Ratio, defined as the average clone size of PA T cells divided by that of NA T cells. Red indicates higher clonal expansion of PA T cells. Circle size and annotated numbers denote the average clone size of PA T cells. **d–f.** Differential gene expression analysis between PA and NA T cells using three ensemble models: (**d**) `ens_model_1` with $K = 3$, (**e**) `ens_model_2` with $K = 2$, and (**f**) `ens_model_3` with $K = 1$. $\text{Log}_2$ fold changes were calculated using the two-sided Wilcoxon rank-sum test from Scanpy, and p-values were adjusted for multiple testing using the Benjamini-Hochberg method.

**a.**

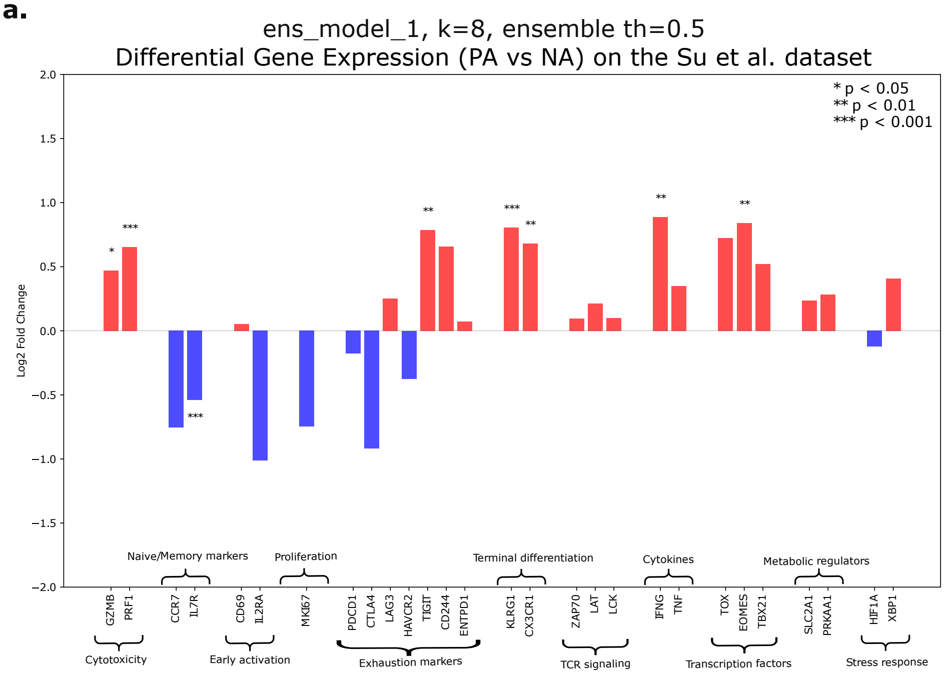
ens_model_1, k=8, ensemble th=0.5
Differential Gene Expression (PA vs NA) on the Su et al. dataset
* p < 0.05
** p < 0.01
*** p < 0.001
Log2 Fold Change
2.0
1.5
1.0
0.5
0.0
−0.5
−1.0
−1.5
−2.0
GZMB
PRF1
CCR7
IL7R
CD69
IL2RA
MKI67
PDCD1
CTLA4
LAG3
HAVCR2
TIGIT
CD244
ENTPD1
KLRG1
CX3CR1
ZAP70
LAT
LCK
IFNG
TNF
TOX
EOMES
TBX21
SLC2A1
PRKAA1
HIF1A
XBP1
Cytotoxicity
Naive/Memory markers
Early activation
Proliferation
Exhaustion markers
Terminal differentiation
TCR signaling
Cytokines
Transcription factors
Metabolic regulators
Stress response

**b.**

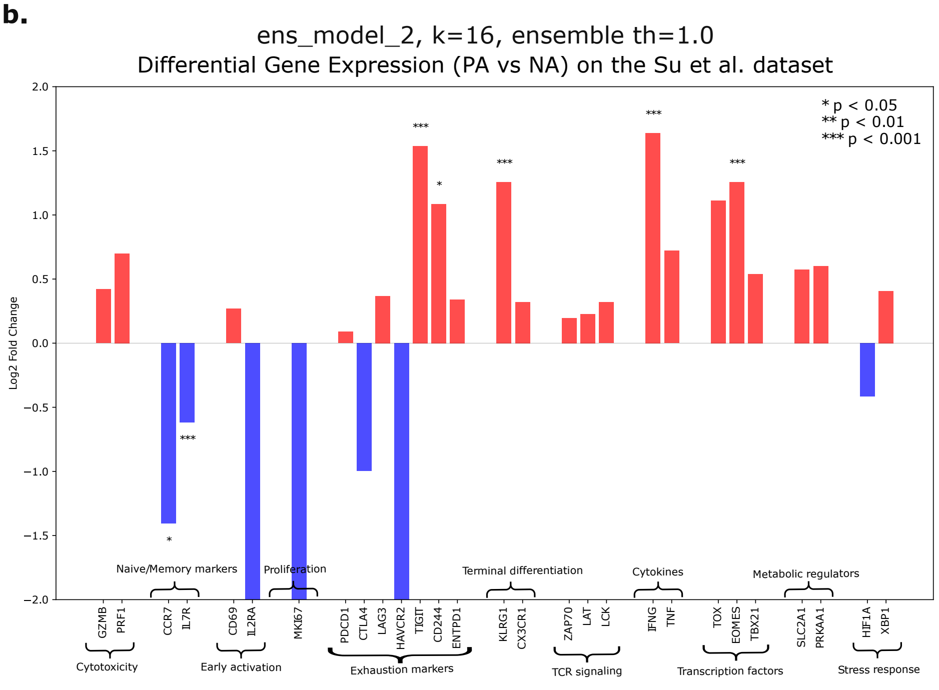
ens_model_2, k=16, ensemble th=1.0
Differential Gene Expression (PA vs NA) on the Su et al. dataset
* p < 0.05
** p < 0.01
*** p < 0.001
Log2 Fold Change
2.0
1.5
1.0
0.5
0.0
−0.5
−1.0
−1.5
−2.0
GZMB
PRF1
CCR7
IL7R
CD69
IL2RA
MKI67
PDCD1
CTLA4
LAG3
HAVCR2
TIGIT
CD244
ENTPD1
KLRG1
CX3CR1
ZAP70
LAT
LCK
IFNG
TNF
TOX
EOMES
TBX21
SLC2A1
PRKAA1
HIF1A
XBP1
Cytotoxicity
Naive/Memory markers
Early activation
Proliferation
Exhaustion markers
Terminal differentiation
TCR signaling
Cytokines
Transcription factors
Metabolic regulators
Stress response

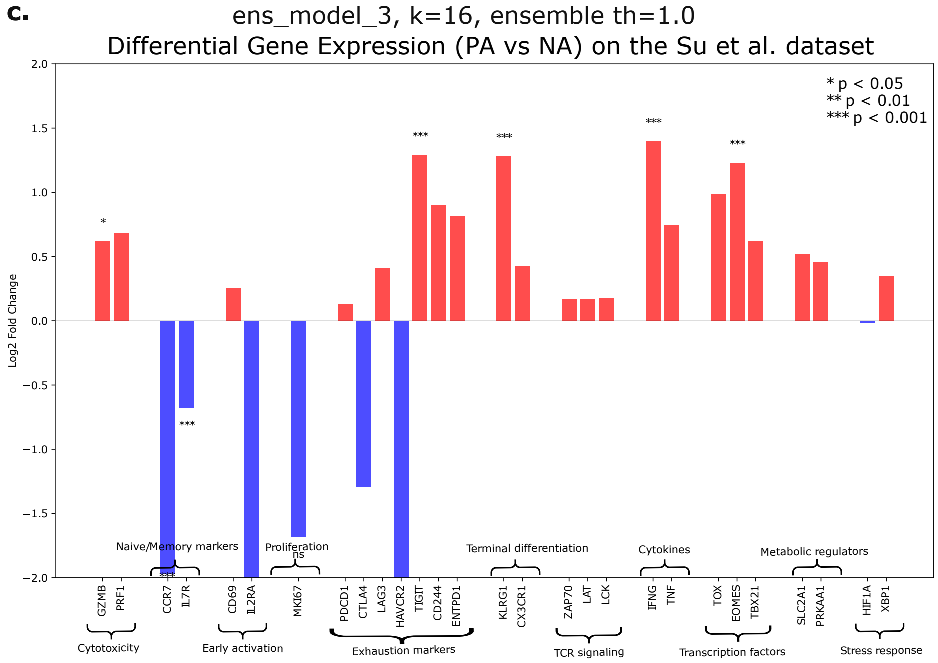

**Supplementary Fig.** 3: **Consistency analysis of three ensemble models on the Su et al. dataset**. **a–c.** Differential gene expression analysis between Phenotype-Associated (PA) and Not-Associated (NA) T cells using three ensemble models: (**a**) `ens_model_1` with generation number $K = 8$, (**b**) `ens_model_2` with $K = 16$, and (**c**) `ens_model_3` with $K = 16$. The ensemble threshold of 0.5 indicates that a TCR is labeled as PA if at least two models agree, while a threshold of 1.0 requires consensus among all three models. $\text{Log}_2$ fold changes were computed using the two-sided Wilcoxon rank-sum test implemented in Scanpy, and p-values were corrected for multiple testing using the Benjamini-Hochberg method.

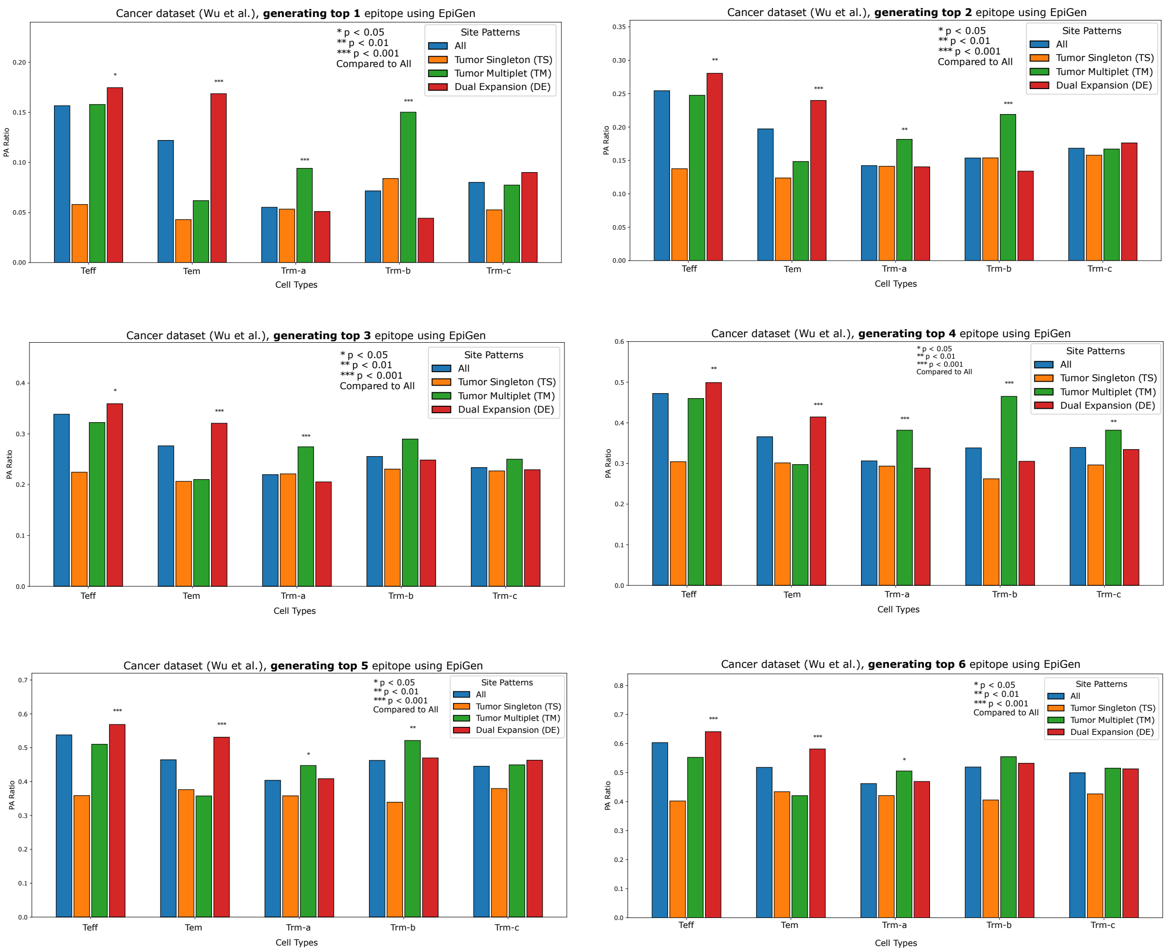

**Supplementary Fig. 4**: **Sensitivity analysis of Phenotype-Association ratios across multiple epitope generation values.** Analysis of Phenotype-Associated (PA) T cell proportions across different cell subtypes and site patterns as a function of epitope generation count ($K$). Six values of $K$ (1, 2, 3, 4, 5, and 6) were considered. The PA ratio, defined as the proportion of PA T cells within each TCR repertoire, was calculated for increasing values of $K$, leveraging EpitopeGen's capability to generate multiple potential epitope sequences per TCR. One-sided Fisher's exact tests were performed to compare PA proportions of each site pattern to the 'All' category within each cell subtype. P-values were adjusted using the Benjamini–Hochberg method (*$p < 0.05$, **$p < 0.01$, ***$p < 0.001$). Sample sizes (n) for site patterns [All, Tumor Singleton, Tumor Multiplet, Dual Expanded]: Teff (n = 7522, 276, 735, 5742), Tem (n = 10425, 1002, 1826, 6162), Trm-a (n = 8438, 637, 914, 4470), Trm-b (n = 4484, 286, 959, 2731), and Trm-c (n = 6117, 361, 1215, 3756). The analysis consistently demonstrates elevated PA ratios in the Dual Expansion pattern in Teff and Tem across different $K$ values, suggesting robust identification of potentially tumor-reactive T cells regardless of generation parameter settings.

**a**

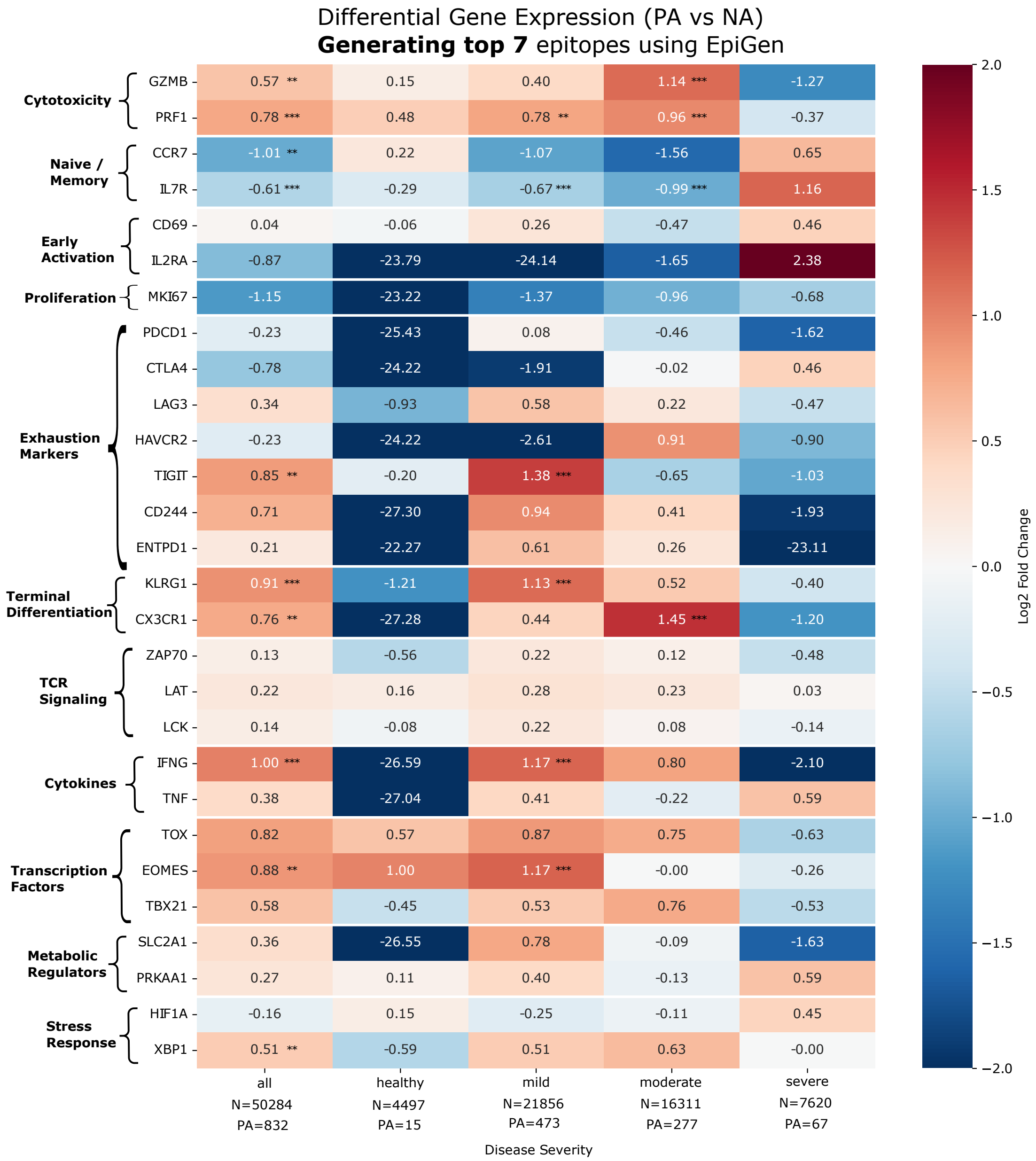
Differential Gene Expression (PA vs NA)
**Generating top 7** epitopes using EpiGen
Cytotoxicity
GZMB 0.57 ** 0.15 0.40 1.14 *** -1.27
PRF1 0.78 *** 0.48 0.78 ** 0.96 *** -0.37
Naive / Memory
CCR7 -1.01 ** 0.22 -1.07 -1.56 0.65
IL7R -0.61 *** -0.29 -0.67 *** -0.99 *** 1.16
Early Activation
CD69 0.04 -0.06 0.26 -0.47 0.46
IL2RA -0.87 -23.79 -24.14 -1.65 2.38
Proliferation
MKI67 -1.15 -23.22 -1.37 -0.96 -0.68
Exhaustion Markers
PDCD1 -0.23 -25.43 0.08 -0.46 -1.62
CTLA4 -0.78 -24.22 -1.91 -0.02 0.46
LAG3 0.34 -0.93 0.58 0.22 -0.47
HAVCR2 -0.23 -24.22 -2.61 0.91 -0.90
TIGIT 0.85 ** -0.20 1.38 *** -0.65 -1.03
CD244 0.71 -27.30 0.94 0.41 -1.93
ENTPD1 0.21 -22.27 0.61 0.26 -23.11
Terminal Differentiation
KLRG1 0.91 *** -1.21 1.13 *** 0.52 -0.40
CX3CR1 0.76 ** -27.28 0.44 1.45 *** -1.20
TCR Signaling
ZAP70 0.13 -0.56 0.22 0.12 -0.48
LAT 0.22 0.16 0.28 0.23 0.03
LCK 0.14 -0.08 0.22 0.08 -0.14
Cytokines
IFNG 1.00 *** -26.59 1.17 *** 0.80 -2.10
TNF 0.38 -27.04 0.41 -0.22 0.59
Transcription Factors
TOX 0.82 0.57 0.87 0.75 -0.63
EOMES 0.88 ** 1.00 1.17 *** -0.00 -0.26
TBX21 0.58 -0.45 0.53 0.76 -0.53
Metabolic Regulators
SLC2A1 0.36 -26.55 0.78 -0.09 -1.63
PRKAA1 0.27 0.11 0.40 -0.13 0.59
Stress Response
HIF1A -0.16 0.15 -0.25 -0.11 0.45
XBP1 0.51 ** -0.59 0.51 0.63 -0.00
all N=50284 PA=832
healthy N=4497 PA=15
mild N=21856 PA=473
moderate N=16311 PA=277
severe N=7620 PA=67
Disease Severity
Log2 Fold Change
2.0
1.5
1.0
0.5
0.0
−0.5
−1.0
−1.5
−2.0

**b**

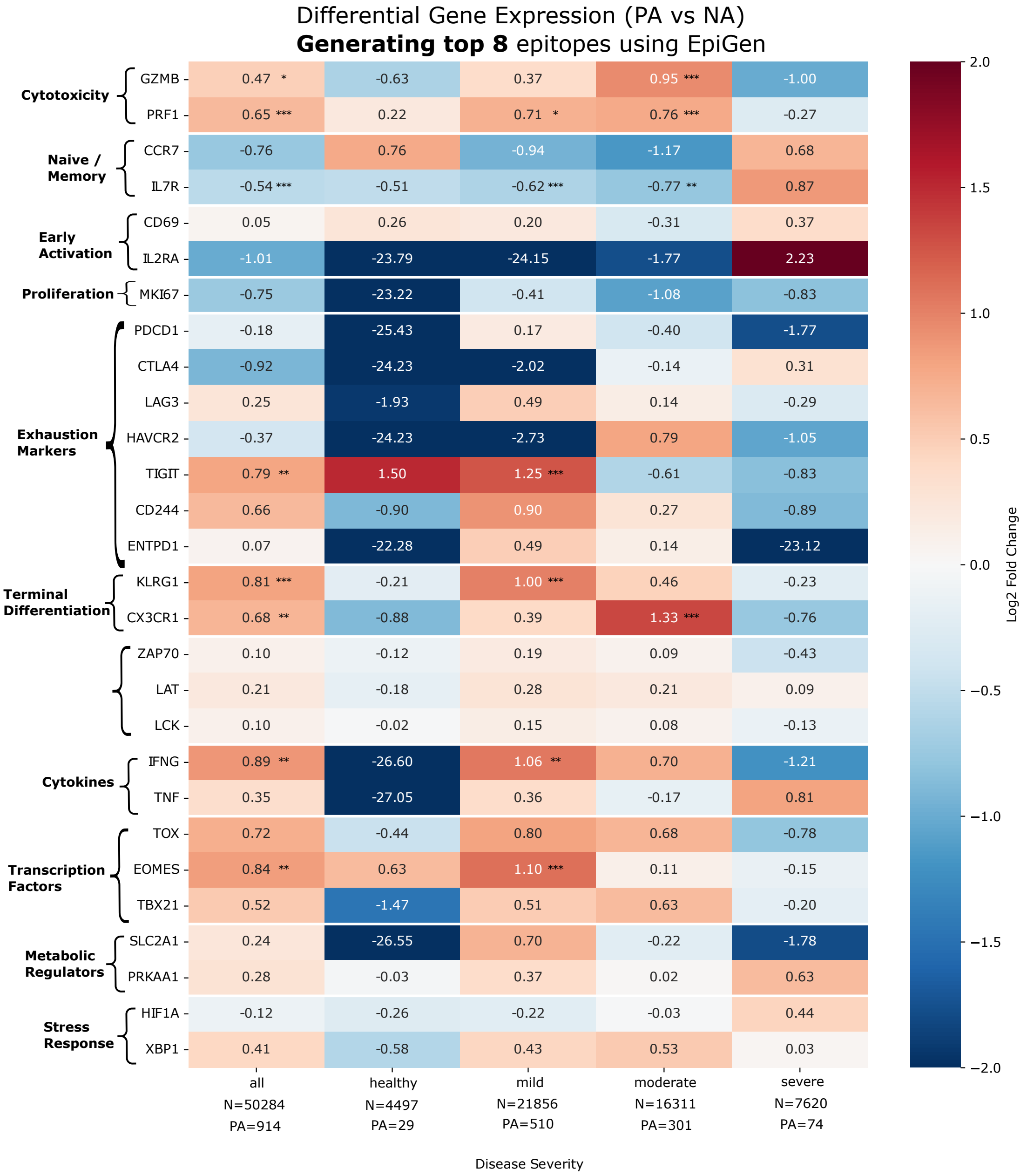
Differential Gene Expression (PA vs NA)
**Generating top 8** epitopes using EpiGen
Cytotoxicity
GZMB 0.47 * -0.63 0.37 0.95 *** -1.00
PRF1 0.65 *** 0.22 0.71 * 0.76 *** -0.27
Naive / Memory
CCR7 -0.76 0.76 -0.94 -1.17 0.68
IL7R -0.54 *** -0.51 -0.62 *** -0.77 ** 0.87
Early Activation
CD69 0.05 0.26 0.20 -0.31 0.37
IL2RA -1.01 -23.79 -24.15 -1.77 2.23
Proliferation
MKI67 -0.75 -23.22 -0.41 -1.08 -0.83
Exhaustion Markers
PDCD1 -0.18 -25.43 0.17 -0.40 -1.77
CTLA4 -0.92 -24.23 -2.02 -0.14 0.31
LAG3 0.25 -1.93 0.49 0.14 -0.29
HAVCR2 -0.37 -24.23 -2.73 0.79 -1.05
TIGIT 0.79 ** 1.50 1.25 *** -0.61 -0.83
CD244 0.66 -0.90 0.90 0.27 -0.89
ENTPD1 0.07 -22.28 0.49 0.14 -23.12
Terminal Differentiation
KLRG1 0.81 *** -0.21 1.00 *** 0.46 -0.23
CX3CR1 0.68 ** -0.88 0.39 1.33 *** -0.76
ZAP70 0.10 -0.12 0.19 0.09 -0.43
LAT 0.21 -0.18 0.28 0.21 0.09
LCK 0.10 -0.02 0.15 0.08 -0.13
Cytokines
IFNG 0.89 ** -26.60 1.06 ** 0.70 -1.21
TNF 0.35 -27.05 0.36 -0.17 0.81
Transcription Factors
TOX 0.72 -0.44 0.80 0.68 -0.78
EOMES 0.84 ** 0.63 1.10 *** 0.11 -0.15
TBX21 0.52 -1.47 0.51 0.63 -0.20
Metabolic Regulators
SLC2A1 0.24 -26.55 0.70 -0.22 -1.78
PRKAA1 0.28 -0.03 0.37 0.02 0.63
Stress Response
HIF1A -0.12 -0.26 -0.22 -0.03 0.44
XBP1 0.41 -0.58 0.43 0.53 0.03
all N=50284 PA=914
healthy N=4497 PA=29
mild N=21856 PA=510
moderate N=16311 PA=301
severe N=7620 PA=74
Disease Severity
Log2 Fold Change
2.0
1.5
1.0
0.5
0.0
−0.5
−1.0
−1.5
−2.0

**c**

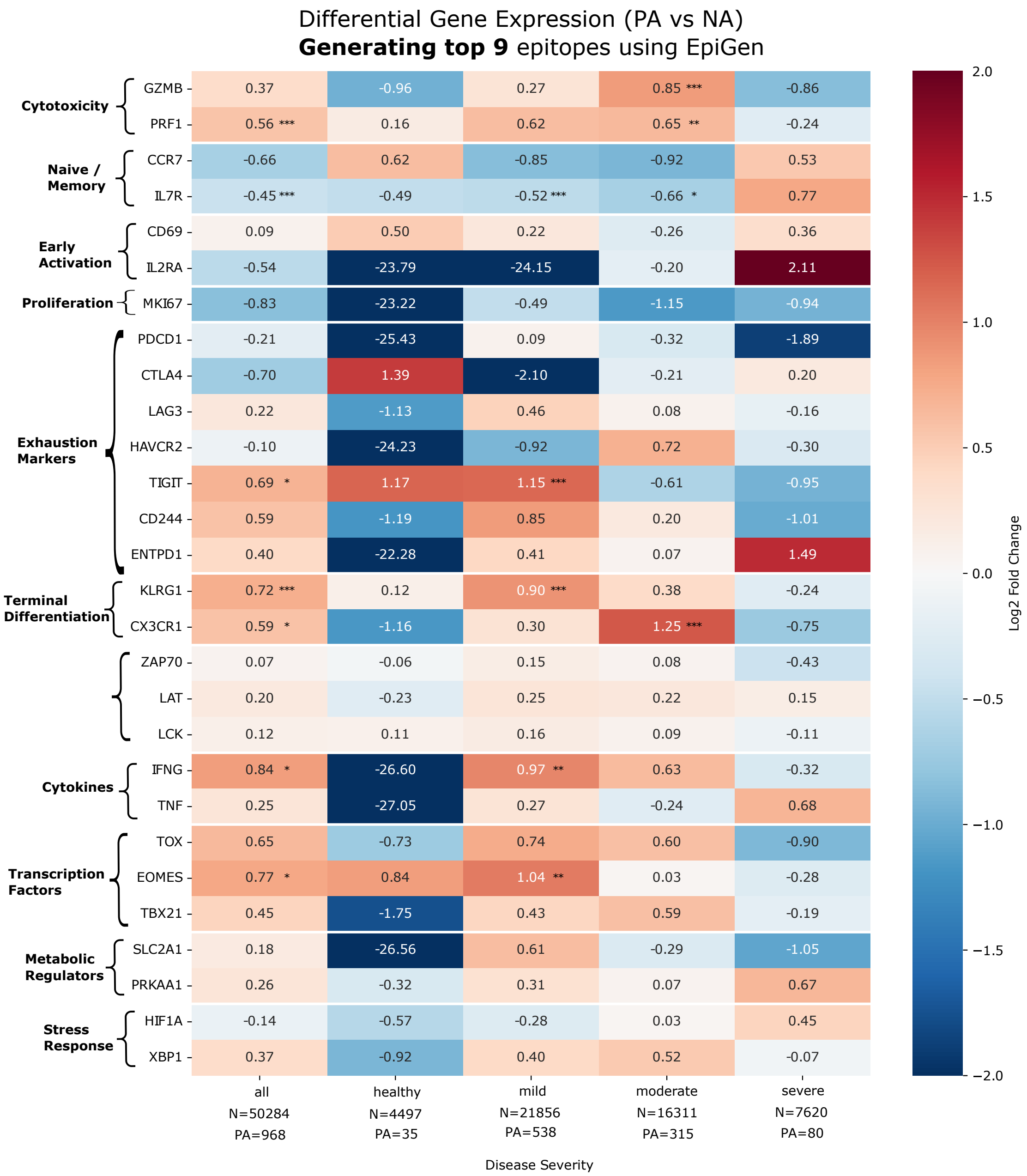

Differential Gene Expression (PA vs NA)
**Generating top 9** epitopes using EpiGen
Cytotoxicity
GZMB 0.37 -0.96 0.27 0.85 *** -0.86
PRF1 0.56 *** 0.16 0.62 0.65 ** -0.24
Naive / Memory
CCR7 -0.66 0.62 -0.85 -0.92 0.53
IL7R -0.45 *** -0.49 -0.52 *** -0.66 * 0.77
Early Activation
CD69 0.09 0.50 0.22 -0.26 0.36
IL2RA -0.54 -23.79 -24.15 -0.20 2.11
Proliferation
MKI67 -0.83 -23.22 -0.49 -1.15 -0.94
Exhaustion Markers
PDCD1 -0.21 -25.43 0.09 -0.32 -1.89
CTLA4 -0.70 1.39 -2.10 -0.21 0.20
LAG3 0.22 -1.13 0.46 0.08 -0.16
HAVCR2 -0.10 -24.23 -0.92 0.72 -0.30
TIGIT 0.69 * 1.17 1.15 *** -0.61 -0.95
CD244 0.59 -1.19 0.85 0.20 -1.01
ENTPD1 0.40 -22.28 0.41 0.07 1.49
Terminal Differentiation
KLRG1 0.72 *** 0.12 0.90 *** 0.38 -0.24
CX3CR1 0.59 * -1.16 0.30 1.25 *** -0.75
ZAP70 0.07 -0.06 0.15 0.08 -0.43
LAT 0.20 -0.23 0.25 0.22 0.15
LCK 0.12 0.11 0.16 0.09 -0.11
Cytokines
IFNG 0.84 * -26.60 0.97 ** 0.63 -0.32
TNF 0.25 -27.05 0.27 -0.24 0.68
Transcription Factors
TOX 0.65 -0.73 0.74 0.60 -0.90
EOMES 0.77 * 0.84 1.04 ** 0.03 -0.28
TBX21 0.45 -1.75 0.43 0.59 -0.19
Metabolic Regulators
SLC2A1 0.18 -26.56 0.61 -0.29 -1.05
PRKAA1 0.26 -0.32 0.31 0.07 0.67
Stress Response
HIF1A -0.14 -0.57 -0.28 0.03 0.45
XBP1 0.37 -0.92 0.40 0.52 -0.07
all N=50284 PA=968
healthy N=4497 PA=35
mild N=21856 PA=538
moderate N=16311 PA=315
severe N=7620 PA=80
Disease Severity
Log2 Fold Change
2.0
1.5
1.0
0.5
0.0
−0.5
−1.0
−1.5
−2.0

**d**

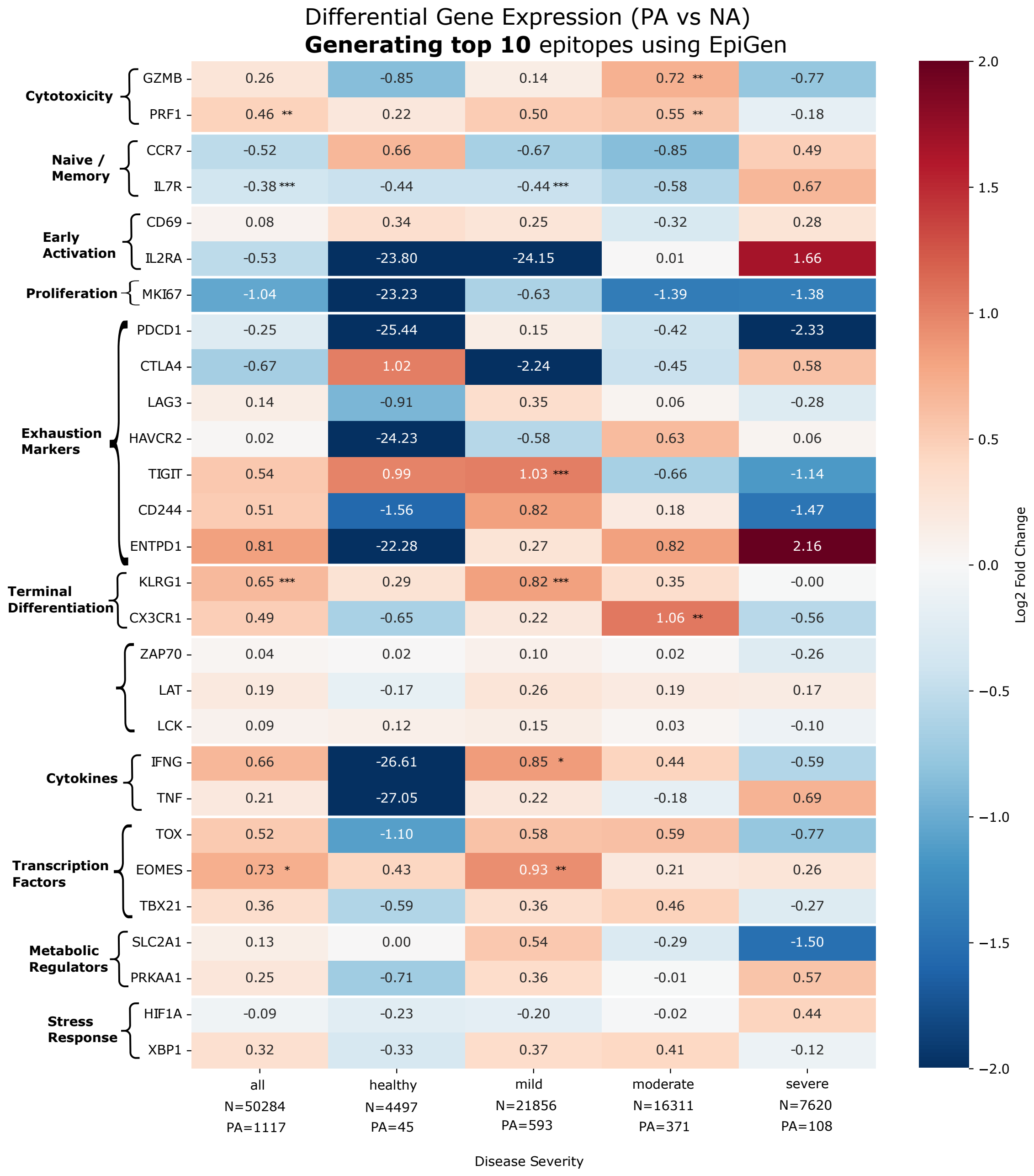

Differential Gene Expression (PA vs NA)
**Generating top 10** epitopes using EpiGen
Cytotoxicity
GZMB 0.26 -0.85 0.14 0.72 ** -0.77
PRF1 0.46 ** 0.22 0.50 0.55 ** -0.18
Naive / Memory
CCR7 -0.52 0.66 -0.67 -0.85 0.49
IL7R -0.38 *** -0.44 -0.44 *** -0.58 0.67
Early Activation
CD69 0.08 0.34 0.25 -0.32 0.28
IL2RA -0.53 -23.80 -24.15 0.01 1.66
Proliferation
MKI67 -1.04 -23.23 -0.63 -1.39 -1.38
Exhaustion Markers
PDCD1 -0.25 -25.44 0.15 -0.42 -2.33
CTLA4 -0.67 1.02 -2.24 -0.45 0.58
LAG3 0.14 -0.91 0.35 0.06 -0.28
HAVCR2 0.02 -24.23 -0.58 0.63 0.06
TIGIT 0.54 0.99 1.03 *** -0.66 -1.14
CD244 0.51 -1.56 0.82 0.18 -1.47
ENTPD1 0.81 -22.28 0.27 0.82 2.16
Terminal Differentiation
KLRG1 0.65 *** 0.29 0.82 *** 0.35 -0.00
CX3CR1 0.49 -0.65 0.22 1.06 ** -0.56
ZAP70 0.04 0.02 0.10 0.02 -0.26
LAT 0.19 -0.17 0.26 0.19 0.17
LCK 0.09 0.12 0.15 0.03 -0.10
Cytokines
IFNG 0.66 -26.61 0.85 * 0.44 -0.59
TNF 0.21 -27.05 0.22 -0.18 0.69
Transcription Factors
TOX 0.52 -1.10 0.58 0.59 -0.77
EOMES 0.73 * 0.43 0.93 ** 0.21 0.26
TBX21 0.36 -0.59 0.36 0.46 -0.27
Metabolic Regulators
SLC2A1 0.13 0.00 0.54 -0.29 -1.50
PRKAA1 0.25 -0.71 0.36 -0.01 0.57
Stress Response
HIF1A -0.09 -0.23 -0.20 -0.02 0.44
XBP1 0.32 -0.33 0.37 0.41 -0.12
all N=50284 PA=1117
healthy N=4497 PA=45
mild N=21856 PA=593
moderate N=16311 PA=371
severe N=7620 PA=108
Disease Severity
Log2 Fold Change
2.0 1.5 1.0 0.5 0.0 −0.5 −1.0 −1.5 −2.0

**Supplementary Fig. 5**: **Impact of epitope generation parameters on differential gene expression patterns across COVID-19 disease severity. a-d.** Analysis of transcriptional differences between Phenotype-Associated (PA) and Not-Associated (NA) T cells across disease severity groups (all, healthy, mild, moderate, and severe) from the Su et al. COVID-19 dataset. Differential expression analysis was performed between PA and NA groups using the two-sided Wilcoxon rank-sum test implemented in Scanpy, and p-values were corrected for multiple testing using the Benjamini-Hochberg method. Four values of epitope generation values ($K$ = 7, 8, 9, and 10) were considered, leveraging EpitopeGen's capacity to generate multiple putative epitope sequences per TCR. The log2 fold changes ($y$-axis) demonstrate consistent transcriptional patterns across different $K$ values, suggesting robust identification of disease-relevant T cell populations regardless of generation parameters. This sensitivity analysis validates the stability of our differential expression findings across varying model parameters.

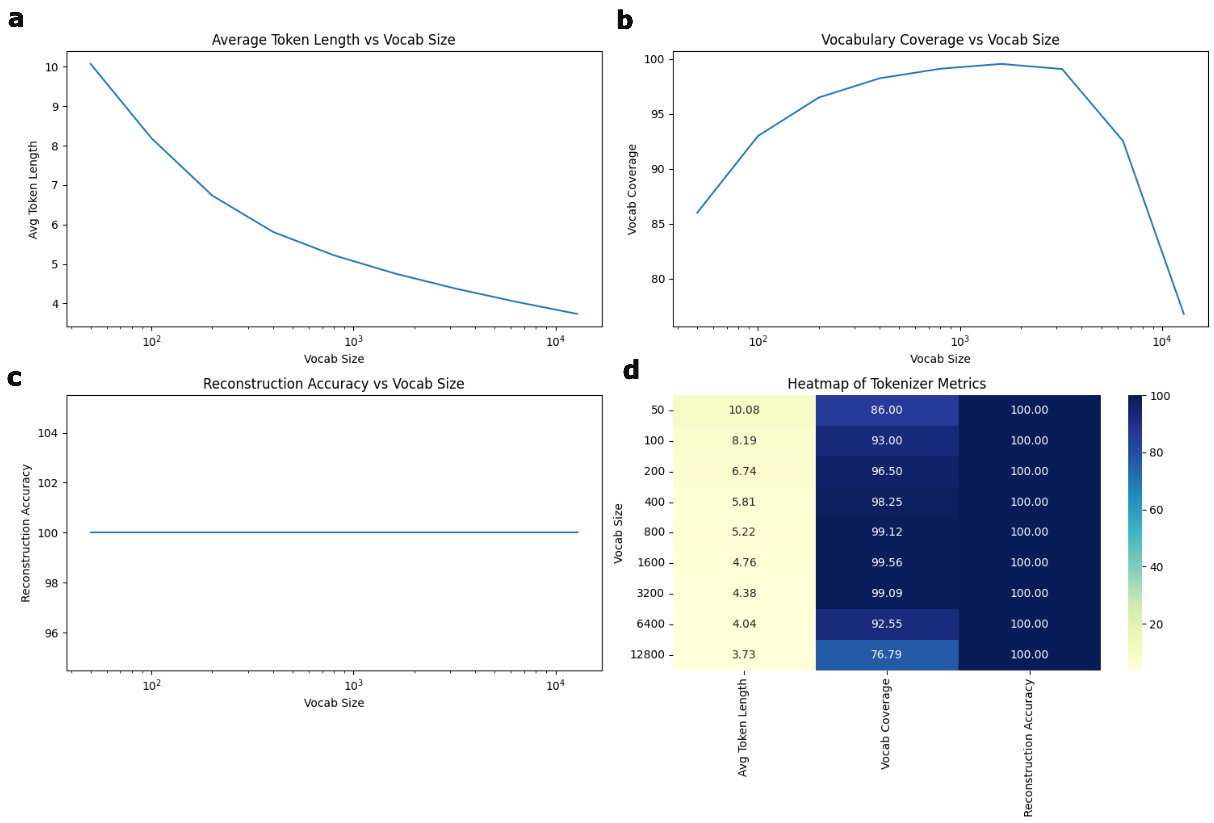

**Supplementary Fig.** 6: **Training of tokenizer using TCR and epitope sequences.** Evaluation metrics across varying vocabulary sizes. **a.** Average token length versus vocabulary size, showing the relationship between sequence tokenization granularity and vocabulary capacity (vocabulary sizes doubled incrementally from 50 to 12,800). **b.** Vocabulary coverage as a function of vocabulary size, demonstrates the proportion of sequences that can be successfully tokenized. **c.** Reconstruction accuracy versus vocabulary size, illustrating the model's ability to faithfully reconstruct original sequences. **d.** Composite heatmap visualization integrates all three metrics (average token length, vocabulary coverage, and reconstruction accuracy) across the range of vocabulary sizes, enabling simultaneous assessment of these key performance indicators.

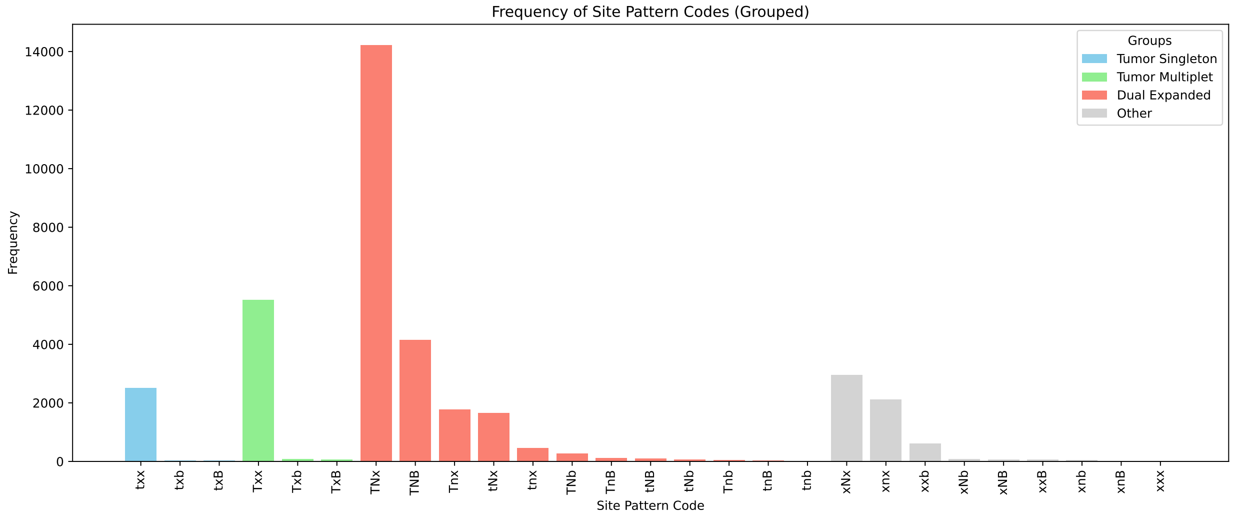

**Supplementary Fig.** 7: **Distribution of site pattern codes in the Wu et al. dataset.** Distribution of T cells across three-digit site pattern codes derived from the Wu et al. dataset. These codes represent distinct T cell localization and expansion patterns, which were subsequently consolidated into four major site pattern categories: All (combining all patterns), Tumor Singleton (observed once in tumor), Tumor Multiplet (observed more than once in tumor), and Dual Expanded (observed in both tumor and normal adjacent tissue). The histogram shows the relative frequency of each of the 26 unique site pattern codes, providing insight into the spatial distribution and clonal expansion patterns of T cells within the tumor microenvironment.

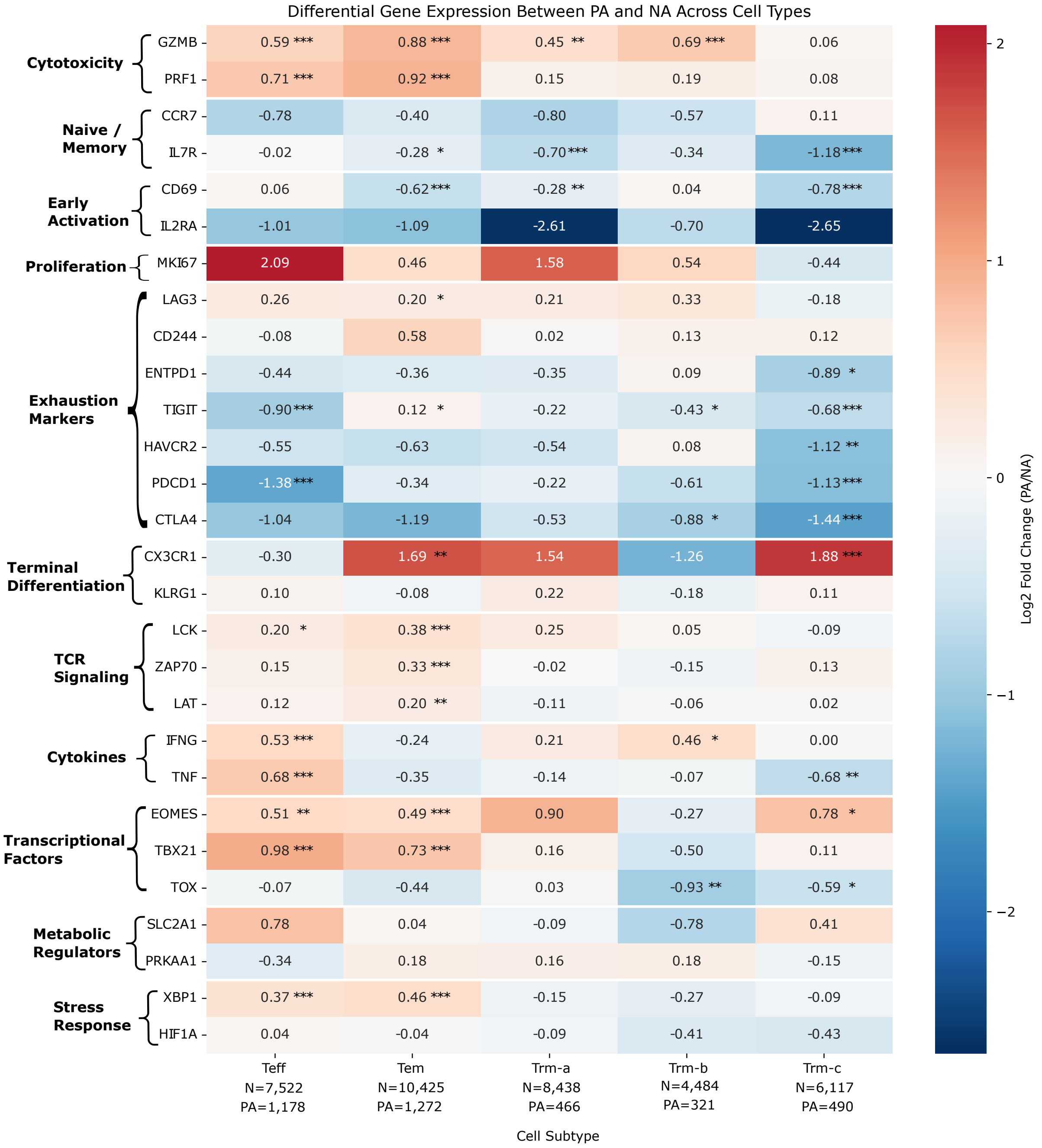

**Supplementary Fig. 8: Cell subtype-specific transcriptional differences between Phenotype-Associated and Not-Associated T cells in cancer.** Comprehensive analysis of differential gene expression between Phenotype-Associated (PA) and Not-Associated (NA) T cells across five distinct $CD8^+$ T cell subtypes (Teff: effector, Tem: effector memory, Trm-a/b/c: tissue-resident memory) in Wu et al. cancer dataset. The heatmap displays the log2 fold changes for curated genes associated with T cell activation states, differentiation, and functional characteristics. Statistical test was performed using the two-sided Wilcoxon rank-sum test implemented in Scanpy where p-values were corrected for multiple testing using the Benjamini-Hochberg method. The significant differences were highlighted by asterisks (*$p < 0.05$, **$p < 0.01$, ***$p < 0.001$).

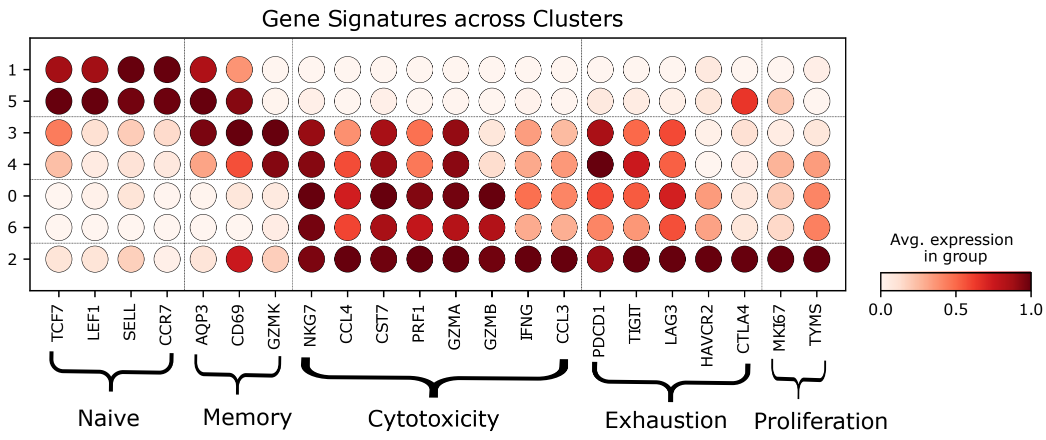

**Supplementary Fig.** 9: **Cluster-specific expression patterns of T cell state-defining gene signatures in COVID-19.** Systematic characterization of T cell populations by computing the mean expression of signature genes across six distinct clusters identified through dimensionality reduction and unsupervised clustering. The dot plot encompasses five key T cell functional programs: (1) Naive state markers (*TCF7*, *LEF1*, *SELL*, *CCR7*) (2) Cytotoxic effector molecules (*NKG7*, *CCL4*, *CST7*, *PRF1*, *GZMA*, *GZMB*, *IFNG*, *CCL3*) (3) Exhaustion markers (*PDCD1*, *TIGIT*, *LAG3*, *HAVCR2*, *CTLA4*) (4) Proliferation indicators (*MKI67*, *TYMS*) (5) Memory-associated genes (*AQP3*, *CD69*, *GZMK*). Color intensity represents mean expression levels, enabling identification and annotation of distinct T cell states in the context of COVID-19 infection.

#### References

1. Shugay, M. *et al.* VDJdb: a curated database of t-cell receptor sequences with known antigen specificity. *Nucleic Acids Res* **46**, D419–D427 (2018).
2. Zhang, W. *et al.* PIRD: Pan immune repertoire database. *Bioinformatics* **36**, 897–903 (2019).
3. Vita, R. *et al.* The immune epitope database (iedb): 2018 update. *Nucleic Acids Research* **47** (2019).
4. Tickotsky, N., Sagiv, T., Prilusky, J., Shifrut, E. & Friedman, N. McPAS-TCR: a manually curated catalogue of pathology-associated T cell receptor sequences. *Bioinformatics* **33**, 2924–2929 (2017).
5. Chen, S.-Y., Yue, T., Lei, Q. & Guo, A.-Y. TCRdb: a comprehensive database for t-cell receptor sequences with powerful search function. *Nucleic Acids Res* **49**, D468–D474 (2021).
6. Jurtz, V. *et al.* NetMHCpan-4.0: Improved Peptide-MHC class I interaction predictions integrating eluted ligand and peptide binding affinity data. *J Immunol* **199**, 3360–3368 (2017).
7. O'Donnell, T. J., Rubinsteyn, A. & Laserson, U. Mhcflurry 2.0: Improved pan-allele prediction of mhc class i-presented peptides by incorporating antigen processing. *Cell Systems* **11**, 42–48.e7 (2020).
8. Huang, X. *et al.* The SysteMHC Atlas v2.0, an updated resource for mass spectrometry-based immunopeptidomics. *Nucleic Acids Research* **52**, D1062–D1071 (2023).
9. Glanville, J. *et al.* Identifying specificity groups in the T cell receptor repertoire. *Nature* **547**, 94–98 (2017).
10. Nolan, S. *et al.* A large-scale database of t-cell receptor beta (TCR$\beta$) sequences and binding associations from natural and synthetic exposure to SARS-CoV-2 (2020).
11. 10x Genomics. A new way of exploring immunity: Linking highly multiplexed antigen recognition to immune repertoire and phenotype. Application Note, 10x Genomics (2022).
12. Klenerman, P., Cerundolo, V. & Dunbar, P. R. Tracking T cells with tetramers: new tales from new tools. *Nature Reviews Immunology* **2**, 263–272 (2002).
13. Masopust David, Murali-Krishna Kaja & Ahmed Rafi. Quantitating the magnitude of the lymphocytic choriomeningitis Virus-Specific CD8 T-Cell response: It is even bigger than we thought. *Journal of Virology* **81**, 2002–2011 (2007).

14. Moutaftsi, M. *et al.* A consensus epitope prediction approach identifies the breadth of murine TCD8+-cell responses to vaccinia virus. *Nature Biotechnology* **24**, 817–819 (2006).

15. Addo, M. M. *et al.* Comprehensive epitope analysis of human immunodeficiency virus type 1 (HIV-1)-specific t-cell responses directed against the entire expressed HIV-1 genome demonstrate broadly directed responses, but no correlation to viral load. *J Virol* **77**, 2081–2092 (2003).

16. Shepherd, F. R. & McLaren, J. E. T cell immunity to bacterial pathogens: Mechanisms of immune control and bacterial evasion. *Int J Mol Sci* **21** (2020).

17. Friot, A. *et al.* Antigen specific activation of cytotoxic CD8(+) T cells by staphylococcus aureus infected dendritic cells. *Front Cell Infect Microbiol* **13**, 1245299 (2023).

18. Kenison, J. E., Stevens, N. A. & Quintana, F. J. Therapeutic induction of antigen-specific immune tolerance. *Nature Reviews Immunology* **24**, 338–357 (2024).

19. Rizzuto, G. A. *et al.* Self-antigen-specific CD8+ T cell precursor frequency determines the quality of the antitumor immune response. *J Exp Med* **206**, 849–866 (2009).

20. Nelson, C. E. *et al.* Robust iterative stimulation with Self-Antigens overcomes CD8(+) T cell tolerance to self- and tumor antigens. *Cell Rep* **28**, 3092–3104.e5 (2019).

21. Pittet, M. J. *et al.* High frequencies of naive Melan-A/MART-1-specific CD8(+) T cells in a large proportion of human histocompatibility leukocyte antigen (HLA)-A2 individuals. *J Exp Med* **190**, 705–715 (1999).

22. Mittal, J., Ponce, M. G., Gendlina, I. & Nosanchuk, J. D. Histoplasma capsulatum: Mechanisms for pathogenesis. *Curr Top Microbiol Immunol* **422**, 157–191 (2019).

23. Stuckey Peter V. & Santiago-Tirado Felipe H. Fungal mechanisms of intracellular survival: what can we learn from bacterial pathogens? *Infection and Immunity* **91**, e00434–22 (2023).

24. Cavicchioli, R., Curmi, P. M., Saunders, N. & Thomas, T. Pathogenic archaea: do they exist? *BioEssays* **25**, 1119–1128 (2003).

25. Gill, E. E. & Brinkman, F. S. L. The proportional lack of archaeal pathogens: Do viruses/phages hold the key? *BioEssays* **33**, 248–254 (2011).

26. Camacho, C. *et al.* BLAST+: architecture and applications. *BMC Bioinformatics* **10**, 421 (2009).

27. NCBI Resource Coordinators. Database resources of the national center for biotechnology information. *Nucleic Acids Res* **46**, D8–D13 (2018).

28. Yin, R. *et al.* TCRmodel2: high-resolution modeling of T cell receptor recognition using deep learning. *Nucleic Acids Res* **51**, W569–W576 (2023).

29. Jumper, J. *et al.* Highly accurate protein structure prediction with AlphaFold. *Nature* **596**, 583–589 (2021).

30. Stranges, P. B. & Kuhlman, B. A comparison of successful and failed protein interface designs highlights the challenges of designing buried hydrogen bonds. *Protein Sci* **22**, 74–82 (2012).

31. Wu, T. D. *et al.* Peripheral T cell expansion predicts tumour infiltration and clinical response. *Nature* **579**, 274–278 (2020).

32. Lu, T. *et al.* Deep learning-based prediction of the T cell receptor–antigen binding specificity. *Nature Machine Intelligence* **3**, 864–875 (2021).

33. Zhang, J., Ma, W. & Yao, H. Accurate TCR-pMHC interaction prediction using a BERT-based transfer learning method. *Briefings in Bioinformatics* **25**, bbad436 (2023).

34. Sennrich, R., Haddow, B. & Birch, A. Neural machine translation of rare words with subword units. In Erk, K. & Smith, N. A. (eds.) *Proceedings of the 54th Annual Meeting of the Association for Computational Linguistics (Volume 1: Long Papers)*, 1715–1725 (Association for Computational Linguistics, Berlin, Germany, 2016).

35. Wolf, T. *et al.* Transformers: State-of-the-art natural language processing. In Liu, Q. & Schlangen, D. (eds.) *Proceedings of the 2020 Conference on Empirical Methods in Natural Language Processing: System Demonstrations*, 38–45 (Association for Computational Linguistics, Online, 2020).

36. Charoentong, P. *et al.* Pan-cancer immunogenomic analyses reveal genotype-immunophenotype relationships and predictors of response to checkpoint blockade. *Cell Reports* **18**, 248–262 (2017).
